# The link between space and time along the human cortical hierarchy

**DOI:** 10.1101/2024.04.15.589551

**Authors:** Valeria Centanino, Gianfranco Fortunato, Domenica Bueti

**Affiliations:** International School for Advanced Studies (SISSA), Trieste, Italy

**Author notes:** These authors contributed equally to this work (Valeria Centanino, Gianfranco Fortunato).

## Abstract

In humans, very few studies have directly tested the link between the neural coding of time and space. Here we combined ultra-high field functional magnetic resonance imaging with neuronal-based modeling to investigate how and where the processing and the representation of a visual stimulus duration is linked to that of its spatial location. Results show a transition in the neural response to duration: from monotonic and spatially-dependent in early visual cortex, to unimodal and spatially-invariant in frontal cortex. This transition begins in extrastriate areas V3AB, and it fully displays in the intraparietal sulcus (IPS), where both unimodal and monotonic responses are present and where neuronal populations are selective to either space, time or both. In IPS, space and time topographies show a specific relationship, although along the cortical hierarchy duration maps compared to spatial ones are smaller in size, less clustered and more variable across participants. These results help to identify the mechanisms through which humans perceive the duration of a visual object with a specific spatial location and precisely characterize the functional link between time and space processing, highlighting the importance of space-time interactions in shaping brain responses.

## Introduction

A key ingredient for a unitary perceptual appraisal of the external environment is the ability to combine the spatial and temporal information of the sensory inputs. In a warm summer night, for instance, we can enjoy the view of a swarm of fireflies since we perceive the position and the duration of the bioluminescence of each firefly. Despite this tight link between the spatial and the temporal aspects of our sensory experience, not many studies have directly investigated how and where the human brain synergistically links these two types of information. In the visual system for example it is unclear to what extent the duration processing of a visual stimulus entails spatial circuits and follows spatial representational rules, i.e., retinotopy for example. In humans, psychophysical studies suggested the presence of both spatially-specific and spatially-independent mechanisms of duration processing. A few studies have shown that the adaptation to a fast moving stimulus[1]–[6] causes spatially-specific biases in duration perception, i.e., the perceptual bias disappears if the stimulus to be timed is presented in a different spatial position from the adaptor stimulus. On the other hand, it has been shown that the perceptual after-effect caused by duration adaptation transfers across visual hemifields and quadrants[7], [8] suggesting a spatially independent temporal processing. These apparently conflicting evidences suggest that duration encoding and representation might be the outcome of a double-stage processing. One early and spatially dependent, which might be related to the processing of visual information in early visual cortices, and one spatially invariant that might arise later in the visual processing hierarchy[9].

In line with the idea of a double-stage processing of duration information but without a clear spatial connotation, recent Functional Magnetic Resonance Imaging (fMRI) studies suggest two possible mechanisms underlying the temporal processing of visual stimuli. Zhou and colleagues have shown that along the visual stream (from primary visual cortex - V1 to the intraparietal sulcus - IPS) stimulus duration information results from a compressive (i.e., non-linear) summation of the sensory input[10]. Conversely, a series of high-spatial resolution fMRI works suggest that temporal information is supported by unimodal tuning mechanisms entailing a topographical organization[11], [12]. Duration maps (or chronomaps) have been identified in a wide network of brain areas spanning from lateral-occipital to parietal, frontal and premotor regions. Interestingly, chronomaps not only appear in some of the brain regions showing a monotonic encoding of stimulus duration, but they have also been found to partially overlap with retinotopic maps[11]. In addition, a recent reanalysis of the original findings of Harvey et al. 2020[13] has shown a transition between monotonic and unimodal responses to stimulus duration along the cortical hierarchy and a switch between these two types of response in area V5/MT. Although these previous fMRI studies are crucial for understanding the mechanism underlying duration processing, they overlooked the spatial dimension of the stimuli in both the experimental design and modeling approaches. As a result, they could not establish a direct link between the representation of spatial and temporal information. This omission prevents these studies from reconciling the behavioral evidence described above and from uncovering common organizational principles of spatial and temporal information processing.

In this study we therefore sought to understand the extent to which the processing of time is linked to that of space along the visual hierarchy, whether this link entails monotonic or unimodal responses to stimulus duration and whether there is a relationship between spatial and duration topographies. To address these issues, we asked human participants to judge the duration of a small visual stimulus varying in both duration and spatial position within foveal and parafoveal ranges and we measured brain activity with high-spatial resolution fMRI (7 Tesla). The simultaneous manipulation of stimulus’s duration and position allowed us also to understand how and where, along the dorsal visual stream, brain responses change as function of the combination of stimulus’s duration and position, and to directly identify the neuronal populations selective to either or both stimulus features (i.e., duration and spatial position).

## Results

In this study we used a single interval duration discrimination task where participants (13 healthy volunteers) were asked to compare the duration of a visual stimulus (i.e., comparison stimulus) varying at each trial in both duration (i.e., display time) and spatial position (i.e., display location on the screen) to a previously internalized reference duration. The task was to report whether the comparison stimulus (ranging from 0.2 to 0.8 s) was longer or shorter than the reference stimulus (0.5 s, see figure 1a). The comparison stimulus could be presented, with respect to a fixation cross, at either 2.5*^◦^* or 0.9*^◦^* of visual angle in the lower left or lower right quadrant of the visual field. Comparison durations varied pseudo-randomly across trials, spatial positions varied sequentially (see *Task and experimental design* for more details). This modality of stimulus’s presentation ensured participants engagement in the duration discrimination task while minimizing possible biases due to changes in spatial attention[14]. Figure 1a shows a pictorial representation of the trial structure, the close-up highlights the four spatial positions of the stimulus. Each participant performed 10 blocks (48 trials each, 2 trials per combination of stimulus duration and position) acquired in separate fMRI runs. Participants also underwent two retinotopy runs which allowed us to precisely estimate individual visual field maps with an independent dataset (see *Retinotopy runs* and *Retinotopic mapping*).

**Figure 1:**
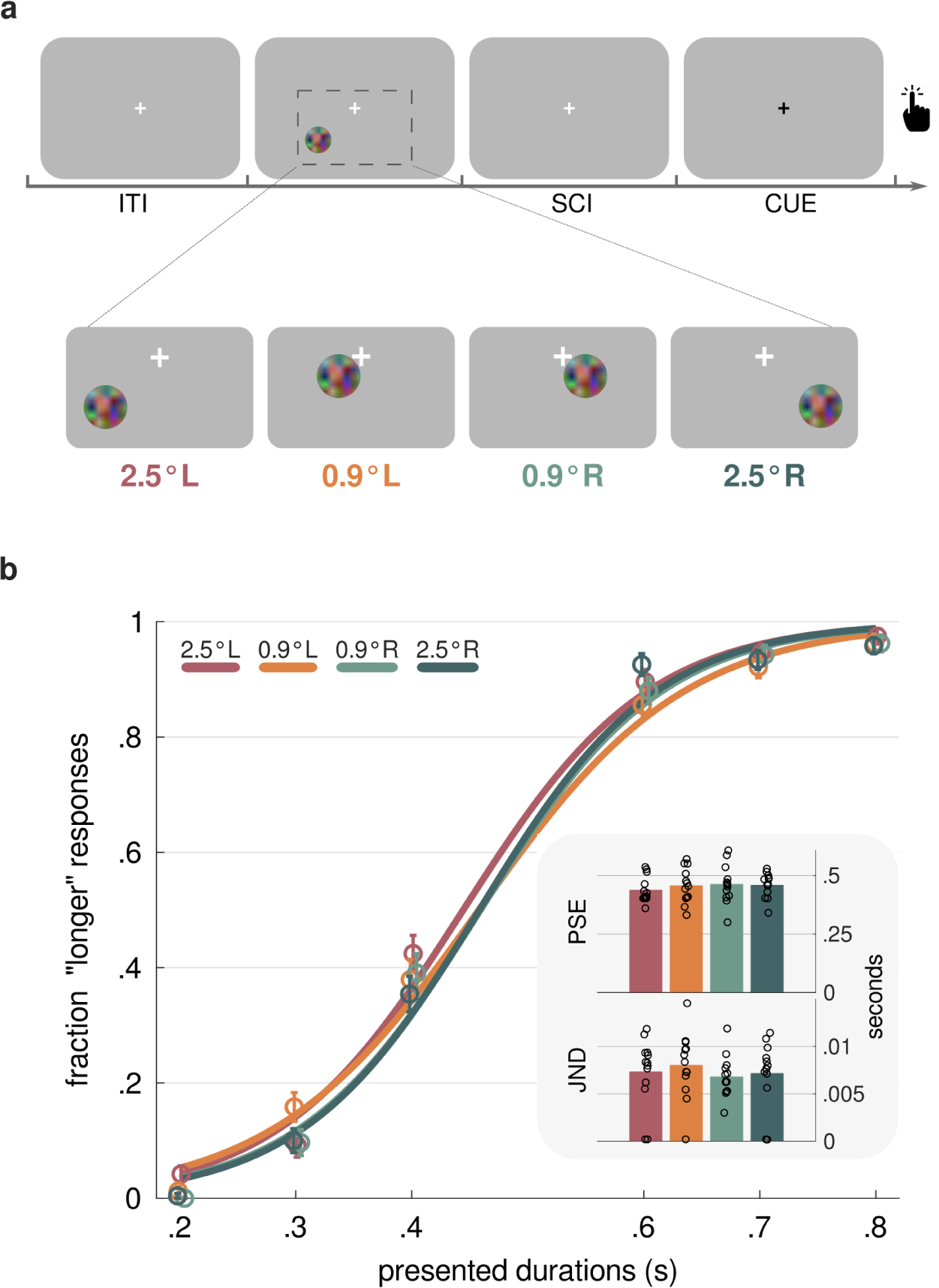
Experimental procedure and behavioral results. **(a)** In each trial, one of six different comparison durations (i.e., 0.2, 0.3, 0.4, 0.6, 0.7, 0.8 s) was displayed at a either 2.5*^◦^* or 0.9*^◦^* of visual angle in the lower-left (L) or lower-right (R) quadrant of the visual field. Durations varied trial-by-trial in a pseudo-random fashion, whereas positions varied sequentially, from 2.5*^◦^* L to 2.5*^◦^* R and backwards, as shown in the close-up. Participants were asked to compare the duration of the comparison stimulus with that of an internalized reference and to report with a key press which one was longer. After a randomized interval from the offset of the comparison (stimulus-cue interval - SCI, uniformly drawn between 0.9-1.2 s), the response was cued with a color switch of the fixation cross from white to black. Trials were interleaved by a uniformly distributed inter-trial interval (ITI) spanning from 1.8 to 2.5 s. The fixation cross was displayed at the center of the screen throughout the experiment (see *Stimuli and Experimental Procedure*). **(b)** Group psychometric curves are shown color-coded according to the spatial position of the comparison stimulus. Circles represent the average fraction of “comparison longer than reference” responses across participants for each comparison duration, and error bars represent standard errors. In the inset, bar plots represent the average PSE (top) and JND (bottom) values across participants for each spatial position of the stimulus. The color code is as in (a). Black circles represent individual PSE and JND values (see *Behavioral performance analysis*).

### The spatial position of the stimulus does not affect its duration discrimination

We analyzed the behavioral data to make sure participants accurately performed the duration discrimination task and duration judgements were not affected by the display location of the comparison stimulus. For each spatial position of the comparison stimulus, we estimated psychometric curves on the mean fraction of “comparison longer than reference” responses for each comparison duration (see *Behavioral performance analysis*). In addition, for each spatial position of the comparison stimulus we also derived individual Point of Subjective Equality (PSE) and Just Noticeable Difference (JND) values. Figure 1b shows group psychometric curves color-coded according to the stimulus position, and both group and individual PSE and JND values are displayed in the inset. To test duration discrimination differences among spatial positions, individual PSE and JND values were analyzed with two linear mixed effect (LME) models (PSE model formula: *PSE ∼ StimulusPosition*+(1*|subjectID*), marginal *R*^2^ = 0.02, conditional *R*^2^ = 0.70; JND model formula: *JND ∼ StimulusPosition* + (1*|subjectID*), marginal *R*^2^ = 0.01, conditional *R*^2^ = 0.61). A type III ANOVA on model estimates revealed no main effect of spatial position for both PSE and JND values (see supplementary tables 1 and 2). These results confirm that the display location of the stimulus did not induce any bias nor changed duration perception sensitivity.

### How brain responses to stimulus’s duration change along the cortical hierarchy

We started our investigations by asking to which extent the duration processing of a visual stimulus entails spatial circuits and follows spatial representational rules. We first performed a general linear model (GLM) analysis on fMRI data (see *General Linear Model (GLM) analysis*), using as events of interest the 24 unique combinations of the 6 durations and 4 positions of the comparison stimulus time-locked to its offset. The GLM beta weights were then used to perform a voxel-wise modeling with the population receptive field (pRF) method (see *Population Receptive Field (pRF) modeling*). To be able to tell if duration modulates brain responses in spatial circuits and if this happens in association or not with spatial responses, we tested four different neuronal response models (pRFs). These pRFs were tailored to capture different tuning properties of the BOLD responses linked to either space, time or to both. The Compressive Monotonic Time (CMT) model assumes a neuronal response scaling monotonically and sub-additively to increasing stimulus’s durations[10], and independently of the stimulus’s spatial position (model highlighted in dark blue in figure 2). The Gaussian Space (GS) and Gaussian Time (GT) models describe two independent unimodal tuning mechanisms for stimulus position and duration[11], [12], [15] only (models highlighted in red and light blue respectively in figure 2). The Compressive Monotonic Time and Space (CMTS) model represents the combination of the GS and the CMT models and assumes instead a spatially-specific neuronal response which scales monotonically and sub-additively to increasing stimulus’s durations (model highlighted in orange in figure 2). See supplementary figure 1 for an example of the result of the fitting procedure for each neural response model.

**Figure 2:**
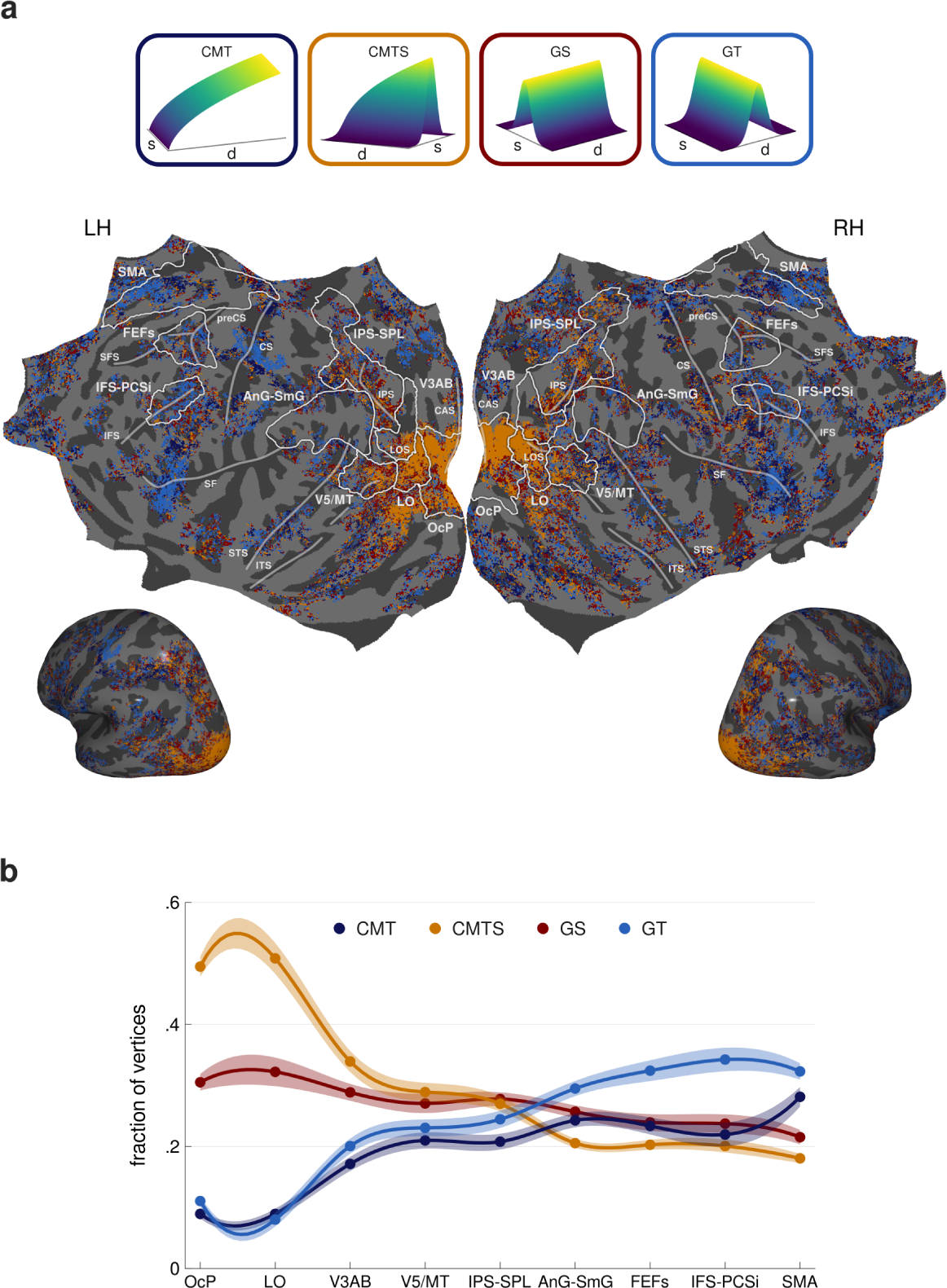
Models’ performance comparison. **(a)** The group-level vertex-wise distribution of winning neural response models is plotted onto a common surface (fsaverage). The distribution was obtained by resampling all individual winning models’ distributions onto fsaverage surface and computing for each vertex the mode across participants. Vertices were excluded if they had non-integer values after the surface resampling, if they showed more than one winning model, or if they were not labeled with a winning model in at least 7 participants out of 13. This distribution is therefore highly conservative, and it was computed for visualization purposes only (see *Neuronal response models comparison*). Each vertex is color-coded according to its winning model as shown in the top panel. Bright white lines outline the 9 bilateral ROIs in which models’ distribution was quantified. Semi-transparent white lines mark principal sulci. **(b)** The group-level fraction of vertices assigned to each model is plotted for each ROI outlined in (a), ordered from occipital to frontal areas. Each dot represents the mean fraction of vertices across participants and hemispheres for each model, color-coded as in (a), and the shaded area is its standard error. For visualization purposes we also plotted lines obtained with a spline interpolation across dots. ROI legend: OcP = occipital pole, LO = lateral occipital, V3AB = visual areas V3A and V3B, V5/MT = visual area V5/MT, AnG-SmG = angular gyrus and supramarginal gyrus, IPS-SPL = intraparietal sulcus and superior parietal lobule, IFS-PCSi = inferior frontal sulcus and precentral sulcus - inferior part, FEFs = frontal eye fields, SMA = supplementary motor area; sulci legend: CAS = calcarine sulcus, LOS = lateral occipital sulcus, ITS = inferior temporal sulcus, STS = superior temporal sulcus, IPS = intraparietal sulcus, SF = Sylvian fissure, CS = central sulcus, IFS = inferior frontal sulcus, SFS = superior frontal sulcus, preCS = precentral sulcus; LH = left hemisphere, RH = right hemisphere.

In each individual subject, we assessed models’ performance by comparing their cross-validated *R*^2^ (see *R*^2^ *cross-validation*). We used a winner-take-all procedure to assign to each vertex of the cortical surface the model with the highest cross-validated *R*^2^ value (i.e., best fitting model). Figure 2a shows the group-level result of this procedure. We then computed for each participant the fraction of vertices assigned to each model in 9 bilateral regions of interest (ROIs). Since in this first analysis we tested pRFs models purely sensitive to time (i.e., GT, CMT), we decided to focus our investigation on a wide set of ROIs, some of which were outside the spatial circuits. The ROIs spanned from the occipital pole to the inferior frontal cortex and included brain regions that, according to previous works, are engaged in either spatial[16] or temporal[12], [17] processing (see *Atlas-based ROIs*). Figure 2b shows the group-level results averaged across hemispheres. To analyze these data, we used a three-way repeated measure ANOVA with model type, brain hemisphere and ROI as factors (see *Neuronal response models comparison*). The ANOVA showed a significant effect of model type (F(3,36) = 24.25, p *<* .0001) and a significant interaction between model type and ROI (F(24,288) = 33.16 p *<* .0001), indicating that brain responses in different ROIs were best captured by different models. In particular, in the occipital pole (OcP) and in the lateral occipital cortex (LO) the CMTS model outperformed the other models (all t(864) *>* 9.26, p *<* .001), suggesting that in those regions stimulus duration is encoded via a monotonic increase in the amplitude of spatially-specific neuronal responses. Interestingly, in V3A and V3B (V3AB), a space-invariant temporal encoding mechanism, represented by the GT and the CMT models, started to appear (for both CMT and GT, all t(864) *>* 4.08 resulting from the comparison between V3AB, OcP and LO, p *<* 0.005), while the explanatory power of the CMTS model decreased (all t(864) *<* −7.74, p *<* 0.001). In intraparietal and superior parietal areas (IPS-SPL) the GT model was represented as much as the models assuming a spatially-specific response (CMTS and GS models), and it prevailed over the CMTS model from the inferior parietal lobule (AnG-SmG) onwards (all t(864) *>* 4.45, p *<* 0.001). The GT model also prevailed over the GS model from the frontal eye fields (FEFs) onwards (all t(864) *>* 4.22, p *<* 0.001). In addition, the two spatially-invariant duration response models (CMT and GT models) were equally represented in V5/MT, IPS-SPL, AnG-SmG, and in the supplementary motor area (SMA). The unimodal response model to durations (GT) prevailed over the monotonic one (CMT) in the FEFs (t(864) = 4.51, p *<* 0.001) and in the inferior frontal sulcus (IFS-PCSi, t(864) = 6.12, p *<* 0.001). These results suggest that along the cortical hierarchy temporal processing gradually detaches from spatial processing and that in downstream areas it is mainly supported by unimodal responses of spatially-invariant neuronal populations.

In addition, the IPS-SPL exhibited the coexistence of neuronal populations with different tuning mechanisms. This last result seems to fit with the role of IPS-SPL as place in the cortical hierarchy where spatial and temporal information are integrated[18]. The details of the ANOVA and of the Bonferroni-corrected multiple comparisons tests are reported in the supplementary tables 3 - 7. In summary, these findings show a transition along the dorsal visual stream of brain responses to stimulus’s duration: from monotonic and spatially-dependent responses in occipital cortex (OcP and LO), to unimodal and spatially-invariant responses in frontal cortex (FEFs, IFS-PCSi, and SMA). This transition started in V3AB and in this area but also in V5/MT and parietal cortex (IPS-SPL, AnG-SmG) the two types of response were either present in different proportions (in V3AB, V5/MT, and AnG-SmG) or equally matched (in IPS-SPL).

### How duration processing interacts with eccentricity processing

The results from the previous section show the coexistence from extrastriate area V3AB to superior and inferior parietal lobule (IPS-SPL, AnG-SmG) of brain responses sensitive to either space and time separately or to space and time together. To explore in-depth the interactions between these stimulus features in eliciting brain responses in these brain regions, we next tested another neuronal response model (Gaussian Space and Time, GST), which assumes unimodal tuning functions for both stimulus position and duration (see *Population Receptive Field (pRF) modeling*). The GST model describes the BOLD response with five parameters: the stimulus’s spatial position and duration eliciting the greatest neuronal response (*µ_s_* and *µ_d_*, respectively), the sensitivity of the response (*σ_s_* and *σ_d_*), and the orientation of the tuning function (*θ*), which appraises the contribution of each stimulus dimension to the neuronal response. An example of the results of the fitting procedure is presented in supplementary figure 2. The GST model allowed us to capture responses described by the previously tested CMT, GS, GT, and CMTS models (see supplementary figure 1), but differently from those models, enabled us to directly estimate the interaction between stimulus’s position and duration in eliciting brain responses. Since our goal was to investigate the relationship between visual duration and eccentricity processing, we decided to focus our analyses on visuo-spatial circuits. We indeed analyzed the parameters of the new GST model within 8 bilateral ROIs that according to individual eccentricity maps, estimated with an independent dataset (i.e., the retinotopic fMRI runs, see *Retinotopy runs* and *Retinotopic mapping* for more details), belonged to the dorsal visual stream. These ROIs were located in occipital cortex (3 ROIs covering dorsal V1 and V2, V3A and V3B, V5/MT), parietal cortex (4 ROIs covering the intraparietal sulcus i.e., IPS0, IPS1, IPS2, IPS3), and frontal cortex (1 ROI covering the FEFs), see *Custom-made ROIs*. Figure 3a illustrates the eccentricity maps of one participant, with the ROIs marked by blue, red, and white lines.

**Figure 3:**
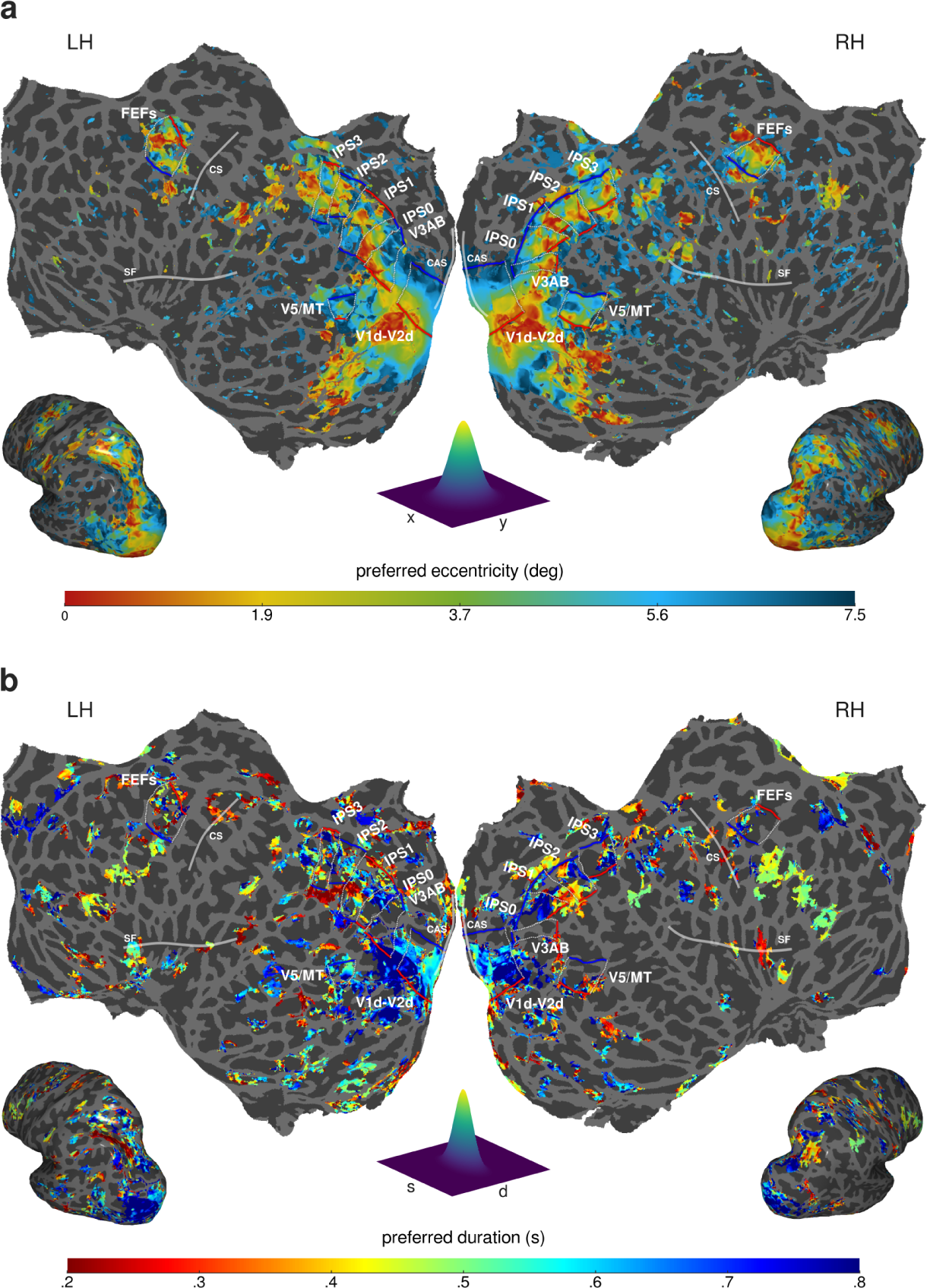
Maps of pRFs preferred eccentricity and duration. The distribution of preferred eccentricity (a) and preferred duration (b) of an example participant are shown projected onto the inflated and flattened native cortical surface. Different eccentricity and duration preferences are color-coded. The eccentricity map was obtained from the pRF modeling of the retinotopy data with a two-dimensional Gaussian function (see *Retinotopic mapping*). The duration map was obtained from the pRF modeling of the experimental data with the GST model (see *Gaussian Space-Time (GST) model*). A set of eight bilateral ROIs was identified based on individual retinotopic maps. Red and blue lines correspond respectively to the low and to the high borders of the eccentricity progression within each ROI. Lateral borders are represented with dashed white lines (see *Custom-made ROIs*). Semi-transparent white lines outline principal sulci. ROI legend: V1-2d = dorsal primary and secondary visual areas, V3AB = visual areas V3Aand V3B, V5/MT = visual area V5/MT, IPS0-3 = different portions of the intraparietal sulcus, FEFs = frontal eye fields. Sulci legend: CAS = calcarine sulcus, SF = Sylvian fissure, CS = central sulcus, LH = left hemisphere, RH = right hemisphere.

To characterize the space-time interactions along the visual hierarchy, in this second set of analyses we studied the changes in the GST model parameters. The first parameter we checked was the *µ_d_*parameter, which represents the duration preference. We focused on *µ_d_* in the first place to check the presence of duration unimodal tuning within eccentricity-defined ROIs. Figure 3b illustrates the cortical distribution of *µ_d_* in the two hemispheres of one participant, whereas figure 4a shows the group-level distribution of *µ_d_* within each ROI. To assess whether duration preferences changed across ROIs, we used a LME model (see *pRFs duration preference*) with ROI and hemisphere as factors and subjects as random intercept (model formula: *µ_d_ ∼ ROI ∗Hemisphere*+(1*|subjectID*), marginal *R*^2^: 0.22, conditional *R*^2^: 0.30). Type III ANOVA on model estimates showed a main effect of ROI (F(7,180) = 7.41 p *<* .001), and a main effect of hemisphere (F(1,180) = 5.42 p *<* .05). As also shown in figure 4a, the average preferred duration decreased significantly from occipital to frontal regions, approaching the mean of the full range of tested durations in the latter (all t(180) *>* 4.19, p *<* 0.005 comparing V1-2d with IPS1, IPS2, IPS3, and FEFs; all t(180) *>* 3.28, p *<* 0.05 comparing V3AB and V5/MT with FEFs). The full set of statistics is reported in the supplementary tables 8 - 10; all reported p values were Bonferroni corrected for multiple comparisons. Beside the average *µ_d_* values, what is worth noticing here is the spread of the distributions which is skewed towards longer durations in V1-2d up to V5/MT and becomes much broader from IPS1 to FEFs. These results indicate that in early visual areas neuronal populations maximally respond to longer durations, in line with our previous observation that in these regions the best fitting model is the CMST. In parietal and frontal visual areas instead, there are neural populations tuned to a wide range of durations, in agreement with the prominence in these regions of vertices best fitted by the GT model (see figure 2).

**Figure 4:**
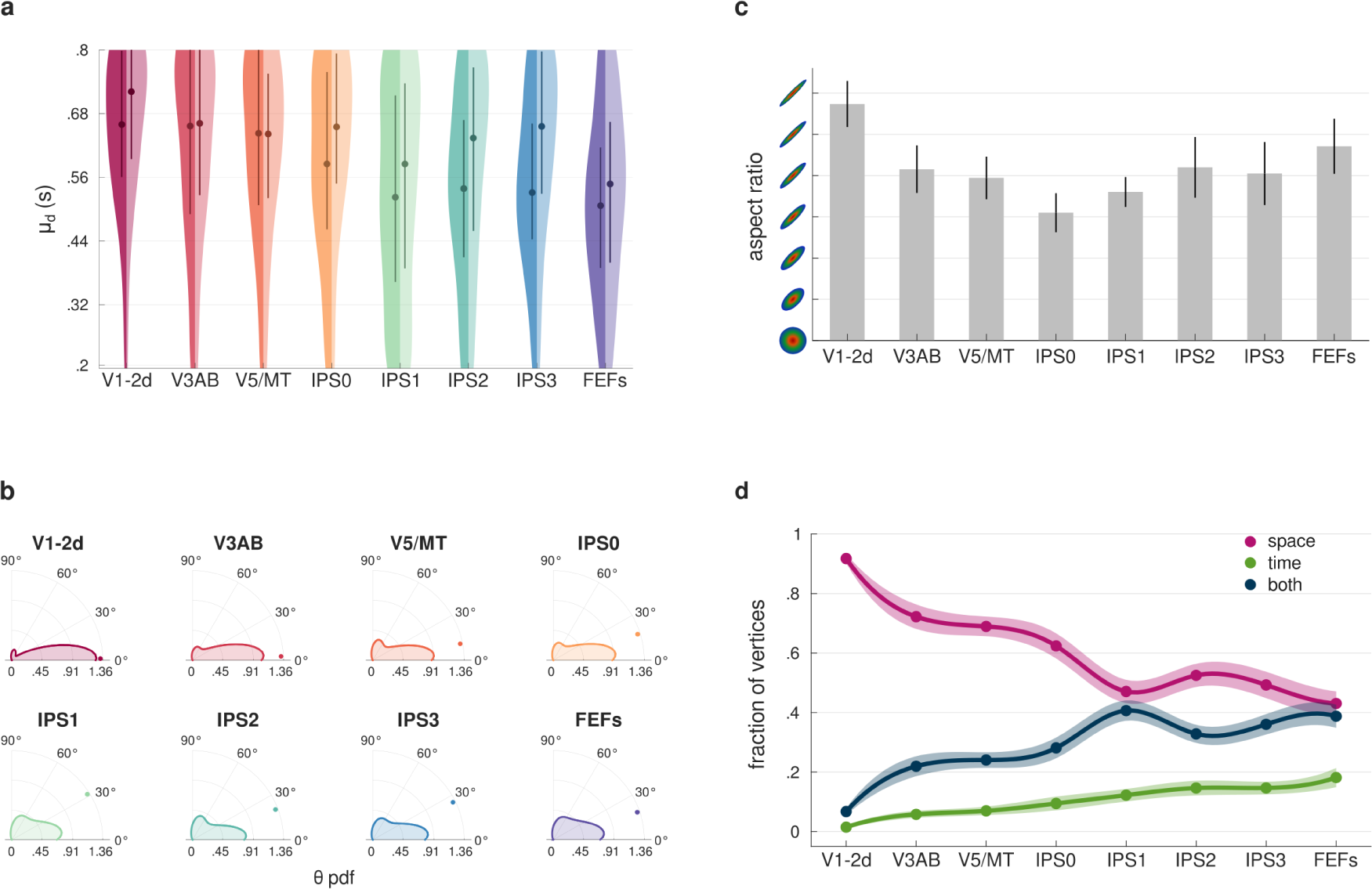
Changes in pRFs parameters. **(a)** Each violin plot represents the group-level distribution of pRFs preferred duration (*µ_d_* parameter estimated by the GST model) in the different ROIs. The left side of the distributions refers to the left hemisphere (darker shades), while the right side refers to the right hemisphere (lighter shades). Dots indicate the median of each distribution, thick lines represent the interquartile range. The distributions’ kernels were estimated with 15% bandwidth. See *pRFs duration preference*. **(b)** Each polar plot displays the distribution of pRFs orientation (*θ* parameter estimated by the GST model) in the different ROIs. The pRF orientation reflects the contribution of either one or two stimulus’s dimensions in generating the response function. Values of 0 and 90 reflect the contribution of a single dimension, whereas values in between indicate a contribution of two dimensions. The dots represent the median of each distribution: V1-2d = 0.4*^◦^*, V3AB = 1.9*^◦^*, V5/MT = 10.3*^◦^*, IPS0 = 16.9*^◦^*, IPS1 = 30.4*^◦^*, IPS2 = 19.7*^◦^*, IPS3 = 24.6*^◦^*, FEFs = 17.7*^◦^*. The distributions’ kernels were estimated with 15*^◦^* bandwidth. See *pRFs orientation*. **(c)** Each bar plot represents the median pRFs aspect ratio across participants and hemispheres for each ROI. The aspect ratio was computed as the ratio between the pRF major and the minor axes (*σ* parameters estimated by the GST model), and it describes the sensitivity of the neuronal response. Round shapes (aspect ratio = 1) indicate that the neuronal response is equally sensitive to changes in both stimulus position and duration, while elongated shapes (aspect ratio *>* 1) indicates a greater sensitivity to changes in one dimension only. In the plot, values span from 1 to 7. The error bars represent the standard error of the median. See *pRFs aspect ratio*. **(d)** The group-level fraction of vertices assigned to each type of selectivity is plotted for each ROI. The pRFs selectivity was derived combining the *σ* and the *θ* parameters estimated by the GST model. This indicates the stimulus dimension (i.e., either the spatial position, the duration, or both) mainly leading the neural response. Each dot is the mean fraction of vertices, across participants and hemispheres, displaying a given type of selectivity (pink for space, green for time, and blue for both). The shaded area is its standard error. For visualization purposes we also plotted lines obtained with a spline interpolation across dots. See *pRFs selectivity*. ROI legend as in figure 3.

We next checked the effect of changes in stimulus’s duration on eccentricity preferences (i.e., *µ_s_*) by looking at the consistency of the *µ_s_*parameter estimated with the retinotopic mapping runs with the *µ_s_* obtained from the GST model (see *pRFs eccentricity preference*). We computed the Kendall’s *τ* correlation coefficient between the two sets of eccentricity preferences for each ROI and participant. Z-scores transformed correlation coefficients were entered in a LME model with ROI and hemisphere as factors and subjects as random intercept (model formula: z-score Kendall’s *τ ∼ ROI ∗ Hemisphere* + (1*|subjectID*), marginal *R*^2^: 0.49, conditional *R*^2^: 0.53). Type III ANOVA on model estimates showed a main effect of ROI (F(7,180) = 29.26 p *<* .0001), and no effect of hemisphere nor interaction. The main effect of ROI indicates that the correlation between the two estimates of eccentricity preferences changed along the visual hierarchy (see supplementary figure 3). Specifically, there was a gradual worsening of correlation coefficients from early visual areas to parietal and frontal regions (all t(180) *>* 4.63, p *<* 0.0005 comparing V1-2d with all the other ROIs; all t(180) *>* 3.86, p *<* 0.005 comparing V3AB with IPS0, IPS1, IPS2, IPS3, and FEFs; all t(180) *>* 2.89, p *<* 0.05 comparing V5/MT with IPS1, IPS2, IPS3, and FEFs). See the supplementary tables 11 and 12 for the full set of statistics; all reported p values were Bonferroni corrected for multiple comparisons. The better correlation of eccentricity preferences in occipital cortex might suggest that in early visual regions BOLD responses mainly reflect changes in stimulus spatial positions, whereas from parietal cortex onward they likely reflect changes in both stimulus’s position and duration.

Next we considered other properties of the GST tuning function, i.e., its orientation, its sensitivity and the combination of orientation and sensitivity. The orientation of the tuning function (i.e., *θ* parameter of the pRF) tells whether time and space contribute independently or jointly in generating the brain response. In this respect, it informed us about the number of stimulus’s dimensions driving the response. Orientations close to either 0*^◦^* or 90*^◦^* indicate a response driven by a single stimulus’s dimension, whereas those between 0*^◦^* and 90*^◦^* indicate a response driven by both dimensions. Figure 4b shows the group-level distribution of the *θ* parameter within each ROI. To test their differences across ROIs, we used the Fisher’s non-parametric test and compared participants’ median *θ* values (see *pRFs orientation*). Results showed a significant increase of the median *θ* parameter from occipital to frontal ROIs (*χ*^2^-Pg(7) = 30.77, p = 0.0001), with a significantly lower value in V1-2d (median *θ* value = 0.4*^◦^*) compared to V5/MT, IPS0, IPS1, IPS2, IPS3, and FEFs (median *θ* values *>* 10.3*^◦^*, all *χ*^2^-Pg(1) *>* 12.46, p *<* 0.05). We also found a significantly lower median *θ* in V3AB compared to IPS1 (*χ*^2^-Pg(1) = 18.61, p = 0.0028). The full set of statistics is reported in supplementary tables 13 and 14; p values were estimated using a 9999 iteration random permutation test and were Bonferroni corrected for multiple comparisons. These findings indicate that neuronal responses in early visual areas (V1-2d, V3AB) are primarily driven by changes of one stimulus dimension, while at higher levels of the visual processing hierarchy, responses are increasingly modulated by the interplay between the stimulus’s position and duration.

We then assessed the sensitivity of the response function in each ROI (see *pRFs aspect ratio*). The sensitivity, which defines the shape of the pRF, was computed as the ratio between the major and the minor axis of the response function (i.e., the ratio between *σ_max_*and *σ_min_*). A thin elongated shape indicates a neuronal response greatly sensitive to changes of a stimulus dimension only, whereas a completely round shape reflects a neural response equally sensitive to changes in both stimulus dimensions. Figure 4c shows for each ROI the median aspect ratio across participants and hemispheres. The plot highlights a U-shaped pattern of aspect ratio changes along the visual hierarchy, with aspect ratios decreasing from V1-V2d to IPS0 and then slightly increasing from IPS0 to FEFs. To test these differences across ROIs, we used a LME model with ROI and hemisphere as factors and subjects as random intercept (model formula: Aspect Ratio *∼ ROI ∗ Hemisphere* + (1*|subjectID*), marginal *R*^2^: 0.10, conditional *R*^2^: 0.26). Type III ANOVA on model estimates showed a main effect of ROI (F(7,180) = 2.80 p *<* .01) and hemisphere (F(7,180) = 4.88, p *<* 0.05). Specifically, the aspect ratio was significantly lower in IPS0 compared to V1-V2d in the right hemisphere only (t(180) = −3.92, p = 0.004). The full set of statistics is reported in supplementary tables 15-17; all p values were Bonferroni corrected for multiple comparisons. These results indicate that the sensitivity of neuronal responses varies along the visual hierarchy, and in early portions of IPS (IPS0 and IPS1) neuronal populations were more sensitive to changes in both stimulus position and duration compared to the other cortical regions.

Finally, we combined the orientation (*θ* parameter) and the sensitivity (*σ* parameters) of the response function. This helped us to identify *which* stimulus dimension (eccentricity, duration, or both) mainly drives the neuronal response in each vertex, and to classify the vertices accordingly (see *pRFs selectivity*). Figure 4d shows the group-level results averaged across hemispheres. A three-way repeated measure ANOVA was used to analyze these data with stimulus’s dimension, hemisphere and ROI as factors. The ANOVA revealed a significant effect of stimulus’s dimension (F(1,24) = 72.16, p *<* .0001) and, more interestingly, a significant interaction between stimulus’s dimension and ROI (F(14,168) = 17.43 p *<* .0001). The interaction indicates that in different ROIs neuronal responses were selective for different stimulus’s dimensions. Specifically, in V1d-V2d, V3AB, V5/MT, and IPS0 the selectivity for the spatial dimension prevailed over that for the temporal dimension and their combination (all t(576) *>* 7.77, p *<* 0.0001). However from V3AB onwards the selectivity for the spatial dimension gradually decreased (all t(576) *>* 4.42 comparing V1-2d with V3AB, V5/MT, and IPS0), whereas the selectivity for both space and time dimensions increased (all t(576) *<* −3.45, p *<* 0.05 comparing V1-2d with V3AB, V5/MT, and IPS0). In IPS1 and FEFs the selectivity for the spatial dimension and for both space and time were equally represented, whereas in IPS2 and IPS3 both types of selectivity were not significantly different from IPS0. The selectivity for the temporal dimension was always under-represented in all ROIs (all t(576) *<* −3.66, p *<* 0.0001 comparing temporal selectivity to the other types in all ROIs - excluding the comparison with the selectivity for both dimensions in V1-2d), however it significantly increased from occipital to frontal regions (t(576) = −3.78, p = 0.005 comparing V1-2d and FEFs). See the supplementary tables 18 - 22 for the full set of statistics; all reported p values were Bonferroni corrected for multiple comparisons. These results indicate that in early visual areas (from V1d-V2d to MT/V5) neuronal populations exhibit higher selectivity for changes of the spatial dimension of the stimulus, whereas in parietal and frontal visual areas (IPS and FEFs) there are also neuronal populations selective for concurrent changes of both stimulus’s eccentricity and duration.

Overall, these findings show a transition in brain responses to changes of stimulus’s eccentricity and duration. In early visual areas i.e., from V1-2d to V3AB, responses are mainly driven by spatial changes (i.e., lower *θ* values, more elongated pRF shapes and higher fraction of vertices showing spatial selectivity). From V5/MT onwards and, more decisively in early portions of IPS (IPS0 and IPS1) response seems driven by changes of both stimulus’s dimensions (*θ* values around 30*^◦^*, rounded pRF shapes and neuronal populations selective to either space, time or both). In this respect, it is noteworthy that common responses to stimulus position and duration occurred in parietal areas where a selectivity to the full range of tested durations was observed (see the *µ_d_* distributions in figure 4a).

### How duration maps are associated with eccentricity maps

The relationship between duration and spatial processing was finally assessed by looking at the relationship between duration and eccentricity maps. Specifically, we studied if and how the spatial distribution of preferred durations (*µ_d_*) estimated by the GST model was associated with that of preferred eccentricities obtained by modeling the retinotopy data (i.e., purely/standard eccentricity maps). To achieve this goal, we employed two distinct spatially informed analysis tools - spatial gradients and Moran’s I statistics - and focused the analyses on the visual ROIs identified using the retinotopic mapping runs (see *Custom-made ROIs*). Figure 3 illustrates eccentricity and duration maps of one sample participant with ROIs marked by blue, red, and white lines.

First, to identify how maps unfold on the cortical surface we used a method as data-driven as possible like the computation of spatial gradients. Spatial gradients were computed for each eccentricity and duration map in all participants (see *Spatial gradients analysis*). This method allowed us to determine the spatial direction of preference change at each cortical location. To identify the *main* direction of change in preference (i.e., the main direction of map’s unfolding), we summed the gradient vector field within a map for each ROI and subject. This sum results in a single vector to which we refer as global gradient. Figure 5a-b shows the result of this procedure for maps in the left hemisphere of one participant (for spatial gradients of all participants see supplementary figures 5-8). Eccentricity (panel a) and duration (panel b) maps are presented with their corresponding isolines (i.e., lines joining locations with equal preference’s values in the map); global gradients are shown in the insets.

**Figure 5:**
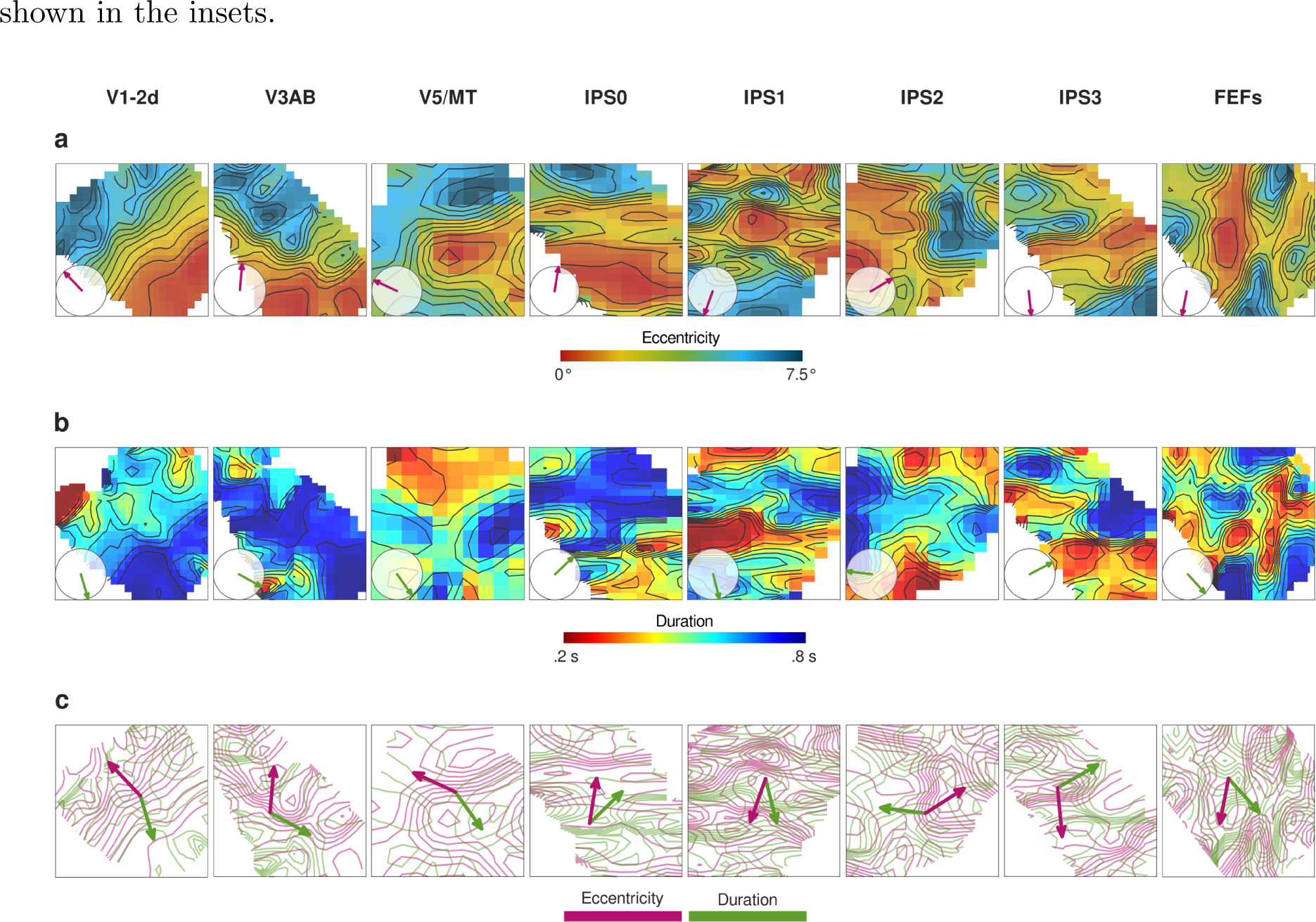
Eccentricity and duration spatial gradients. Eccentricity **(a)** and duration **(b)** maps are shown with their corresponding isolines (in black) for all ROIs of the left hemisphere of one participant. Isolines are separated by 50 ms change of duration preference and 0.5*^◦^* of eccentricity preference. The insets display the global gradient (i.e., the main direction of change) of each map, computed as the sum of its gradient vector field. Panel **(c)** displays eccentricity and duration isolines overlaid to show the gradients intersection. The two global gradients are plotted together to highlight the angle (*αg*) in between. We considered *αg* as a measure of the spatial relationship between preference changes in eccentricity and duration maps. Vectors were normalized (by dividing the vector by its magnitude) to ease visualization. See *Spatial gradients analysis*. ROI legend as in figure 3.

Figure 6a displays individual global gradients of duration (green arrows) and eccentricity (pink arrows) maps. Upon visual inspection it seems that in early visual areas (V1-2d, V3AB) eccentricity global gradients are consistent across subjects; this consistency though becomes weaker from V5/MT onwards (see also [16]). Duration global gradients instead are highly variable across subjects in all ROIs. The presence of a topographic relationship between eccentricity and duration maps can be inferred from the angles between their respective gradient vector fields. This is shown in figure 5c as superimposition of eccentricity and duration isolines (see supplementary figures 9 and 10 for all participants). To identify the relationship between topographies we then computed the angle between eccentricity and duration global gradients (i.e., *α_g_*, see the arrows in figure 5c) in each participant and ROI. Figure 6b illustrates for each ROI the group-level distribution of *α_g_* values (see also supplementary figure 4). At a visual inspection it seems that these distributions of *α_g_* are different in the different ROIs. Specifically, in V3AB, V5/MT, IPS1, and FEFs the distribution of *α_g_* is widespread, indicating an inconsistent association between eccentricity and duration maps in these cortical locations. On the other hand, in V1-2d and IPS2 the *α_g_*values mainly fall between 150*^◦^* and 180*^◦^*, and in IPS0 and IPS3 *α_g_* are most frequently close to 90*^◦^*. These results suggest the existence of ROI-specific spatial relationships between duration and eccentricity maps. In V1-2d and in IPS2 the two maps change along the same orientation on the cortical surface but in opposite direction (*α_g_*close to 180*^◦^*). In IPS0 and IPS3 instead maps change independently on the cortical surface, following orthogonal directions (*α_g_* close to 90*^◦^*). It is worth emphasizing here that in V1-2d it is not fully appropriate to talk about duration maps, since first, in this area the best pRF fitting model was CMTS (see figure 2) and second, the fitting with the GST model revealed a distribution of duration preferences skewed towards long durations (see the analysis on the *µ_d_* parameter previously described). For these reasons, the topographic relationship between eccentricity and duration maps in V1-V2d has to be taken with caution. In contrast, in IPS0, IPS2, and IPS3, where all durations are represented, results clearly showed that duration and eccentricity maps changed maintaining a very specific relationship. To statistically prove a difference in the *α_g_*distributions along the different cortical locations in the hierarchy, we ran the Fisher’s non-parametric test on participants’ median *α_g_*. Results showed significant changes in the median values along the cortical hierarchy (*χ*^2^-Pg(7) = 27.08, p = 0.0003). Specifically, the median of the *α_g_* distribution decreased from V1-2d (median *α_g_*= 147*^◦^*, spread = 18.33*^◦^*) to FEFs (median *α_g_*= 80*^◦^*, spread = 26.36*^◦^*). The only exception to this decrease was observed in IPS2 (*α_g_* = 127*^◦^*, spread = 27.50*^◦^*), where the median of the distribution was not significantly different from V1-2d (*χ*^2^-Pg(1) = 3.85, p = 1). In V5/MT, IPS0, IPS3, and FEFs, median *α_g_*values were significantly lower than in V1-2d (all *χ*^2^-Pg(1) *>* 12.46, p *<* 0.05). The full set of statistics is reported in the supplementary tables 23 and 24. P values were estimated using a 9999 iteration random permutation test and were Bonferroni corrected for multiple comparisons.

**Figure 6:**
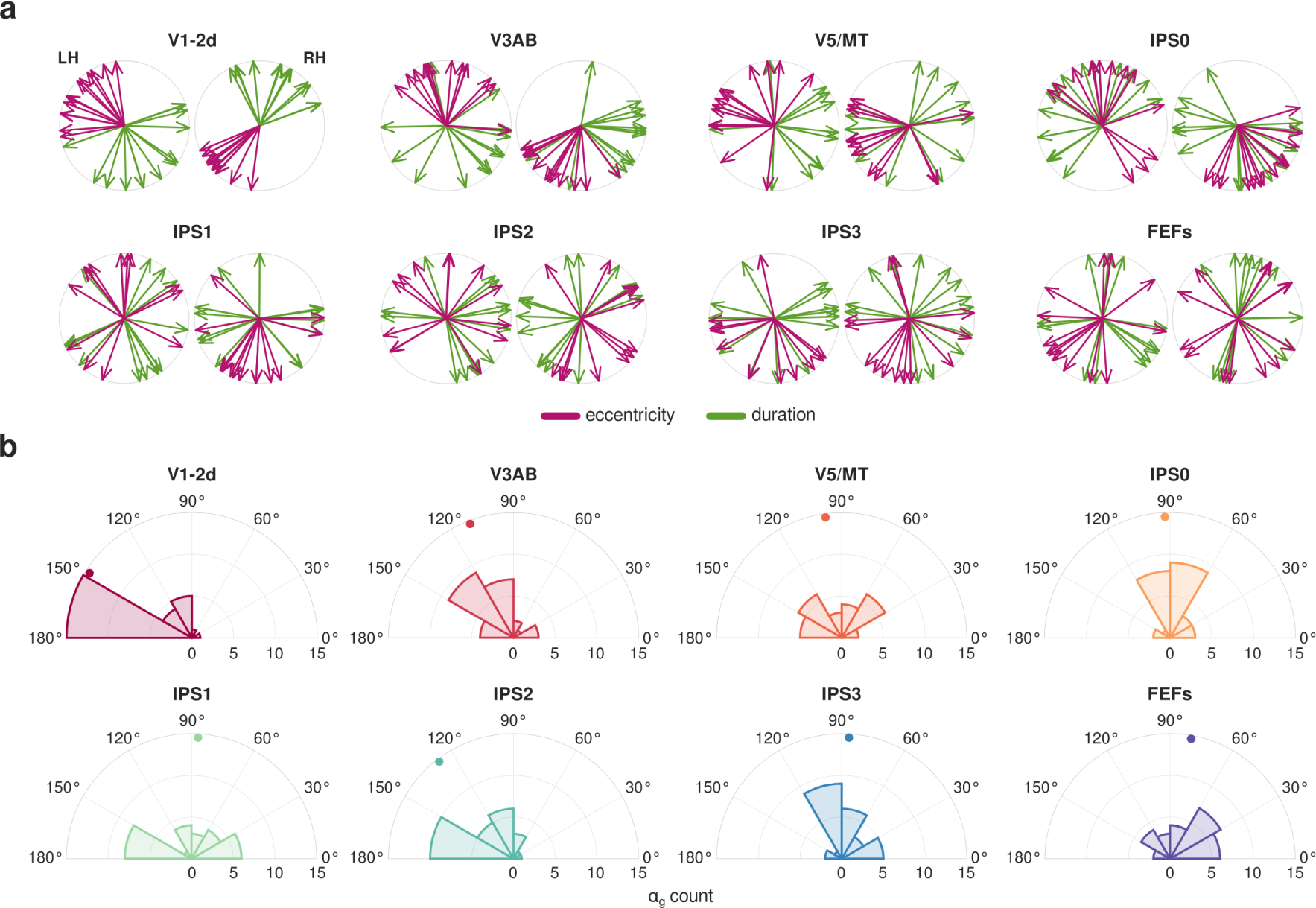
Global gradients and angles. **(a)** Arrows represent individual eccentricity (in pink) or duration (in green) global gradients for each ROI in the left (polar plot on the left) and the right (polar plot on the right) hemisphere. The global gradient represents the main direction of change of a map. Vectors were normalized (by dividing the vector by its magnitude) to ease visualization. **(b)** Polar histograms display for each ROI the distribution across participants and hemispheres of angles between eccentricity and duration global gradients (*αg*). We considered *αg* as a measure of the spatial relationship between eccentricity and duration maps. An *αg* of 0*^◦^* or 180*^◦^* signifies that maps change on the cortical surface following the same orientation in either the same or the opposite direction. An *αg* of 90*^◦^* implies that maps change independently, along orthogonal directions. The dots represent the median of each distribution: V1-2d = 147*^◦^*, V3AB = 111*^◦^*, V5/MT = 98*^◦^*, IPS0 = 93*^◦^*, IPS1 = 87*^◦^*, IPS2 = 127*^◦^*, IPS3 = 86*^◦^*, FEFs = 80*^◦^*. See *Spatial gradients analysis*. ROI legend as in figure 3.

The analysis of spatial gradients revealed that eccentricity and duration maps are linked when considering how maps change along the cortical surface. However, this method does not consider any direct association between eccentricity and duration preferences, which might indeed represent another aspect of the relationship between the maps. To this aim, we calculated for each ROI and participant the bivariate global Moran’s I, which served as an indicator of the spatial correlation between eccentricity and duration maps. Essentially, this statistics measures to what extent the eccentricity preference at each cortical location within a map correlates with the averaged duration preference of its neighbors and summarizes the nature of the overall spatial relationship between the two preferences. The bivariate Moran’s I was computed for each ROI and each subject considering vertices with a duration preference estimated with the GST model and an eccentricity preference obtained from the retinotopic mapping runs (see *Bivariate Moran’s I statistics*). Supplementary figure 11 displays the group-level distributions of this statistics. A Moran’s I close to 0 indicates spatial randomness (i.e., no spatial association, hence no clustering) between the two preferences. Values higher or lower than zero indicate spatial association, characterized by respectively a positive (i.e., vertices with high or low eccentricity preferences tend to be close to vertices with same high or low duration preferences), or a negative (vertices with high eccentricity preferences tend to be close to those with low duration preferences, and vice versa) correlation. The distributions of bivariate Moran’s I showed a consistent negative Moran’s I in V1-2d only (median = −0.4 in both hemispheres), while in the other ROIs the median Moran’s I was close to 0 (between −0.16 and 0.04 in the left hemisphere and between −0.18 and 0.03 in the right hemisphere). These results indicate that eccentricity and duration preferences are clustered in V1-2d only, where high eccentricity preferences are likely to be located near low duration preferences, and vice versa. In all the other ROIs, there was no spatial association between eccentricity and duration preferences.

To summarize, the results of this section showed the presence in IPS of a relationship between duration and eccentricity maps in their unfolding along the cortical surface. In IPS2, the two maps changed in opposite directions (*α_g_* mainly at 180*^◦^*), in IPS0 and IPS3 their change in preference instead followed orthogonal directions (*α_g_* mainly at 90*^◦^*). However, with the exception of V1-2d, in all the rest of cortical locations there was no spatial association between duration and eccentricity preferences.

### Comparing duration with eccentricity maps

In the final set of analyses we explored the link between spatial and temporal processing by comparing the spatial properties of duration and eccentricity maps, i.e., their degree of clustering, the strength and the extent of their topographic organization. To achieve this goal, we used two spatial statistical methods, Moran’s I and variogram.

First, we checked the “quality” of spatial clustering within the maps by computing the univariate global Moran’s I for each eccentricity and duration map independently (see *Univariate Moran’s I statistics*). The univariate Moran’s I is a measure of spatial autocorrelation and in our case it quantified the correlation between each vertex’s preference and the averaged preference of its 12 nearest neighbors. Figure 7a shows the group-level distributions of this statistics. As previously described for the bivariate Moran’s I, a value of 0 indicates spatial randomness, whereas values of 1 or −1 indicate spatial clustering, characterized by either a positive or negative spatial autocorrelation. The results revealed a positive Moran’s I for both eccentricity and duration maps, indicating spatial clustering of vertices with similar preferences. In eccentricity maps, median Moran’s I values were above 0.76 in all ROIs, although there was a gradual decay of Moran’s I values along the cortical hierarchy (i.e., the highest value was in V1-2d, the lowest in FEFs). In duration maps instead the Moran’s I values remained relatively stable along the hierarchy, with median values above 0.58 in all ROIs and with broader distributions than eccentricity maps. These results indicate that duration preferences, while less clustered than eccentricity preferences and with higher inter-individual variability in their spatial arrangement, are still organized in a non-random fashion in all ROIs.

**Figure 7:**
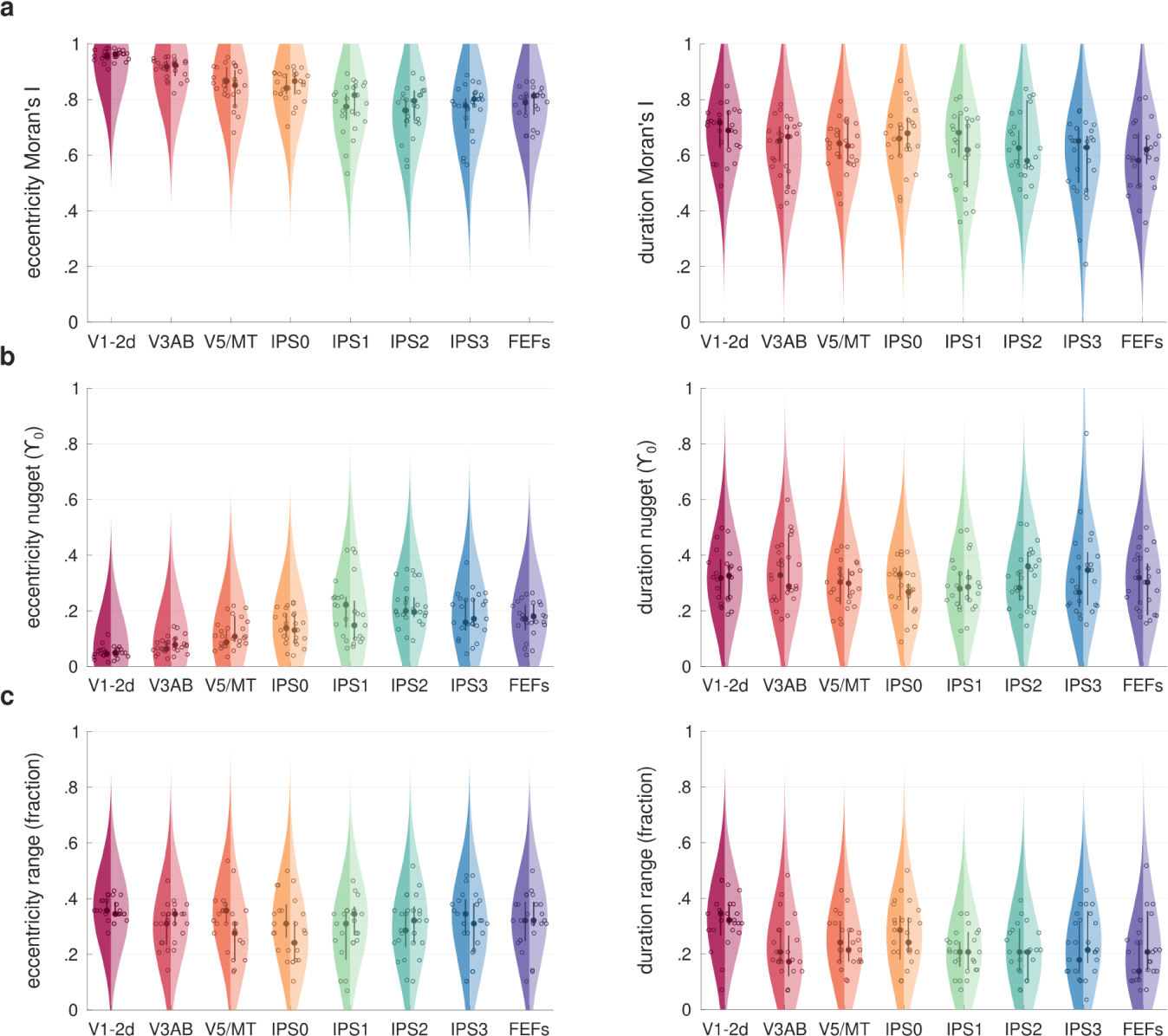
Group-level distributions of spatial statistics measures: Moran’s I, nugget, and range. Each violin plot shows the group-level distribution of the univariate Moran’s I **(a)**, the variogram’s nugget **(b)** and the variogram’s range **(c)** of eccentricity (leftward panel) and duration (rightward panel) maps in the different ROIs. The left side of the violins (darker shade) refers to the left hemisphere, the right side refers to the right hemisphere (lighter shade). Dots represent the median of the distribution, and circles correspond to individual data points. Thick lines show the interquartile range, the distributions’ kernels were estimated with 15% bandwidth. **(a)** The Moran’s I is a measure of spatial autocorrelation. A value of 0 indicates spatial randomness, whereas values of 1 or −1 indicate spatial clustering, characterized by either a positive or negative spatial autocorrelation. **(b)** The nugget corresponds the variance between vertices located at the minimum distance in the map, and it reflects the strength of the spatial autocorrelation in the map (lower values indicate a stronger spatial autocorrelation). Nuggets are expressed in units of variance of the entire map. **(c)** The range is the distance between vertices required to reach the variance of the whole map, and it assesses the extent of the spatial autocorrelation in the map. Ranges are expressed as a fraction of the full extent of the map. See *Spatial properties of eccentricity and duration maps*. ROI legend as in figure 3.

After assessing the presence of clustering in eccentricity and duration maps, we next explored the strength and the extent of their spatial autocorrelation. To achieve this goal we built the experimental variogram of each map. The variogram is a method used to compute spatial autocorrelation, but differently from the Moran’s I used previously, it is based on variance estimation, and it assesses the similarity in preference of vertices located at varying distances from each other (see *Variogram*). For example in a map one would expect high similarity (i.e., low variance) between neighboring vertices and low similarity (i.e., high variance) between spatially distant ones. We extracted two parameters of the variogram: the nugget and the range. The nugget corresponds to the variance observed at the shortest distance between vertices in the map, and it quantifies the strength of the spatial autocorrelation in the map. Low nuggets indicate low randomness of neighboring vertices and in turn a strong spatial autocorrelation. The range instead is the spatial distance required to reach the sample variance, and it appraises the extent of the spatial autocorrelation in the map. High ranges indicate a greater extent of the spatial autocorrelation. Figure 7 shows the group-level distribution of eccentricity and duration nuggets (panel b) and ranges (panel c) for each ROI. We tested individual nuggets and ranges of each eccentricity and duration map with two different LME models with ROI, map type (i.e., eccentricity or duration map), and hemisphere as factors and subjects as random intercept (nugget model formula: Nuggets *∼ ROI ∗ MapType ∗ Hemisphere* + (1*|subjectID*), marginal *R*^2^: 0.54, conditional *R*^2^: 0.56; range model formula: Ranges *∼ ROI ∗ MapType ∗ Hemisphere* + (1*|subjectID*), marginal *R*^2^: 0.24, conditional *R*^2^: 0.25). Concerning nuggets, type III ANOVA on model estimates showed a main effect of ROI (F(7,372) = 6.00 p *<* .001), a main effect of map type (F(7,372) = 396.34, p *<* 0.001), and an interaction between ROI and map type (F(7,372) = 8.32, p *<* 0.001). Specifically, nuggets of eccentricity maps were significantly lower than those of duration maps (t(372) = −19.9, p *<* 0.001), indicating a stronger spatial autocorrelation in eccentricity maps. In addition, differences of nuggets across ROIs depended on the type of map under consideration. In particular, eccentricity nuggets increased from early visual areas to parietal and frontal regions (all t(372) *>* 3.8, p *<* 0.001 comparing IPS0, IPS1, IPS2, IPS3, and FEFs with V1-2d; all t(372) *>* 4.07, p *<* 0.005 comparing IPS1, IPS2, IPS3, and FEFs with V3AB; all t(372) *>* 3.6, p *<* 0.05 comparing IPS1 and IPS2 with V5/MT), whereas duration nuggets showed no significant differences between ROIs. This result indicates that along the cortical hierarchy the strength of the spatial autocorrelation degrades in eccentricity maps, and it remains stable in duration maps. See the supplementary tables 25-27 for the full set of statistics; all reported p values were Bonferroni corrected for multiple comparisons.

Concerning ranges, type III ANOVA on model estimates showed a main effect of ROI (F(7,372) = 5.08 p *<* .001), and a main effect of map type (F(7,372) = 69.64 p *<* .001), while no interaction. Specifically, eccentricity maps showed wider ranges than duration maps (t(372) = 8.34, p *<* 0.001), indicating that the spatial autocorrelation of eccentricity maps extended over a greater portion of the cortical surface compared to duration maps. Moreover, duration ranges in V1-2d were wider compared to the other ROIs except for IPS0 (all t(372) *>* 3.26, p *<* 0.05 comparing V1-2d with V3AB, V5/MT, IPS1, IPS2, IPS3, and FEFs). The wider range observed in V1-2d for duration preferences may be due to the great number of vertices in this area responding maximally to longer duration (i.e., best fitted by the CMTS model). If the majority of the vertices exhibit long duration preference, the variance in the map will be small and the distance to reach the sample variance necessarily large. See supplementary tables 28-30 for the full set of statistics; all reported p values were Bonferroni corrected for multiple comparisons.

Overall, these findings suggest that duration preferences in visual areas are clustered in maps. Duration maps though are different from eccentricity ones. They show a lower degree of spatial autocorrelation (as suggested by both the Moran’s I and variograms’ nugget), and they are smaller in size. Yet, differently from eccentricity maps, they show similar spatial properties throughout the visual hierarchy.

## Discussion

In the present study, by means of spatially-dependent and spatially-independent population response models to stimulus duration, we explored the relationship between the cortical processing and representation of visual duration and stimulus position. Specifically we asked: a) if and how the processing and the representation of the duration of a visual stimulus is routed in the visuo-spatial cortical hierarchy, b) whether the link between duration and spatial position processing entails monotonic or unimodal “Gaussian-like” responses, and c) what is the relationship between retinotopic (eccentricity) and chronotopic maps.

The results show that stimulus duration is encoded in the response amplitude of spatially-selective neuronal populations in early visual cortices (V1-2d, via monotonic responses) and gradually along the cortical hierarchy by spatially-invariant duration-tuned populations (via unimodal responses). Unimodal duration-tuned responses first appear in extrastriate areas V3AB and V5/MT, their presence increase in the IPS and becomes predominant in frontal and premotor areas. In humans, both monotonic and unimodal topographically-organized responses to visual stimulus duration have been reported before in a wide network of brain regions. In visual and parietal cortices both unimodal and monotonic responses were identified [10], [13], [17], whereas in premotor and prefrontal cortices only unimodal responses were reported[11], [12]. Our results, while replicating these previous findings, go beyond them showing for the first time how brain responses to stimulus duration are linked to those to stimulus position and how this link unfolds along the visual cortical hierarchy. The relationship between stimulus duration and position is very tight in early visual cortices, where monotonic responses to stimulus duration are spatially-specific (CSMT is the winning model), i.e., only vertices coding for the tested eccentricity position of the stimulus are active. This relationship becomes a bit looser in extrastriate regions V3AB, V5/MT and in inferior parietal lobule, where multiple response profiles coexist, i.e., responses selective to position only, to position and time together and to time only. In frontal regions (SMA, FEFs and IFS-PCSi) responses to stimulus duration detach from those to stimulus position and spatially-invariant unimodal responses to stimulus duration become the most prominent pRF model (GT is the winning model). These results are in line with the idea of a multi-stage processing of temporal information[9], where duration is extracted locally in early visual cortices by spatially-tuned neuronal units that can be targeted by fast-moving adaptors[1], [3], [6]. And it is subsequently decoded and represented in duration-selective and spatially-independent neuronal populations, susceptible to spatially invariant duration adaptations, in parietal cortex[7], [8], [15], [17]. Our findings corroborate earlier works showing a key involvement of early visual areas in both duration processing and perception[19]– [23]. As suggested by electrophysiological and imaging works in rodents[24], [25], one might imagine that computationally the duration of a visual stimulus is encoded in early sensory areas via accumulation of sensory inputs by means of spatially selective and duration sensitive (via monotonic responses) neuronal populations. This information once integrated, perhaps in extrastriate areas or inferior parietal cortices, might be decoded later in the cortical hierarchy in premotor and prefrontal regions by means of unimodally tuned duration-specific and spatially-invariant neuronal populations. Neuronal populations selective to specific durations have been described before in monkey’s medial-premotor cortex[26], [27]. In this region, for example, the production of millisecond temporal intervals is associated with the activation of duration tuned neurons, with neural trajectories of different speeds and with the sequential activation of different neurons. Importantly, most of these computations take place for both visual and auditory temporal intervals[28]. These results together with many others neuroimaging studies in humans (see for instance [29]–[32] and [33] for a recent review) show a key role of medial premotor cortex (i.e., SMA) in temporal computations across tasks and sensory modalities[34]. A previous work by Hendrikx and colleagues[13] showed the switch between monotonic and unimodal responses to stimulus duration in extrastriate area V5/MT. Here we showed the presence of unimodal tuning to duration earlier in the visual hierarchy, in area V3AB, and we observed that monotonic responses to stimulus duration are tightly linked to spatial processing and coexist with spatially-invariant duration responses until the parietal lobule. In our work compared to Hendrikx and colleagues, by using different neuronal response models that combined durations with spatially-selective responses, we were able to better capture and specify this transition along the cortical hierarchy. In addition, by introducing in our experiment parametric changes in stimulus position, we were also able to better characterize the dependency between duration and eccentricity preferences.

The relationship between space and time appears in its complexity in the intraparietal sulcus, where not only spatially-specific monotonic and spatially-independent unimodal responses to durations coexist, but where brain responses, modeled as a bivariate Gaussian function (GST model), show selectivity to either space and (to a lesser extent) time separately or to both together (i.e., round aspect ratio, medians of the *θ* parameter around 30*^◦^*). This last result is in line with a previous finding from our group showing that the simultaneous change in duration and numerosity of a visual stimulus leads to changes in brain responses from occipital to frontal regions. Compared to baseline conditions where only a single stimulus dimension was manipulated, response functions became more sensitive to both duration and numerosity and their preferences changed. Interestingly, preference changes were more pronounced in parietal and frontal regions[35]. Parietal cortex is the brain area where different kinds of integration processes are known to happen, for example multisensory[36] and visuo-motor[37], and where different visual magnitudes like stimulus size, duration and numerosity are processed. The intraparietal sulcus is also the place where space, time, size and numerosity maps have been found to partially overlap (for a review see [38]). One can therefore imagine that this is the brain area where the information coming from multiple stimulus dimensions, such as position, size and duration is linked and brought together to create a unitary representation of the stimulus at hand[18], [39].

The IPS is also the area where the spatial progression of eccentricity and time topographies is linked. This relationship is not a point-wise, one-to-one matching of duration and eccentricity preferences, but rather a link in the way the two maps unfold along the cortical surface. In IPS2 maps change along the same orientation but in opposite directions. In IPS0 and IPS3 maps progression is orthogonal. This result suggests that in different portions of the IPS there is a common principle governing the spatial progression of the maps. As the topographic organization is thought to improve the efficiency of the neural computation and communication[40], we hypothesize that the sharing of the same principles across different maps might serve a similar purpose. The differences observed across ROIs might indicate different steps of the integration between the two stimulus dimensions and point to a complex link between eccentricity and duration maps instead of a simple overlap[38]. Another indication of the complexity of the interaction between eccentricity and duration maps comes from the observation that the correlations between eccentricity maps estimated with the GST model on the experimental data (where eccentricity and duration were varied together) and those estimated on the retinotopic mapping data (where only eccentricity was varied) tend to worsen from parietal cortex onwards. This is probably a consequence of the fact that this area is sensitive to both stimulus’s features, when these features are manipulated together in the same visual object.

Finally, the use of spatial statistics tools (i.e., Moran’s I and variogram) allowed us to assess the spatial properties of duration maps compared to eccentricity ones. The results show that differently from eccentricity maps, whose clustering (univariate Moran’s I) tends to decrease along the visual hierarchy, duration preferences are stable across visual areas. However, compared to eccentricity maps, their degree of clustering is always inferior, their size (range of variograms) is smaller and their variance between nearest neighbors (nugget of variograms) higher. These results show a worse quality of duration maps compared to eccentricity ones. These differences might reflect the peculiarity of duration as stimulus feature to be mapped. First, the duration of a stimulus is fully available only after its offset and second, there are neither sensory organs nor pathways dedicated to its processing. The processing of temporal information might therefore exploit brain mechanisms beneficial for its neural computations and communication (e.g., unimodal tuning functions and topography) but adapt them to the unique nature of time. It is worth emphasizing here that the duration maps analyzed in this study were identified within eccentricity maps. These maps reflect the cortical representation of a visual stimulus’s duration covering a very specific location in the visual field. For these reasons, the duration maps presented here may be different from other duration maps existing beyond visual areas[11], [12].

In conclusion in this work we show that time is linked to spatial processing to a different extent depending on the functional stage of the duration processing. Time exploits visuo-spatial circuits and it is encoded in duration sensitive neural units earlier in the visual hierarchy, when time is extracted from sensory inputs. When these inputs are integrated and time becomes an “object” or a representation, in extrastriate areas and IPS for example, it is encoded in both spatially specific and spatially independent neuronal units. At this stage temporal and spatial maps are associated and brain responses are influenced by changes of both stimulus dimensions. Spatially independent units which are unimodally tuned to duration are mainly engaged at the decoding stage, when time is readout in premotor and inferior frontal regions for motor and decision purposes.

## Materials and Methods

### Participants

Thirteen healthy volunteers participated in this study (6 females; mean age = 29.6, SD = 7.3; 2 left-handed participants). All volunteers had normal or corrected-to-normal visual acuity. The experimental procedures were approved by the International School for Advanced Studies (SISSA) ethics committee (protocol number 11773) in accordance with the Declaration of Helsinki, and all participants gave their written informed consent to participate in the experiment.

### Stimuli and Experimental Procedure

#### Stimuli

Participants were presented with visual stimuli displayed on a BOLD screen (Cambridge Research Systems 32-inch LCD widescreen, resolution = 1920 × 1080 pixels, refresh rate = 120 Hz) placed outside the scanner bore at a total viewing distance of 210 cm and viewed via a mirror. The stimuli were colored circular patches of Gaussian noise subtending 1.5*^◦^* of visual angle, changing dynamically frame by frame and presented on a grey background. Each stimulus was constructed by randomly selecting RGB values from a Gaussian distribution of mean = 127 and SD = 35 for each of its pixels and frames. This ensured that the average stimulus luminance was constant and independent of its duration. To prevent the perception of flickers induced by the fast-changing rate in the stimulus, we scaled down its pixel resolution (scaling factor = 12.33). This created a blurring effect that homogenized the local contrasts of the stimulus over frames and minimized possible flickering effects[41]. The entire experimental procedure was generated and delivered using MATLAB and Psychtoolbox-3[42]. An identical set-up was used during participants training.

#### Task and experimental design

Participants were asked to perform a single interval discrimination task in which they had to compare the duration (i.e., display time) of a comparison stimulus to that of a reference stimulus internalized during the training procedure. The task was to report whether the comparison stimulus was longer or shorter than the reference. The reference duration was 0.5 s, and the comparison durations were 0.2, 0.3, 0.4, 0.6, 0.7, and 0.8 s. The comparisons were presented in different spatial positions (i.e., display locations on the screen), which could be either at 0.9*^◦^* or 2.5*^◦^* of visual angle diagonally from the center of the screen in the lower-left or lower-right visual quadrant (see the close-up in figure 1a). Stimuli did not overlap across spatial positions. A white fixation cross (0.32*^◦^* of visual angle) was displayed at the center of the screen throughout the experiment. In each trial, the comparison stimulus was presented in a specific position and entailed a specific duration. After a randomized interval from the offset of the stimulus (stimulus-cue interval (SCI) uniformly distributed between 0.9 and 1.2 s), the participants’ response was cued with a color switch (from white to black) in the fixation cross. The response was allowed within a 2 s window, but no emphasis was placed on reaction times. Participants were instructed to provide their responses by pressing one of two buttons on a response pad with their right index finger or right middle finger to express the choices “comparison longer than reference” and “comparison shorter than reference” respectively. No feedback was provided after the response. A uniformly distributed inter-trial interval (ITI) between 1.8 and 2.5 s interleaved the trials. See figure 1a for a pictorial representation of the trial structure. The stimulus duration varied trial by trial in a pseudorandomized and counterbalanced fashion, whereas its position varied sequentially and cyclically to minimize attentional switching effects on the duration judgement[14]. Each cycle started and ended at 2.5*^◦^* in the lower-left quadrant and comprised a clockwise and counterclockwise presentation of the stimulus in all positions, from 2.5*^◦^* lower-left to 2.5*^◦^* lower-right and backwards (i.e., 2.5*^◦^* L, 0.9*^◦^* L, 0.9*^◦^* R, 2.5*^◦^* R, 2.5*^◦^* R, 0.9*^◦^* R, 0.9*^◦^* L, 2.5*^◦^* L). Each half cycle (i.e., when the presentation order turned from clockwise to counterclockwise and vice versa) was followed by a 2.64 s (2 TR) interval. To ensure a balanced presentation of all combinations of durations and positions within each block, a cycle was repeated six times, and each duration was presented twice in each position, for a total of 48 trials per block. Each participant performed 10 blocks inside the scanner acquired in separate fMRI runs. Duration randomization differed in each block, whereas the position sequence was always the same. Stimuli presentation was synchronized with the scanner acquisition at the beginning and at the middle of each cycle. Participants were instructed to maintain their gaze at the fixation cross while performing the task, and eye movements were monitored online and recorded with an MR-compatible eye-tracking system (R Research Eyelink 1000 Plus) placed inside the scanner bore.

#### Training

Participants underwent a training procedure outside the scanner to familiarize with the stimuli and the task. First, they were asked to internalize the duration of the reference stimulus. In this phase participants passively viewed a 0.5 s stimulus which was presented at the center of the screen three times (inter-stimulus interval uniformly distributed between 1.8 and 2.5 s) in each trial. They were free to complete as many trials as they needed to feel confident they had internalized the duration of the stimulus. Next, participants performed the first training block of the duration discrimination task. The task structure was identical to the one described earlier (see *Task and Experimental Design*), but all comparisons were presented at the center of the screen. This was done to ensure that the participants were able to correctly discriminate the comparisons from the reference stimulus. Finally, participants performed a second training block equivalent to the experimental blocks to familiarize with the experimental procedure. Throughout the training phase, participants received visual feedback about their performance and eye movements.

### Retinotopy runs

Participants underwent two retinotopy runs which allowed for a precise identification of their visual field maps. We used the stimulation paradigm of the Human Connectome Project 7T Retinotopy Dataset[43]. In brief, the stimulus was a bar-shaped aperture filled with a texture of colorful objects at multiple scales on an achromatic pink-noise background. The bars were constrained to a circular region subtending 10*^◦^* of visual angle, and the width of each bar was 1.25*^◦^* of visual angle (1/8 of the mapped visual field). The background was uniformly gray. Four bar orientations (0*^◦^*, 45*^◦^*, 90*^◦^*, and 135*^◦^*) and two motion directions were used, ending up with eight different bar configurations. Each run consisted of blank periods and bar movements as follows: 16 s blank period, 4 bar movements (directions: right, up, left, down) of 32 s each, 12 s blank period, 4 bar movements (directions: upper-right, upper-left, lower-left, lower-right) of 32 s each, 16 s blank period. The last 4 s of each 32 s bar movement were blank. Bars apertures were animated at 15 Hz by randomly selecting one of 100 texture images (avoiding the consecutive presentation of the same texture). A fixation cross (0.32*^◦^* of visual angle) was displayed at the center of the screen throughout the experiment, and participants were instructed to maintain their gaze on it. To aid fixation, the color of the cross switched randomly between green, red, and white, and participants were asked to press a button whenever the color changed; furthermore, a semitransparent fixation grid was superimposed on the display throughout the experiment.

### MRI acquisition

MRI data were acquired with a Philips Achieva 7T scanner equipped with an 8Tx/32Rx-channel Nova Medical head coil. T2*-weighted functional images were acquired using a three-dimensional EPI sequence with anterior-posterior phase encoding direction and the following parameters: voxel resolution = 1.8 mm isometric; repetition time (TR) = 1.32 s; echo time (TE) = 0.017 s; flip angle = 13 degrees; bandwidth = 1750 Hz/px. Universal kt-points pulses were used to achieve a more homogeneous flip angle throughout the brain[44]. The matrix size was 112×112×98, resulting in a field of view of 200(AP) x 200(FH) x 176.4(LR) mm. At the end of each run, 4 volumes were acquired with the opposite phase encoding direction in order to perform susceptibility distortion correction (see *MRI data preprocessing*). A minimum of 190 volumes (acquisition time *≈* 4 minutes) was acquired for each experimental run. For each retinotopy run, 228 volumes were acquired (acquisition time = 5 minutes). Peripheral pulse and respiratory signals were recorded simultaneously with the fMRI data acquisition using the Philips MR Physiology wireless recording system. The finger clip of the peripheral pulse unit was placed on the subject’s left ring finger, and the respiratory sensor was placed over the diaphragm of the subject and secured with a band. Eye movements were monitored and recorded with an eye-tracking system (SR Research Eyelink 1000 Plus) mounted onto a hot-mirror system and located inside the scanner bore. High-resolution T1-weighted images were obtained using the MP2RAGE pulse sequence[45] optimized for 7T (voxel size = 0.7 x 0.7 x 0.7 mm, matrix size = 352 x 352 x 263).

### MRI data processing

Pulse-oximetry and respiratory components were regressed out from the BOLD traces before the preprocessing. We converted the physiological signals into slice-based regressors using RetroTS.py (AFNI), and we performed the Retrospective Image Correction with a custom routine based on 3dretroicor (AFNI). This procedure was applied only whether physiological signals were reliable in their frequency spectrum (physiological signals were removed in 108 out of 156 runs; both pulse and respiratory signals were removed in 67/108 runs, only pulse signals in 29/108 runs, only respiratory signals in 12/108 runs). Data were preprocessed using fMRIPrep 21.0.2[46], [47], which is based on Nipype 1.6.1[48], [49]. See the supplementary methods for details about the pipeline. The BOLD time series of the retinotopy runs were also high-pass filtered by removing the first six components from the discrete cosine transform of the data, and then were converted to percentage of signal change.

### Behavioral performance analysis

The behavioral analysis aimed to assess whether duration discrimination performance was affected by the display location of the stimulus. The individual behavioral data were grouped by the position of the stimulus, and the mean fraction of “comparison longer than reference” responses was computed for each comparison duration to build individual psychometric curves. In addition, we computed the mean fraction of “comparison longer than reference” responses across participants for each comparison duration to build group psychometric curves (figure 1b). All psychometric curves were fitted using the MATLAB built-in function glmfit with a logit link function. For each participant we derived the Point of Subjective Equality (PSE; i.e., the comparison duration equally likely to be judged as longer or shorter than the reference duration) and the Just Noticeable Difference (JND; i.e., the minimum difference between reference and comparison stimulus to be detected 75% of the times) which are respectively a proxy for bias and sensitivity of the duration discrimination judgments. Individual PSE and JND values were analyzed with two linear mixed effect (LME) models, using the lme4 R package[50], with the following formulas:

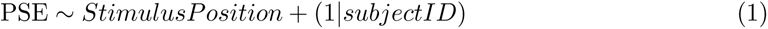

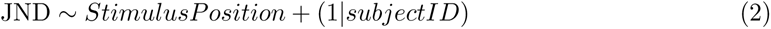

LME model variance explained was computed using the MuMIn package[51]. The Satterwaite’s method[52] implemented in the lmer package[53] was used to estimate the degrees of freedom for the LME model ANOVA.

### General Linear Model (GLM) analysis

Functional data resampled on the cortical native surface were initially analyzed using a GLM approach with the GLMdenoise toolbox[54]. For each run, the design matrix included one regressor for each combination of comparison duration and position time-locked to the offset of the stimulus (events of interest) and one regressor time-locked to the onset of the response (event of no interest). Thus, 25 events were modeled (6 stimulus durations x 4 stimulus positions + response). As GLM-denoise automatically estimates noise regressors, no motion correction parameters were entered in the procedure. Regressors were convolved with the canonical hemodynamic response function. For each subject, this procedure yielded a set of 100 bootstrapped beta weights for each vertex. The median beta weight across bootstraps, converted to percentage of signal change, was used in the following analysis steps.

### Population Receptive Field (pRF) modeling

In order to characterize the tuning properties of BOLD responses to different combinations of stimulus duration and position, we applied the pRF modeling[55] and predicted GLM betas relative to stimulus offsets. We tested five different pRF models to account for different neuronal responses elicited by our experimental manipulations.

#### Compressive Monotonic Time (CMT) model

The CMT model assumes a monotonic neuronal response to stimulus duration *d*, invariant to stimulus position. The neuronal response is thus described by the following equation:

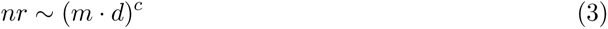

where *m* is the slope of the linear increase and *c* is an exponent modulating the compression of the response[10].

#### Compressive Monotonic Time and Gaussian Space (CMTS) model

The CMTS model assumes a monotonic neuronal response to stimulus duration (as the CMT model) coupled with a unimodal response to stimulus position *s*, represented by a Gaussian function. The model is described by the following equation:

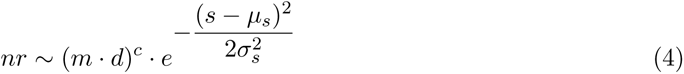

where *µ_s_*represents the preferred spatial position (i.e., the stimulus position eliciting the greatest neuronal response) and *σ_s_* represents the sensitivity of the response to changes in stimulus position.

#### Gaussian Space (GS) model

The GS model assumes a unimodal neuronal response to the spatial position of the stimulus (as in CMTS), invariant to its duration, and it is described by a univariate Gaussian function:

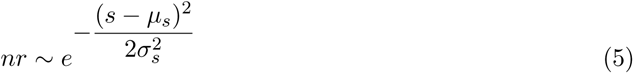

#### Gaussian Time (GT) model

The GT model describes a unimodal neuronal response to the duration of the stimulus, invariant to its spatial position, thus it is the reciprocal of the GS model:

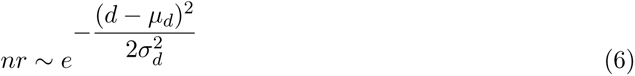

where *µ_d_* represents the preferred duration (i.e., the stimulus duration eliciting the greatest neuronal response) and *σ_d_*the sensitivity of the response to changes in stimulus duration.

#### Gaussian Space-Time (GST) model

The GST model assumes a unimodal neuronal response to both the temporal and the spatial dimensions of the stimulus. Therefore, it is described by a bivariate Gaussian function with the following equation:

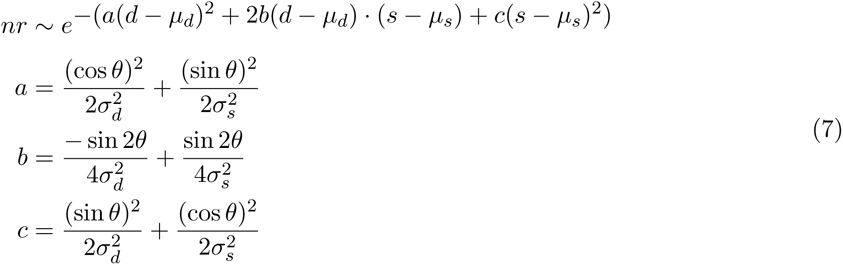

where *θ* represents the orientation of the neuronal response function.

The above-described models are able to capture a variety of neuronal responses to our experimental manipulation, including different kinds of sensitivity that neuronal population might have relative to changes of stimulus duration and position (i.e., purely “space-modulated” responses, purely “time-modulated” responses, and responses modulated by both stimulus dimensions) and different tuning functions for temporal information (i.e., monotonic or unimodal neuronal responses).

#### Fitting procedure

A two-dimensional matrix was used to represent the spatial and temporal dimensions of the stimulus, with durations and positions expressed in arbitrary units from 1 to 100. We built 24 matrices, each representing a unique combination of stimulus duration and position present in our experimental manipulations. These matrices were then stacked along the third dimension according to the arrangement of the GLM beta weights. The predicted neuronal response was derived by first multiplying the neuronal response function (with varying parameters) by the stimulus space and then integrating over the stimulus space. For each vertex of the cortical surface, we optimized the parameters of the predicted neuronal response function by minimizing the residual sum of squares relative to the GLM betas. This optimization process was performed in two steps: a grid search, which tested the performance of a large set of parameters, followed by an iterative procedure, which used the winning parameters of the grid fit as seed to explore previously untested parameter combinations. This latter step was based on the Nelder-Mead method[56] implemented in MATLAB fminsearchbnd function. The entire fitting procedure was restricted to vertices with at least one positive GLM beta, and only parameters that could explain at least 10% of the variance in the grid fit were optimized in the iterative fit. Vertices showing a negative pRF were excluded from further analyses. The above-described procedures were performed using custom-made functions in MATLAB. Supplementary figures 1 and 2 show an example of the result of the fitting procedure for each pRF model.

#### *R*^2^ cross-validation

The tested pRF models have a different number of free parameters. To overcome this issue and allow a comparison between them, we ran the pRF modeling with a 2-fold interleaved cross-validation procedure. GLM betas were estimated separately for the two halves of the dataset. In order to equalize the splits in terms of fMRI noise, for this procedure we used the denoised data generated by GLMdenoise. We ran the pRF modeling on one of the two splits. The resulting predicted neuronal responses were then compared to the GLM betas of the second split using a linear regression. The linear regression ensured that the variance explained by the pRF model was not dependent on arbitrary changes in the signal (i.e., baseline or response amplitude) between splits. The resulting cross-validated *R*^2^ values were used in the models comparison analysis (see *Neuronal response models comparison*).

#### Retinotopic mapping

We ran the standard pRF modeling[55] on the average BOLD signal of the two retinotopy runs. A two-dimensional Gaussian function was used to model neuronal responses:

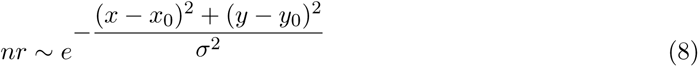

where (*x*_0_*, y*_0_) is the center of the pRF and *σ* is its size. The same apertures used to generate the retinotopy stimuli (see *Retinotopy runs*) were used to represent the *x* and *y* coordinates of the stimulus relative to the screen during the modeling procedure. The apertures matrix was convolved with the hemodynamic response function generated with the MATLAB rmHrfTwogammas function provided by Zhou and colleagues[10]. The flow of the fitting procedure was the same as described before (see *Fitting procedure*). The estimated pRF parameters were then converted into polarity and eccentricity values.

### Regions of interest (ROIs) identification

We selected two different sets of ROIs in each participant’s native space. All ROIs were selected in a blind fashion relative to the experimental functional data, to avoid circularity in the analyses.

#### Atlas-based ROIs

We identified 9 bilateral ROIs using the Destrieux atlas[57], the topological parcellation by Sereno and colleagues[58], and the HCP MMP 1.0 atlas[59]. We included areas known to be involved in either spatial[16] or temporal[12], [17] processing. We selected an ROI covering V1, V2, and V3 (Pole occipital label from Freesurfer aparc.a2009s.annot), a lateral-occipital ROI (DI, LO1, LO2, LO3 labels from Sereno’s parcellation), an occipito-parietal ROI (OPA, V3A, V3B labels from Sereno’s parcellation), an ROI covering the V5/hMT+ complex (MT upper, MT lower, MTc, MSTd, MSTv, FSTd labels from Sereno’s parcellation), an ROI covering the angular and the supramarginal gyri (PGs ROI, IP1 ROI, IP2 ROI, PFm ROI, PF ROI from HCP MMP 1.0 atlas), an ROI partially covering the superior parietal lobule and the intraparietal sulcus (V7, cIPS, LIP0, LIP1, PEc, IPS4, IPS5, aPCu1, aPCu2 labels from Sereno’s parcellation), an ROI covering the inferior frontal sulcus (DLPFC, DLPFCa, DLPFCaud labels from from Sereno’s parcellation), an ROI covering the frontal eye fields (FEF ROI, 6a ROI, i6-8 ROI labels from HPC MMP 1.0 atlas), and an ROI covering the supplementary motor area (SMA1, SMA2, dmFEF, dmFAF labels from Sereno’s parcellation and the medial part of the BA6 Freesurfer label). These ROIs were used in the models comparison analysis (see *Neuronal response models comparison*).

#### Custom-made ROIs

We drew 8 bilateral ROIs belonging to the dorsal visual stream. An initial guess of ROIs locations was provided by the topological parcellation by Sereno and colleagues[58] and by the maximum likelihood probabilistic retinotopic atlas by Wang and colleagues[60] projected onto the subjects’ native cortical surface. Afterwards, we modified ROI boundaries guided by eccentricity and polar angle maps derived from the retinotopic modeling (see *Retinotopic mapping*), and the eccentricity progression was considered the leading feature to draw ROIs contours. We selected 3 occipital ROIs (dorsal V1 and 2d, V3A and V3B, V5/MT), 4 parietal ROIs covering the intraparietal sulcus (IPS0, IPS1, IPS2, IPS3), and 1 frontal ROI covering the frontal eye fields (FEFs). Within each ROI we also drew four borders to better outline eccentricity maps: two borders marked the low and the high side of the eccentricity progression (named low and high borders); the other two were conjunction borders (named lateral borders). In some ROIs the eccentricity map was better captured by two high borders instead of a low and a high border. To place borders, two main criteria were applied[35]: map continuity (i.e., map’s vertices should belong to the same spatial cluster) and progression continuity (i.e., map’s vertices should be arranged following one eccentricity gradient only). These ROIs were used to investigate the properties of BOLD responses when position and duration co-vary in the stimulus (see *Properties of GST model*) and to study spatial and duration topographies (see *Spatial relationship between eccentricity and duration maps* and *Spatial properties of eccentricity and duration maps*). Only vertices within the borders were considered in the analyses.

#### Neuronal response models comparison

With this analysis, we aimed to investigate if and how neuronal responses to stimulus durations are linked to those of spatial location along the cortical hierarchy. To this purpose, we compared the goodness of fit of the CMT, CMTS, GS, and GT models. For each participant, we performed a vertex-wise winner-take-all procedure on models’ cross-validated *R*^2^ values. This procedure enabled us to identify for each vertex its winning model (i.e., the model with the higher *R*^2^), obtaining a winning models’ distribution for each participant. Figure 2a shows, for illustrative purposes, the group-level winning models’ distribution. We obtained it by first resampling individual distributions on a common surface (FreeSurfer fsaverage) using FreeSurfer’s mri surf2surf and then computing for each vertex the mode across participants. We excluded from the final distribution vertices that had more than one associated winning model, vertices with non-integer values due to the surface resampling, and vertices without winning model assignment in at least 7 subjects out of 13. To quantify the distribution of winning models at group level, for each participant and each ROI (see *Atlas-based ROIs*), we computed the fraction of vertices assigned to each model. These data were analyzed using a three-way repeated measure ANOVA with model type, ROI, and hemisphere as factors. The ANOVA was performed with the anova test function in R (rstatix package[61]). Marginal means were estimated with the lm function in R (model: *fraction of vertices ∼ ROI ∗Hemi∗ModelType*) and compared using the emmeans package[62] which uses the Kenward-Roger’s method[63] to estimate degrees of freedom. We did not include the GST model in this analysis because, given its inherent generality, it encompasses all the other models. Therefore, it would prevail over them in terms of goodness of fit, preventing the emergence of the actual pattern of population response functions.

### Properties of GST model

In this set of analyses, we studied the parameter of the GST model to characterize the properties of BOLD responses when position and duration co-vary in the stimulus. We focused within 8 bilateral ROIs belonging to the dorsal visual stream and defined by means of their eccentricity maps (see *Custom-made ROIs*).

#### pRFs duration preference

To assess how the distribution of the *µ_d_* parameter (i.e., preferred duration) changed across ROIs, for each participant we computed the average *µ_d_* within each ROI and we tested the following LME model:

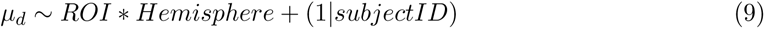

We applied the same methodology described in *Behavioral performance analysis*, and the estimated marginal means were compared using the emmeans package.

#### pRFs eccentricity preference

We performed a correlation analysis between eccentricity preferences estimated on the retinotopic data and those estimated by the GST model (i.e., *µ_s_* converted into eccentricity values). The vertex-wide Kendall’s correlation coefficient between the two sets of eccentricity was computed for each participant and ROI and then transformed into z-score to be entered in a LME model. The model formula was specified as follows:

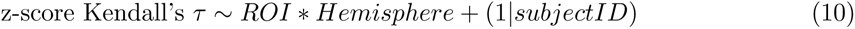

We used the same procedure described previously (see *Behavioral performance analysis* and *pRFs duration preference*).

#### pRFs orientation

We tested whether the distribution of the *θ* parameter (i.e., the orientation of the pRF) changed across ROIs using the Fisher’s non-parametric test for a common median direction[64]. This test was implemented with custom functions in R, based on the description provided by Pewsey and colleagues[65], and subsequently validated using the CircStat toolbox in MATLAB[66]. The test was performed by comparing individual median *θ* values across ROIs. Medians were calculated using the circular package in R. P values were estimated using a random permutation test, in which the test statistics was computed 9999 times on data with shuffled labels. The resulting p value was computed as the probability of finding a test statistics in the random permutations greater than in the unshuffled data.

#### pRFs aspect ratio

We calculated the aspect ratio of each vertex as 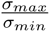 and we tested the log-transformed median aspect ratio of each ROI and participant with the following LME model:

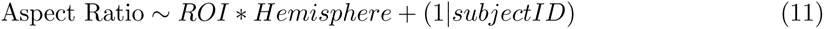

We used the same methodology described in *Behavioral performance analysis* and in *pRFs duration preference*. Aspect ratios greater than 50 were excluded from the analysis. The log transformation was used to improve normality in the sample data and ensure robust model performance.

#### pRFs selectivity

We identified for each vertex which stimulus dimension (i.e., the spatial position, the duration, or both) mainly led its response. This was possible by combining the *θ* and the *σ* parameters as follows:

- space-selective responses were defined by two different conditions: *θ* parameter horizontally oriented (i.e., *θ <* 20*^◦^* or *θ >* 160*^◦^*) coupled with *σ_d_* greater than *σ_s_*, or *θ* parameter vertically oriented (i.e. 70*^◦^ < θ <* 110*^◦^*) coupled with *σ_s_* greater than *σ_d_*;
- time-selective responses are complementary to space-selective responses: *θ* parameter vertically oriented coupled with *σ_d_* greater than *σ_s_*, or *θ* parameter horizontally oriented coupled with *σ_s_*greater than *σ_d_*;
- responses selective to both space and time were defined by a *θ* parameter falling within the ranges 20*^◦^* - 70*^◦^* or 110*^◦^* - 160*^◦^*.

For each subject and ROI we computed the fraction of vertices assigned to each type of selectivity and we performed a three-way repeated measure ANOVA with stimulus dimension (i.e., space, time, and both), ROI, and hemisphere as factors (same methodology as described in *Neuronal response models comparison*).

### Spatial relationship between eccentricity and duration maps

We designed the following analyses to investigate the spatial relationship between eccentricity and duration maps. Specifically, we studied eccentricity preferences estimated on the retinotopy data and duration preferences (*µ_d_*) estimated with the GST model on the experimental data. We considered eccentricity maps derived from the retinotopy data - rather than those estimated by the GST model on the experimental data - to rely on spatial maps obtained through the classical stimulation paradigm. These analyses were conducted within the previously described 8 bilateral ROIs belonging to the dorsal visual stream (see *Custom-made ROIs*).

#### Spatial gradients analysis

We implemented a pipeline based on spatial gradients as a data-driven method to represent and compare maps’ unfolding along the cortical surface. For each individual ROI, we first created smoothed versions of eccentricity and duration maps. We divided the distance along the *x* and *y* axes (i.e., difference between the maximum and the minimum *x* and *y* coordinates) of each map in bins of 2 mm, creating a grid that overlaid the map space. We then computed the weighted average of preference values (either eccentricity or duration) of vertices falling within each cell of the grid. Weights were determined using a Gaussian filter with a full width half maximum of 4 mm, centered on each grid cell. We considered weights lower than 0.001 as 0. Using this smoothed version of the map, we computed the spatial gradient using the MATLAB gradient function. Figure 5 shows eccentricity (panel a) and duration (panel b) smoothed maps along with the isolines representing the spatial gradients for all the ROIs in the left hemisphere of one example participant. Finally, we summed over the *x* and *y* components of each gradient and computed the resulting vector. This single vector, which we named global gradient, represents the main direction of preference change of each map (insets in figure 5a,b). When calculating global gradients, we excluded cells in the grid with gradient magnitudes greater than 1*^◦^* for eccentricity and greater than 0.1 s for duration, as these extensive variations could be driven by noisy portions of the map. The spatial relationship between eccentricity and duration maps can be inferred from the angles between their spatial gradients (see figure 5c). In light of this, for each ROI and participant we calculated the angles between eccentricity and duration spatial gradients (*α*), and between eccentricity and duration global gradients (*α_g_*, highlighted in figure 5c). Any statistical test on the distributions of *α* values was not feasible, given the multimodality of these circular distributions (see supplementary figure 4). For this reason, we only focused on the distributions of *α_g_* values, which were analyzed using the Fisher’s non-parametric test (as described in *pRFs orientation*) to detect any changes across ROIs.

#### Bivariate Moran’s I statistics

The bivariate Moran’s I statistics[67], [68] quantifies the overall spatial correlation between two variables[69], which in our case were eccentricity and duration preferences. In brief, this statistics computes, within each individual ROI, the correlation between the eccentricity preference of each vertex and the averaged duration preference of its neighbors. The result is a single value that indicates the degree and the nature of the clustering between the two types of preference. Preference values were transformed in z-score for each ROI independently. We included in the analysis only vertices showing both eccentricity and duration preferences, estimated respectively on the retinotopy data and on the experimental data fitted with the GST model. We employed a neighborhood structure comprising the first 12 nearest neighbors of each vertex. P values were estimated through a random permutation test, in which the statistics was computed 999 times shuffling eccentricity preferences over vertices. Since each individual ROI has a unique neighborhood structure, the statistical comparison of bivariate Moran’s I values across them was not feasible. We implemented this entire procedure with custom functions in MATLAB based on and validated with the GeoDa software[70].

### Spatial properties of eccentricity and duration maps

These analyses aimed to characterize the spatial properties of duration maps as compared to eccentricity maps. As before, we used duration preferences estimated by the GST model on the experimental data and the eccentricity preferences estimated on the retinotopy data, and we focused on the 8 bilateral visual ROIs previously described (see *Custom-made ROIs*).

#### Univariate Moran’s I statistics

The univariate Moran’s I statistics quantifies the global spatial autocorrelation within a sample. In our case, it measures within each eccentricity and duration map the correlation between the preference at each vertex and the averaged preference of its neighbors, providing a single value that indicates the degree and the nature of preferences’ clustering. The statistics was computed on preference values transformed in z-score for each ROI independently, and a neighborhood structure of 12 vertices was employed. This statistical analysis was implemented as previously described for the bivariate version (see *Bivariate Moran’s I statistics*).

#### Variogram

The experimental variogram is a graphical representation of the spatial autocorrelation in a sample based on variance estimation among points located at different distances. For each individual ROI, we computed the distance between each vertex in a map (either eccentricity or duration) and then grouped the distance data into 30 bins. To build the variograms, we computed the variance between preferences of vertices belonging to the same bin. This procedure was performed with custom functions implemented in MATLAB based on the GeostatsPy package[71]. The variograms were used to subsequently extract two parameters: the nugget (i.e., the variance at the minimum distance between points), and the range (i.e., the distance required to reach the sample variance). These are respectively a proxy for the strength and for the extent of the spatial autocorrelation within the ROI. The ranges were then divided by the maximum distance between vertices in the sample to normalize these values with respect to the full extent of the map. We computed these parameters for the eccentricity and duration maps of all participants and ROIs. These data were then tested with the following LME models and an identical procedure as previously described (see *Behavioral performance analysis* and *pRFs duration preference*):

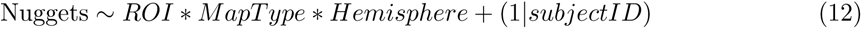

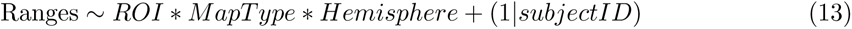

where *MapType* identifies eccentricity or duration maps.

## Acknowledgments

This project has received funding from the European Research Council (ERC) under the European Union’s Horizon 2020 research and innovation programme grant agreement no. 682117 BIT-ERC-2015-CoG and from the Italian Ministry of University and Research under the call FARE (project ID: R16X32NALR) and under the call PRIN2017 (project ID: XBJN4F) to D.B.

