## supplementary_materials for "The link between space and time along the human cortical hierarchy"

#### Supplementary Figures

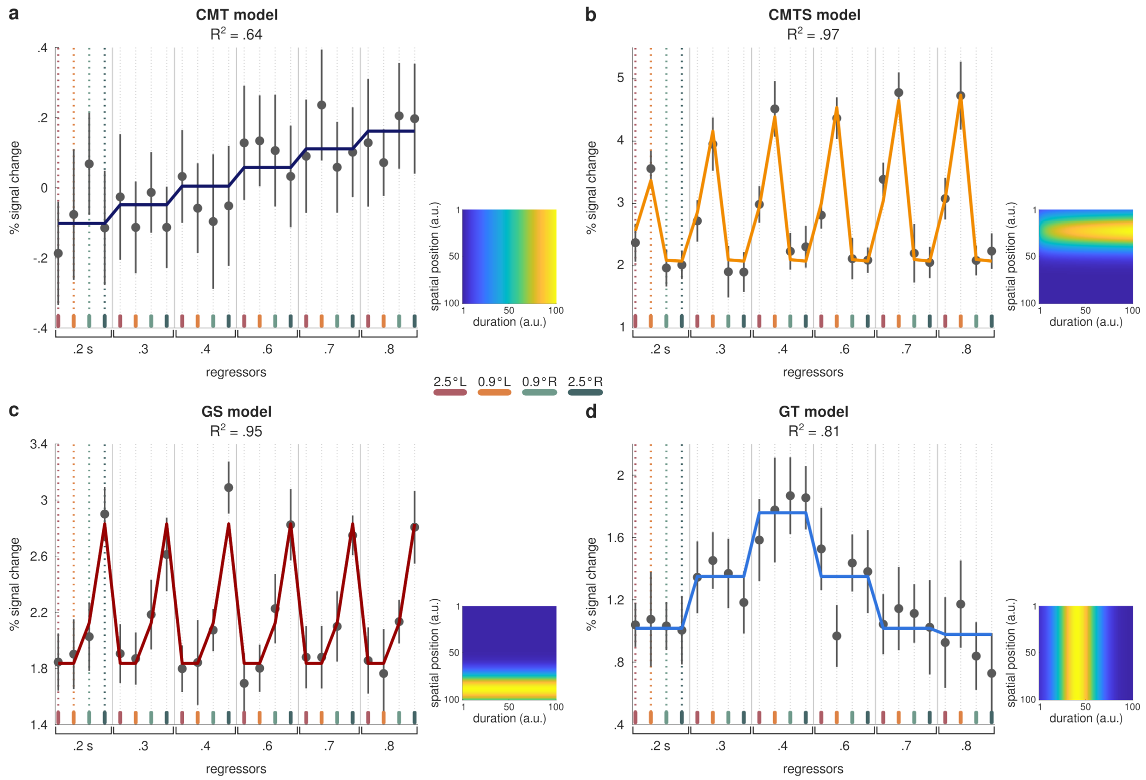


Supplementary figure 1: **pRF models.** Results of the pRF modeling for the CMT **(a)**, CMTS **(b)**, GS **(c)** and GT **(d)** models are shown in four example vertices (the goodness of fit is reported in each panel). Solid lines represent the predicted response color-coded according to the different pRF models. Black dots represent the median GLM beta weights converted to percentage of signal change (see *General Linear Model (GLM) analysis* in *Materials and Methods*) relative to each combination of stimulus duration and spatial position (color-coded). Error bars represent the standard errors of the median. The insets in each panel represent the population response function in the stimuli space. See *Population Receptive Field (pRF) modeling* in *Materials and Methods*.

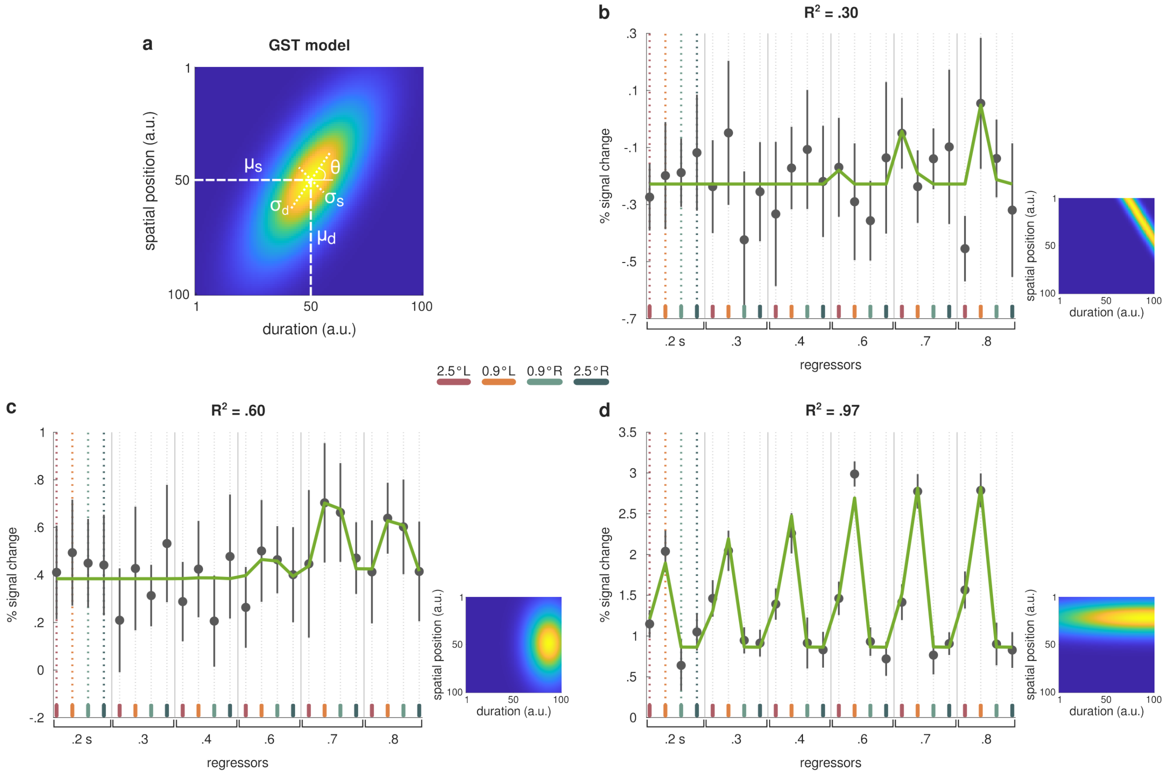


Supplementary figure 2: **GST model.** **(a)** General form of the bivariate population response function, where its preference ($\mu_d$ and $\mu_s$), sensitivity ($\sigma_d$ and $\sigma_s$), and orientation ($\theta$) are displayed. **(b)**, **(c)**, **(d)** Results of the pRF modeling are shown in three example vertices exhibiting different response behaviors captured by the GST model (the goodness of fit is reported in each panel). As in supplementary figure 1, solid lines represent the predicted response, black dots represent the median GLM beta weights converted to percentage of signal change relative to each combination of stimulus duration and spatial position (color-coded), and error bars represent the standard errors of the median. The insets in each panel represent the population response function in the stimuli space. See *Population Receptive Field (pRF) modeling* in *Materials and Methods*.

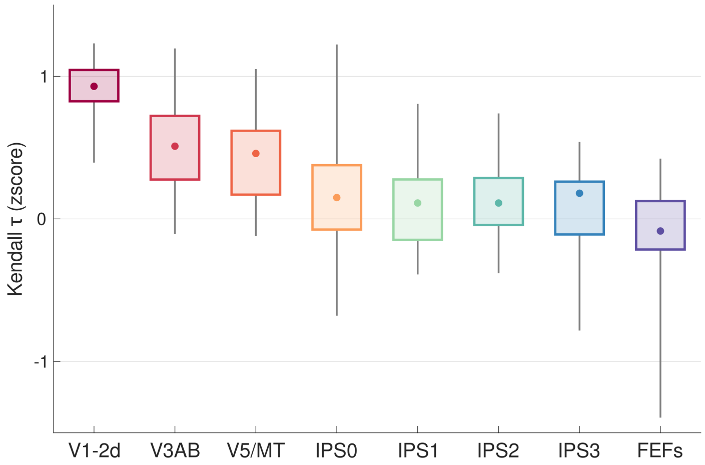


Supplementary figure 3: **Correlation between eccentricity preferences.** Boxplots represent for each ROI (color-coded) the distribution across participants and hemispheres of the vertex-wide Kendall's $\tau$ correlation coefficient (z-score transformed) between eccentricity preferences estimated by the GTS model ($\mu_s$) and eccentricity preferences estimated on the retinotopic data. The boxes represent the interquartile range of each distribution, the extremes of the whiskers are its maximum and minimum. Dots represent the median of the distributions. See *pRFs eccentricity preference* in *Materials and Methods - Properties of GST model*. ROI legend: V1-2d = dorsal primary and secondary visual areas, V3AB = visual areas V3A and V3B, V5/MT = visual area V5/MT, IPS0-3 = different portions of the intraparietal sulcus, FEFs = frontal eye fields.

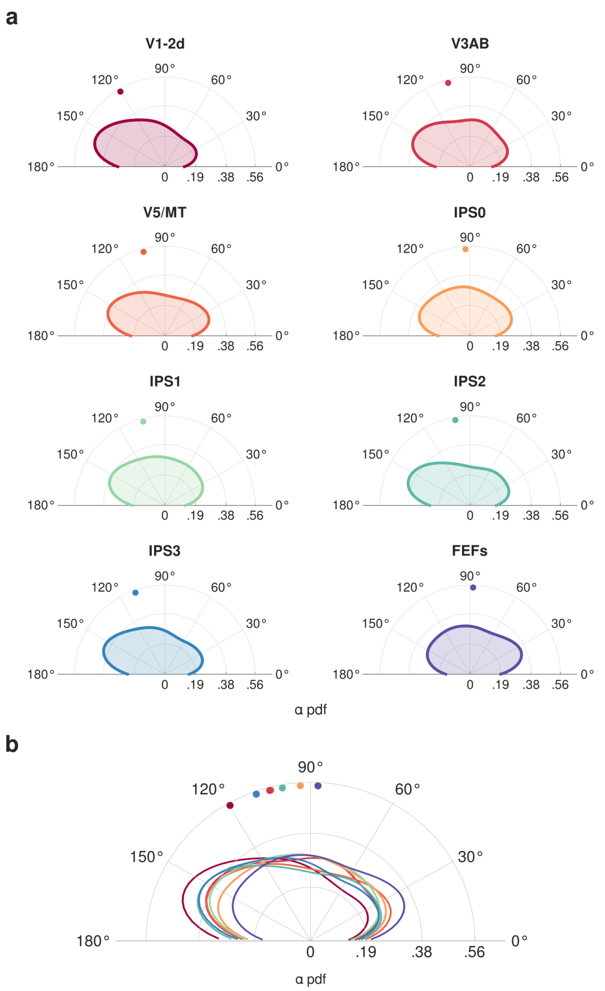


Supplementary figure 4: **Distributions of angles between duration and eccentricity gradient vector fields ($\alpha$). (a)** Polar plots show for each ROI the group-level distribution of angles ($\alpha$) computed between duration and eccentricity gradient vector fields. Dots represent the median of the distributions: V1-2d = 121°, V3AB = 105°, V5/MT = 105°, IPS0 = 93°, IPS1 = 105°, IPS2 = 100°, IPS3 = 110°, FEFs = 88°. Distributions' kernels were estimated with 15° bandwidth. We excluded from $\alpha$ computation spatial gradients with a magnitude smaller than 0.1° for eccentricity and smaller than 0.02 s for duration, in order to ignore negligible changes that could result in spurious angles. **(b)** Distributions' kernels for each ROI (color-coded as in panel a) are superimposed in the same polar plot. Dots represent the median of each distribution. See *Spatial gradients analysis* in *Materials and Methods - Spatial relationship between eccentricity and duration maps*. ROI legend: V1-2d = dorsal primary and secondary visual areas, V3AB = visual areas V3A and V3B, V5/MT = visual area V5/MT, IPS0-3 = different portions of the intraparietal sulcus, FEFs = frontal eye fields.

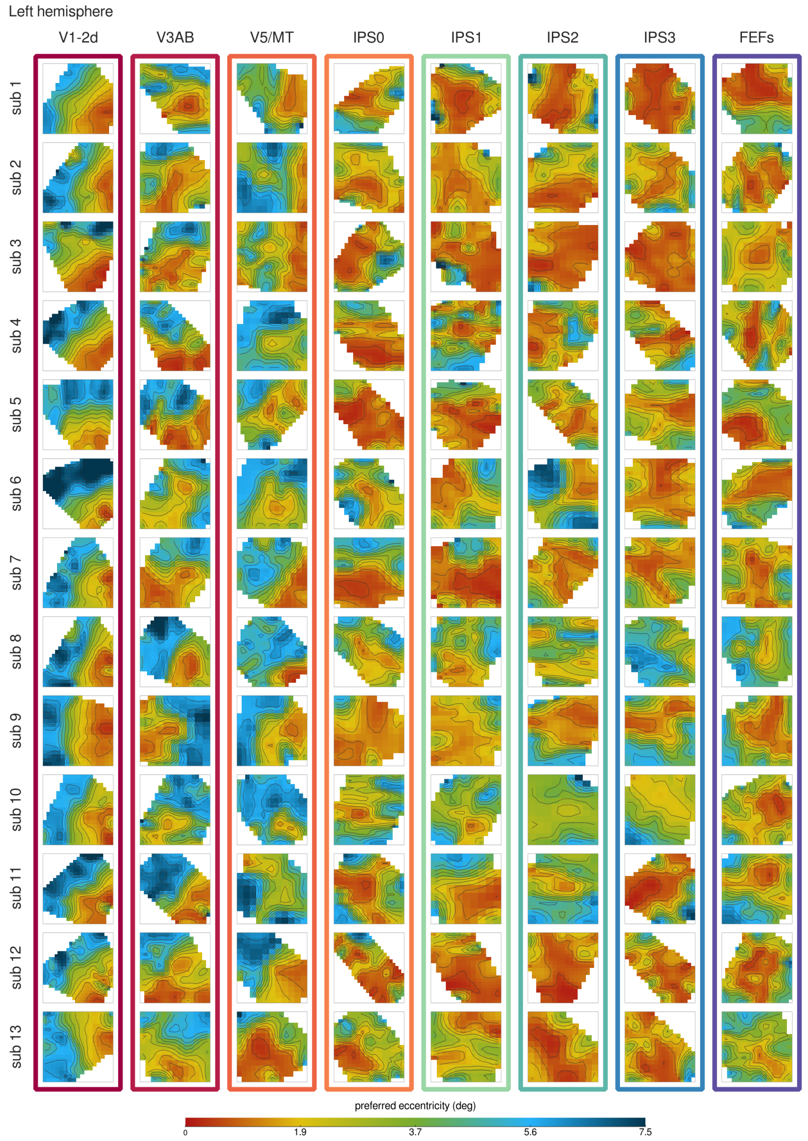


Supplementary figure 5: **Eccentricity spatial gradients.** Eccentricity maps of all participants (in the different rows) are shown for all the ROIs in the **left** hemisphere (in the different columns). The isolines, representing the spatial gradients, are displayed in black, and they are separated by 0.5° change of eccentricity preference. Preferred eccentricities are color-coded from red (0°) to dark blue (7.5°). Maps in the same ROI across participants are delineated by a color-coded outline for easy identification. See *Spatial gradients analysis* in *Materials and Methods - Spatial relationship between eccentricity and duration maps*. ROI legend: V1-2d = dorsal primary and secondary visual areas, V3AB = visual areas V3A and V3B, V5/MT = visual area V5/MT, IPS0-3 = different portions of the intraparietal sulcus, FEFs = frontal eye fields.

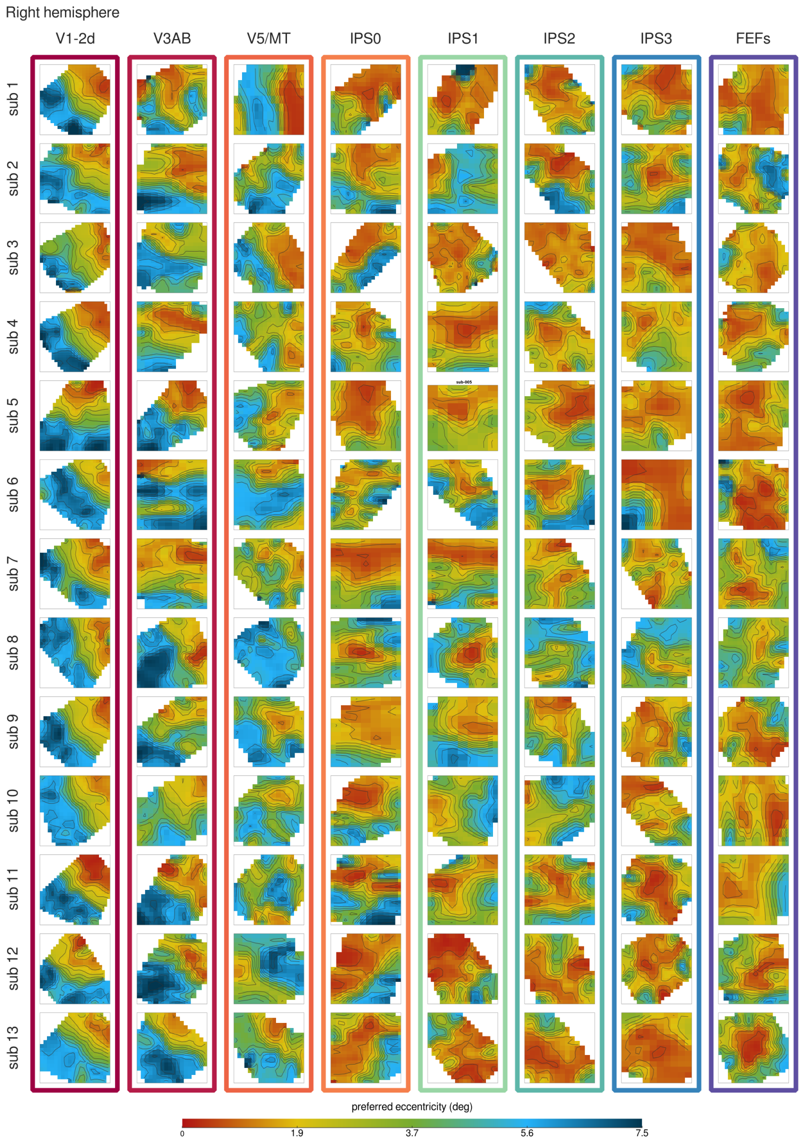


Supplementary figure 6: **Eccentricity spatial gradients.** Eccentricity maps of all participants (in the different rows) are shown for all the ROIs in the **right** hemisphere (in the different columns). The isolines, representing the spatial gradients, are displayed in black, and they are separated by 0.5° change of eccentricity preference. Preferred eccentricities are color-coded from red (0°) to dark blue (7.5°). Maps in the same ROI across participants are delineated by a color-coded outline for easy identification. See *Spatial gradients analysis* in *Materials and Methods - Spatial relationship between eccentricity and duration maps.* ROI legend: V1-2d = dorsal primary and secondary visual areas, V3AB = visual areas V3A and V3B, V5/MT = visual area V5/MT, IPS0-3 = different portions of the intraparietal sulcus, FEFs = frontal eye fields.

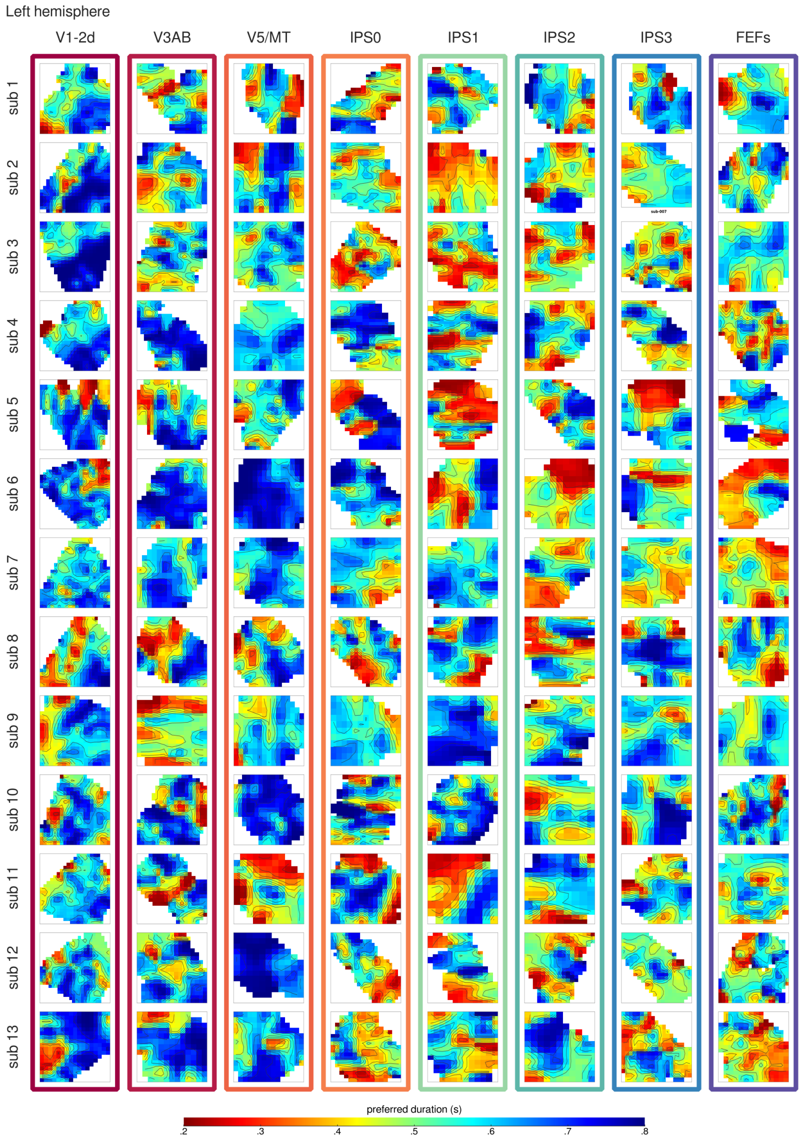


Supplementary figure 7: **Duration spatial gradients.** Duration maps of all participants (in the different rows) are shown for all the ROIs in the **left** hemisphere (in the different columns). The isolines, representing the spatial gradients, are displayed in black, and they are separated by 50 ms change of duration preference. Preferred durations are color-coded from red (.2 s) to blue (.8 s). Maps in the same ROI across participants are delineated by a color-coded outline for easy identification. See *Spatial gradients analysis* in *Materials and Methods - Spatial relationship between eccentricity and duration maps*. ROI legend: V1-2d = dorsal primary and secondary visual areas, V3AB = visual areas V3A and V3B, V5/MT = visual area V5/MT, IPS0-3 = different portions of the intraparietal sulcus, FEFs = frontal eye fields.

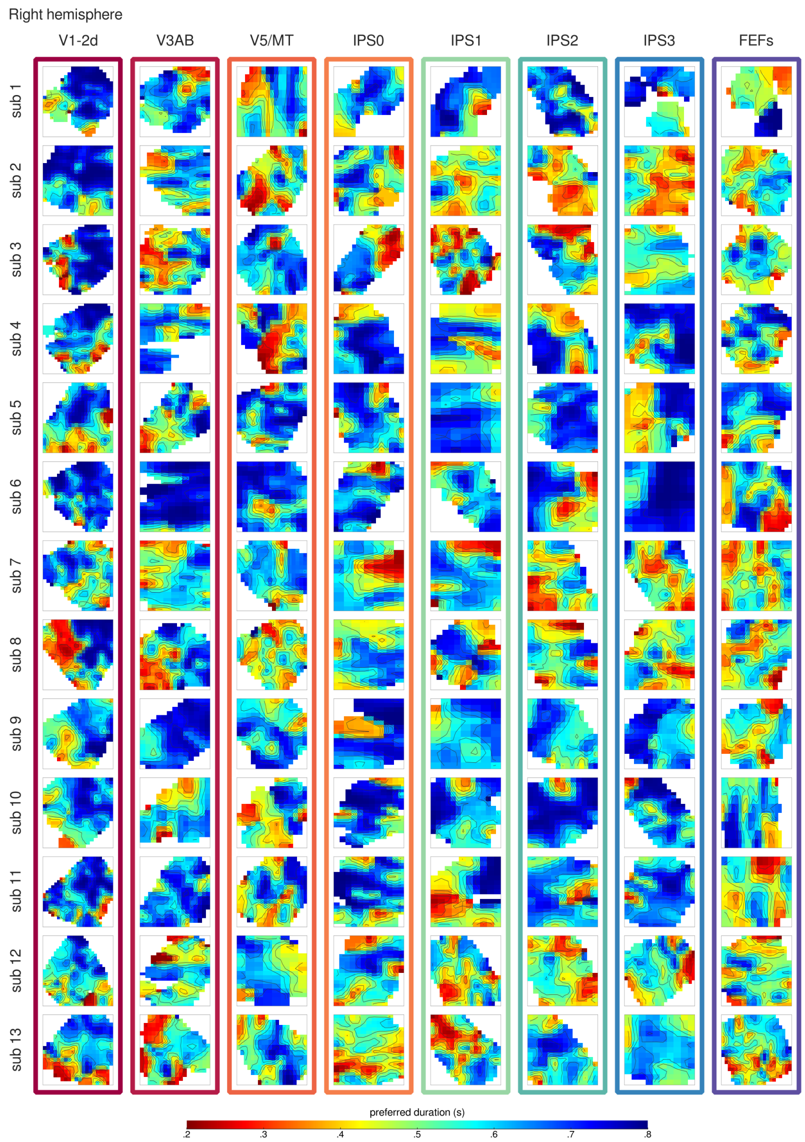


Supplementary figure 8: **Duration spatial gradients.** Duration maps of all participants (in the different rows) are shown for all the ROIs in the **right** hemisphere (in the different columns). The isolines, representing the spatial gradients, are displayed in black, and they are separated by 50 ms change of duration preference. Preferred durations are color-coded from red (.2 s) to blue (.8 s). Maps in the same ROI across participants are delineated by a color-coded outline for easy identification. See *Spatial gradients analysis* in *Materials and Methods - Spatial relationship between eccentricity and duration maps*. ROI legend: V1-2d = dorsal primary and secondary visual areas, V3AB = visual areas V3A and V3B, V5/MT = visual area V5/MT, IPS0-3 = different portions of the intraparietal sulcus, FEFs = frontal eye fields.

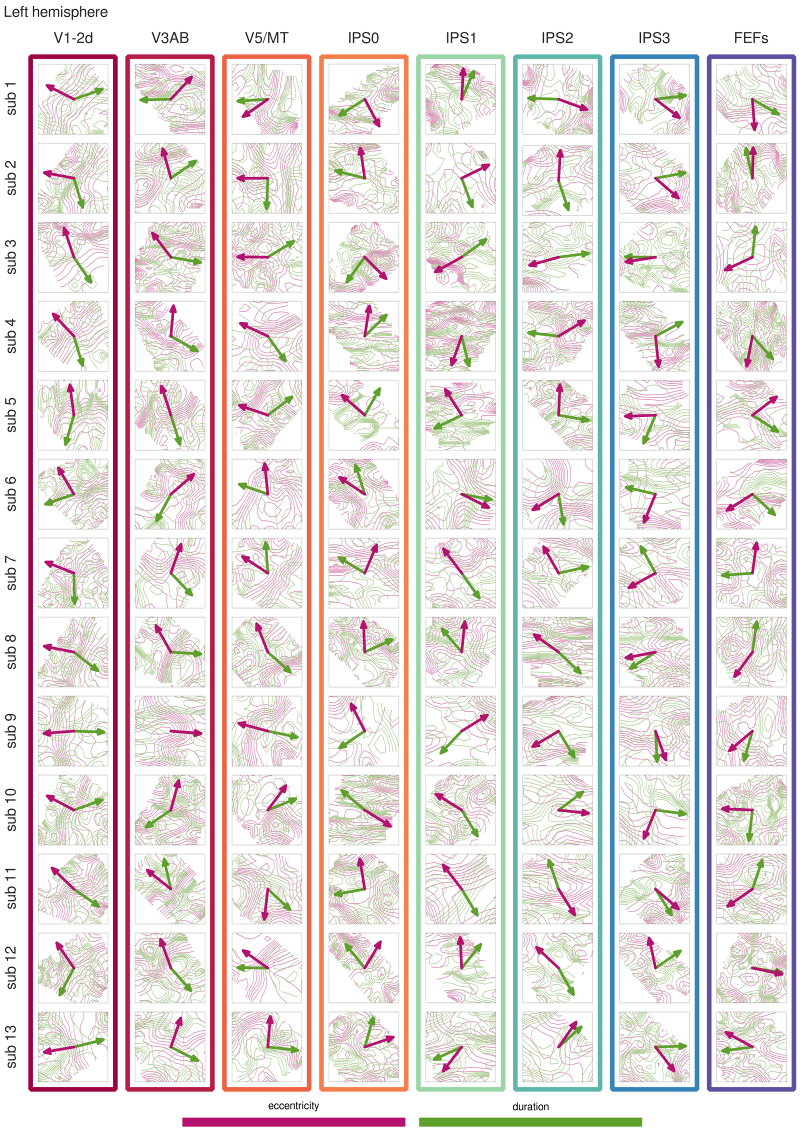


Supplementary figure 9: **Relationship between eccentricity and duration spatial gradients.** Eccentricity (in pink) and duration (in green) isolines of all participants (in the different rows) are displayed overlaid for each ROI in the **left** hemisphere (in the different columns). Eccentricity (pink arrow) and duration (green arrow) global gradients are plotted together to highlight the angle ($\alpha_g$) between them. The $\alpha_g$ served a measure of the spatial relationship between eccentricity and duration maps. Vectors were normalized (by dividing the vector by its magnitude) for easy visualization. Spatial gradients in the same ROI across participants are delineated by a color-coded outline for easy identification. See *Spatial gradients analysis* in *Materials and Methods - Spatial relationship between eccentricity and duration maps*. ROI legend: V1-2d = dorsal primary and secondary visual areas, V3AB = visual areas V3A and V3B, V5/MT = visual area V5/MT, IPS0-3 = different portions of the intraparietal sulcus, FEFs = frontal eye fields.

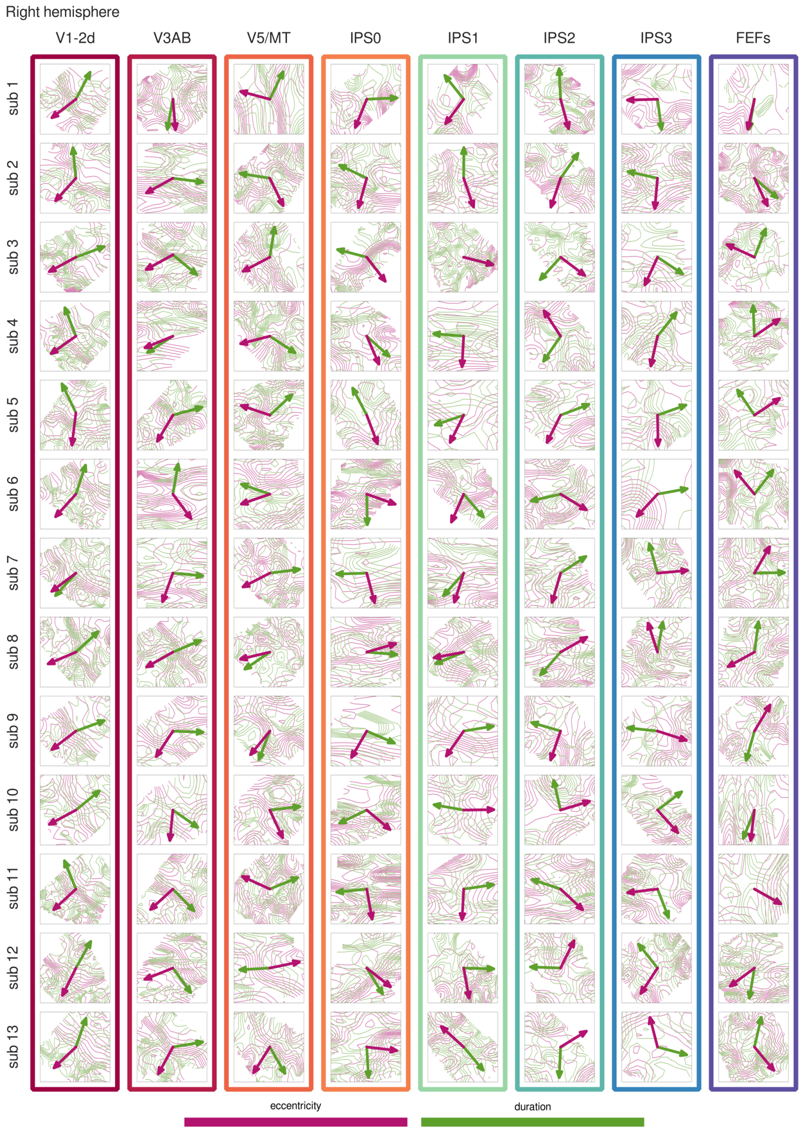


Supplementary figure 10: **Relationship between eccentricity and duration spatial gradients.** Eccentricity (in pink) and duration (in green) isolines of all participants (in the different rows) are displayed overlaid for each ROI in the **right** hemisphere (in the different columns). Eccentricity (pink arrow) and duration (green arrow) global gradients are plotted together to highlight the angle ($\alpha_g$) between them. The $\alpha_g$ served a measure of the spatial relationship between eccentricity and duration maps. Vectors were normalized (by dividing the vector by its magnitude) for easy visualization. Spatial gradients in the same ROI across participants are delineated by a color-coded outline for easy identification. See *Spatial gradients analysis* in *Materials and Methods - Spatial relationship between eccentricity and duration maps*. ROI legend: V1-2d = dorsal primary and secondary visual areas, V3AB = visual areas V3A and V3B, V5/MT = visual area V5/MT, IPS0-3 = different portions of the intraparietal sulcus, FEFs = frontal eye fields.

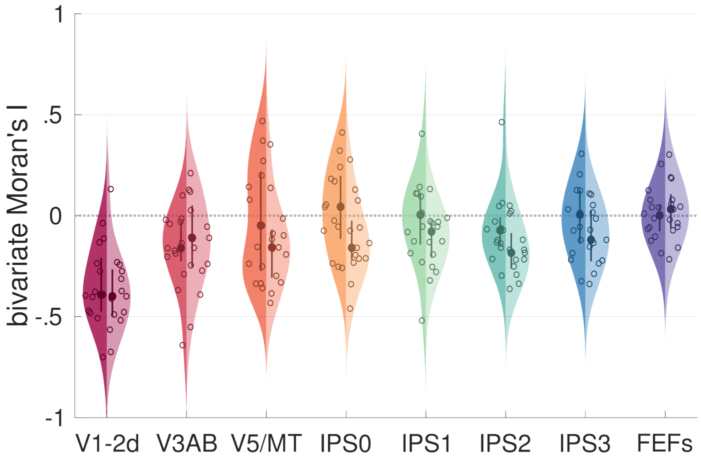


Supplementary figure 11: **Group-level distributions of the bivariate Moran's I across ROIs.** Each violin plot represents the group-level distribution of the bivariate Moran's I in a different ROI. The left side of the distributions (darker shade) refers to the left hemisphere, while the right side (lighter shade) refers to the right hemisphere. Dots indicate the median of the distributions, and circles correspond to individual data points. Thick lines represent the interquartile range of the distributions, the kernels were estimated with 15% bandwidth. The bivariate Moran's I measured the degree of spatial correlation between eccentricity preferences (estimated on the retinotopic data) and duration preferences ($\mu_d$ estimated by the GST model). A bivariate Moran's I close to 0 indicates spatial randomness between the two types of preference, whereas values higher or lower than 0 indicate spatial clustering, characterized by positive or negative correlation respectively. See *Bivariate Moran's I statistics* in *Materials and Methods - Spatial relationship between eccentricity and duration maps*. ROI legend: V1-2d = dorsal primary and secondary visual areas, V3AB = visual areas V3A and V3B, V5/MT = visual area V5/MT, IPS0-3 = different portions of the intraparietal sulcus, FEFs = frontal eye fields.

### Supplementary Tables

| | Sum.Sq | Mean.Sq | NumDF | DenDF | F.value | Pr(>F) |
|---|---|---|---|---|---|---|
| Stimulus Position | 0.00 | 0.00 | 3 | 36.00 | 1.16 | 0.34 |

Supplementary table 1: PSE - type III ANOVA on LME model estimates with Satterthwaite's method for degrees of freedom

| | Sum.Sq | Mean.Sq | NumDF | DenDF | F.value | Pr(>F) |
|---|---|---|---|---|---|---|
| Stimulus Position | 0.00 | 0.00 | 3 | 36.00 | 0.83 | 0.49 |

Supplementary table 2: JND - type III ANOVA on LME model estimates with Satterthwaite's method for degrees of freedom

| | DFn | DFd | F | p | p<.05 | ges |
|---|---|---|---|---|---|---|
| ROI | 8 | 96 | -0.00 | 1.00 | | 0.00 |
| Hemi | 1 | 12 | -0.00 | 1.00 | | 0.00 |
| ModelType | 3 | 36 | 24.25 | 0.00 | * | 0.24 |
| ROI:Hemi | 8 | 96 | -0.00 | 1.00 | | 0.00 |
| ROI:ModelType | 24 | 288 | 33.16 | 0.00 | * | 0.58 |
| Hemi:ModelType | 3 | 36 | 1.60 | 0.21 | | 0.01 |
| ROI:Hemi:ModelType | 24 | 288 | 0.95 | 0.53 | | 0.02 |

Supplementary table 3: CMT, CMTS, GS, and GT models comparison - three-way repeated measure ANOVA

| | W | p | p<.05 |
|---|---|---|---|
| ModelType | 0.33 | 0.04 | * |
| Hemi:ModelType | 0.18 | 0.00 | * |

Supplementary table 4: CMT, CMTS, GS, and GT models comparison - Mauchly's sphericity test

| | GGe | DF[GG] | p[GG] | p[GG]<.05 | HFe | DF[HF] | p[HF] | p[HF]<.05 |
|---|---|---|---|---|---|---|---|---|
| ModelType | 0.57 | 1.71, 20.47 | 0.00 | * | 0.65 | 1.96, 23.51 | 0.00 | * |
| Hemi:ModelType | 0.49 | 1.47, 17.63 | 0.23 | | 0.54 | 1.62, 19.49 | 0.23 | |

Supplementary table 5: CMT, CMTS, GS, and GT models comparison - deviation from sphericity corrections

| | ModelType | estimate | SE | df | t.ratio | p.value |
|---|---|---|---|---|---|---|
| (AnG-SmG) - FEFs | CMT | 0.01 | 0.02 | 864 | 0.46 | 1.00 |
| (AnG-SmG) - (IFS-PCSi) | CMT | 0.02 | 0.02 | 864 | 1.16 | 1.00 |
| (AnG-SmG) - LO | CMT | 0.15 | 0.02 | 864 | 7.63 | 0.00 |
| (AnG-SmG) - OcP | CMT | 0.15 | 0.02 | 864 | 7.63 | 0.00 |
| (AnG-SmG) - SMA | CMT | -0.04 | 0.02 | 864 | -1.92 | 1.00 |
| (AnG-SmG) - (IPS-SPL) | CMT | 0.03 | 0.02 | 864 | 1.74 | 1.00 |
| (AnG-SmG) - V3AB | CMT | 0.07 | 0.02 | 864 | 3.55 | 0.01 |
| (AnG-SmG) - (V5/MT) | CMT | 0.03 | 0.02 | 864 | 1.63 | 1.00 |
| FEFs - (IFS-PCSi) | CMT | 0.01 | 0.02 | 864 | 0.70 | 1.00 |
| FEFs - LO | CMT | 0.14 | 0.02 | 864 | 7.17 | 0.00 |
| FEFs - OcP | CMT | 0.14 | 0.02 | 864 | 7.17 | 0.00 |
| FEFs - SMA | CMT | -0.05 | 0.02 | 864 | -2.38 | 0.64 |
| FEFs - (IPS-SPL) | CMT | 0.03 | 0.02 | 864 | 1.28 | 1.00 |
| FEFs - V3AB | CMT | 0.06 | 0.02 | 864 | 3.09 | 0.07 |
| FEFs - (V5/MT) | CMT | 0.02 | 0.02 | 864 | 1.18 | 1.00 |
| (IFS-PCSi) - LO | CMT | 0.13 | 0.02 | 864 | 6.47 | 0.00 |
| (IFS-PCSi) - OcP | CMT | 0.13 | 0.02 | 864 | 6.47 | 0.00 |
| (IFS-PCSi) - SMA | CMT | -0.06 | 0.02 | 864 | -3.08 | 0.08 |
| (IFS-PCSi) - (IPS-SPL) | CMT | 0.01 | 0.02 | 864 | 0.58 | 1.00 |
| (IFS-PCSi) - V3AB | CMT | 0.05 | 0.02 | 864 | 2.39 | 0.61 |
| (IFS-PCSi) - (V5/MT) | CMT | 0.01 | 0.02 | 864 | 0.48 | 1.00 |
| LO - OcP | CMT | -0.00 | 0.02 | 864 | -0.00 | 1.00 |
| LO - SMA | CMT | -0.19 | 0.02 | 864 | -9.55 | 0.00 |
| LO - (IPS-SPL) | CMT | -0.12 | 0.02 | 864 | -5.89 | 0.00 |
| LO - V3AB | CMT | -0.08 | 0.02 | 864 | -4.08 | 0.00 |
| LO - (V5/MT) | CMT | -0.12 | 0.02 | 864 | -6.00 | 0.00 |
| OcP - SMA | CMT | -0.19 | 0.02 | 864 | -9.55 | 0.00 |
| OcP - (IPS-SPL) | CMT | -0.12 | 0.02 | 864 | -5.89 | 0.00 |
| OcP - V3AB | CMT | -0.08 | 0.02 | 864 | -4.08 | 0.00 |
| OcP - (V5/MT) | CMT | -0.12 | 0.02 | 864 | -5.99 | 0.00 |
| SMA - (IPS-SPL) | CMT | 0.07 | 0.02 | 864 | 3.66 | 0.01 |
| SMA - V3AB | CMT | 0.11 | 0.02 | 864 | 5.47 | 0.00 |
| SMA - (V5/MT) | CMT | 0.07 | 0.02 | 864 | 3.55 | 0.01 |
| (IPS-SPL) - V3AB | CMT | 0.04 | 0.02 | 864 | 1.81 | 1.00 |
| (IPS-SPL) - (V5/MT) | CMT | -0.00 | 0.02 | 864 | -0.11 | 1.00 |
| V3AB - (V5/MT) | CMT | -0.04 | 0.02 | 864 | -1.92 | 1.00 |
| (AnG-SmG) - FEFs | CMTS | 0.00 | 0.02 | 864 | 0.13 | 1.00 |
| (AnG-SmG) - (IFS-PCSi) | CMTS | 0.00 | 0.02 | 864 | 0.23 | 1.00 |
| (AnG-SmG) - LO | CMTS | -0.30 | 0.02 | 864 | -15.07 | 0.00 |
| (AnG-SmG) - OcP | CMTS | -0.29 | 0.02 | 864 | -14.40 | 0.00 |
| (AnG-SmG) - SMA | CMTS | 0.02 | 0.02 | 864 | 1.24 | 1.00 |

| | | | | | | |
|---|---|---|---|---|---|---|
| (AnG-SmG) - (IPS-SPL) | CMTS | -0.06 | 0.02 | 864 | -3.21 | 0.05 |
| (AnG-SmG) - V3AB | CMTS | -0.13 | 0.02 | 864 | -6.66 | 0.00 |
| (AnG-SmG) - (V5/MT) | CMTS | -0.08 | 0.02 | 864 | -4.17 | 0.00 |
| FEFs - (IFS-PCSi) | CMTS | 0.00 | 0.02 | 864 | 0.10 | 1.00 |
| FEFs - LO | CMTS | -0.31 | 0.02 | 864 | -15.20 | 0.00 |
| FEFs - OcP | CMTS | -0.29 | 0.02 | 864 | -14.53 | 0.00 |
| FEFs - SMA | CMTS | 0.02 | 0.02 | 864 | 1.11 | 1.00 |
| FEFs - (IPS-SPL) | CMTS | -0.07 | 0.02 | 864 | -3.35 | 0.03 |
| FEFs - V3AB | CMTS | -0.14 | 0.02 | 864 | -6.79 | 0.00 |
| FEFs - (V5/MT) | CMTS | -0.09 | 0.02 | 864 | -4.30 | 0.00 |
| (IFS-PCSi) - LO | CMTS | -0.31 | 0.02 | 864 | -15.30 | 0.00 |
| (IFS-PCSi) - OcP | CMTS | -0.29 | 0.02 | 864 | -14.63 | 0.00 |
| (IFS-PCSi) - SMA | CMTS | 0.02 | 0.02 | 864 | 1.01 | 1.00 |
| (IFS-PCSi) - (IPS-SPL) | CMTS | -0.07 | 0.02 | 864 | -3.45 | 0.02 |
| (IFS-PCSi) - V3AB | CMTS | -0.14 | 0.02 | 864 | -6.89 | 0.00 |
| (IFS-PCSi) - (V5/MT) | CMTS | -0.09 | 0.02 | 864 | -4.40 | 0.00 |
| LO - OcP | CMTS | 0.01 | 0.02 | 864 | 0.67 | 1.00 |
| LO - SMA | CMTS | 0.33 | 0.02 | 864 | 16.31 | 0.00 |
| LO - (IPS-SPL) | CMTS | 0.24 | 0.02 | 864 | 11.86 | 0.00 |
| LO - V3AB | CMTS | 0.17 | 0.02 | 864 | 8.41 | 0.00 |
| LO - (V5/MT) | CMTS | 0.22 | 0.02 | 864 | 10.90 | 0.00 |
| OcP - SMA | CMTS | 0.31 | 0.02 | 864 | 15.64 | 0.00 |
| OcP - (IPS-SPL) | CMTS | 0.22 | 0.02 | 864 | 11.18 | 0.00 |
| OcP - V3AB | CMTS | 0.16 | 0.02 | 864 | 7.74 | 0.00 |
| OcP - (V5/MT) | CMTS | 0.21 | 0.02 | 864 | 10.23 | 0.00 |
| SMA - (IPS-SPL) | CMTS | -0.09 | 0.02 | 864 | -4.45 | 0.00 |
| SMA - V3AB | CMTS | -0.16 | 0.02 | 864 | -7.90 | 0.00 |
| SMA - (V5/MT) | CMTS | -0.11 | 0.02 | 864 | -5.41 | 0.00 |
| (IPS-SPL) - V3AB | CMTS | -0.07 | 0.02 | 864 | -3.45 | 0.02 |
| (IPS-SPL) - (V5/MT) | CMTS | -0.02 | 0.02 | 864 | -0.95 | 1.00 |
| V3AB - (V5/MT) | CMTS | 0.05 | 0.02 | 864 | 2.49 | 0.46 |
| (AnG-SmG) - FEFs | GS | 0.02 | 0.02 | 864 | 0.87 | 1.00 |
| (AnG-SmG) - (IFS-PCSi) | GS | 0.02 | 0.02 | 864 | 0.97 | 1.00 |
| (AnG-SmG) - LO | GS | -0.07 | 0.02 | 864 | -3.25 | 0.04 |
| (AnG-SmG) - OcP | GS | -0.05 | 0.02 | 864 | -2.41 | 0.59 |
| (AnG-SmG) - SMA | GS | 0.04 | 0.02 | 864 | 2.07 | 1.00 |
| (AnG-SmG) - (IPS-SPL) | GS | -0.02 | 0.02 | 864 | -1.04 | 1.00 |
| (AnG-SmG) - V3AB | GS | -0.03 | 0.02 | 864 | -1.58 | 1.00 |
| (AnG-SmG) - (V5/MT) | GS | -0.01 | 0.02 | 864 | -0.68 | 1.00 |
| FEFs - (IFS-PCSi) | GS | 0.00 | 0.02 | 864 | 0.10 | 1.00 |
| FEFs - LO | GS | -0.08 | 0.02 | 864 | -4.12 | 0.00 |
| FEFs - OcP | GS | -0.07 | 0.02 | 864 | -3.28 | 0.04 |

| | | | | | | |
|---|---|---|---|---|---|---|
| FEFs - SMA | GS | 0.02 | 0.02 | 864 | 1.20 | 1.00 |
| FEFs - (IPS-SPL) | GS | -0.04 | 0.02 | 864 | -1.90 | 1.00 |
| FEFs - V3AB | GS | -0.05 | 0.02 | 864 | -2.45 | 0.52 |
| FEFs - (V5/MT) | GS | -0.03 | 0.02 | 864 | -1.55 | 1.00 |
| (IFS-PCSi) - LO | GS | -0.08 | 0.02 | 864 | -4.22 | 0.00 |
| (IFS-PCSi) - OcP | GS | -0.07 | 0.02 | 864 | -3.38 | 0.03 |
| (IFS-PCSi) - SMA | GS | 0.02 | 0.02 | 864 | 1.10 | 1.00 |
| (IFS-PCSi) - (IPS-SPL) | GS | -0.04 | 0.02 | 864 | -2.01 | 1.00 |
| (IFS-PCSi) - V3AB | GS | -0.05 | 0.02 | 864 | -2.55 | 0.39 |
| (IFS-PCSi) - (V5/MT) | GS | -0.03 | 0.02 | 864 | -1.65 | 1.00 |
| LO - OcP | GS | 0.02 | 0.02 | 864 | 0.85 | 1.00 |
| LO - SMA | GS | 0.11 | 0.02 | 864 | 5.33 | 0.00 |
| LO - (IPS-SPL) | GS | 0.04 | 0.02 | 864 | 2.22 | 0.97 |
| LO - V3AB | GS | 0.03 | 0.02 | 864 | 1.67 | 1.00 |
| LO - (V5/MT) | GS | 0.05 | 0.02 | 864 | 2.58 | 0.37 |
| OcP - SMA | GS | 0.09 | 0.02 | 864 | 4.48 | 0.00 |
| OcP - (IPS-SPL) | GS | 0.03 | 0.02 | 864 | 1.37 | 1.00 |
| OcP - V3AB | GS | 0.02 | 0.02 | 864 | 0.82 | 1.00 |
| OcP - (V5/MT) | GS | 0.03 | 0.02 | 864 | 1.73 | 1.00 |
| SMA - (IPS-SPL) | GS | -0.06 | 0.02 | 864 | -3.11 | 0.07 |
| SMA - V3AB | GS | -0.07 | 0.02 | 864 | -3.66 | 0.01 |
| SMA - (V5/MT) | GS | -0.06 | 0.02 | 864 | -2.75 | 0.22 |
| (IPS-SPL) - V3AB | GS | -0.01 | 0.02 | 864 | -0.55 | 1.00 |
| (IPS-SPL) - (V5/MT) | GS | 0.01 | 0.02 | 864 | 0.36 | 1.00 |
| V3AB - (V5/MT) | GS | 0.02 | 0.02 | 864 | 0.91 | 1.00 |
| (AnG-SmG) - FEFs | GT | -0.03 | 0.02 | 864 | -1.46 | 1.00 |
| (AnG-SmG) - (IFS-PCSi) | GT | -0.05 | 0.02 | 864 | -2.36 | 0.66 |
| (AnG-SmG) - LO | GT | 0.21 | 0.02 | 864 | 10.69 | 0.00 |
| (AnG-SmG) - OcP | GT | 0.18 | 0.02 | 864 | 9.18 | 0.00 |
| (AnG-SmG) - SMA | GT | -0.03 | 0.02 | 864 | -1.39 | 1.00 |
| (AnG-SmG) - (IPS-SPL) | GT | 0.05 | 0.02 | 864 | 2.51 | 0.44 |
| (AnG-SmG) - V3AB | GT | 0.09 | 0.02 | 864 | 4.69 | 0.00 |
| (AnG-SmG) - (V5/MT) | GT | 0.06 | 0.02 | 864 | 3.21 | 0.05 |
| FEFs - (IFS-PCSi) | GT | -0.02 | 0.02 | 864 | -0.91 | 1.00 |
| FEFs - LO | GT | 0.24 | 0.02 | 864 | 12.15 | 0.00 |
| FEFs - OcP | GT | 0.21 | 0.02 | 864 | 10.63 | 0.00 |
| FEFs - SMA | GT | 0.00 | 0.02 | 864 | 0.06 | 1.00 |
| FEFs - (IPS-SPL) | GT | 0.08 | 0.02 | 864 | 3.97 | 0.00 |
| FEFs - V3AB | GT | 0.12 | 0.02 | 864 | 6.15 | 0.00 |
| FEFs - (V5/MT) | GT | 0.09 | 0.02 | 864 | 4.67 | 0.00 |
| (IFS-PCSi) - LO | GT | 0.26 | 0.02 | 864 | 13.05 | 0.00 |
| (IFS-PCSi) - OcP | GT | 0.23 | 0.02 | 864 | 11.54 | 0.00 |

| | | | | | | |
|---|---|---|---|---|---|---|
| (IFS-PCSi) - SMA | GT | 0.02 | 0.02 | 864 | 0.97 | 1.00 |
| (IFS-PCSi) - (IPS-SPL) | GT | 0.10 | 0.02 | 864 | 4.87 | 0.00 |
| (IFS-PCSi) - V3AB | GT | 0.14 | 0.02 | 864 | 7.06 | 0.00 |
| (IFS-PCSi) - (V5/MT) | GT | 0.11 | 0.02 | 864 | 5.57 | 0.00 |
| LO - OcP | GT | -0.03 | 0.02 | 864 | -1.51 | 1.00 |
| LO - SMA | GT | -0.24 | 0.02 | 864 | -12.08 | 0.00 |
| LO - (IPS-SPL) | GT | -0.16 | 0.02 | 864 | -8.18 | 0.00 |
| LO - V3AB | GT | -0.12 | 0.02 | 864 | -6.00 | 0.00 |
| LO - (V5/MT) | GT | -0.15 | 0.02 | 864 | -7.48 | 0.00 |
| OcP - SMA | GT | -0.21 | 0.02 | 864 | -10.57 | 0.00 |
| OcP - (IPS-SPL) | GT | -0.13 | 0.02 | 864 | -6.67 | 0.00 |
| OcP - V3AB | GT | -0.09 | 0.02 | 864 | -4.48 | 0.00 |
| OcP - (V5/MT) | GT | -0.12 | 0.02 | 864 | -5.97 | 0.00 |
| SMA - (IPS-SPL) | GT | 0.08 | 0.02 | 864 | 3.90 | 0.00 |
| SMA - V3AB | GT | 0.12 | 0.02 | 864 | 6.09 | 0.00 |
| SMA - (V5/MT) | GT | 0.09 | 0.02 | 864 | 4.60 | 0.00 |
| (IPS-SPL) - V3AB | GT | 0.04 | 0.02 | 864 | 2.18 | 1.00 |
| (IPS-SPL) - (V5/MT) | GT | 0.01 | 0.02 | 864 | 0.70 | 1.00 |
| V3AB - (V5/MT) | GT | -0.03 | 0.02 | 864 | -1.48 | 1.00 |

Supplementary table 6: CMT, CMTS, GS, and GT models comparison - ROIs contrasts for each model type (Bonferroni corrected)

| | ROI | estimate | SE | df | t.ratio | p.value |
|---|---|---|---|---|---|---|
| CMT - CMTS | AnG-SmG | 0.04 | 0.02 | 864 | 1.86 | 0.38 |
| CMT - GS | AnG-SmG | -0.01 | 0.02 | 864 | -0.70 | 1.00 |
| CMT - GT | AnG-SmG | -0.05 | 0.02 | 864 | -2.60 | 0.06 |
| CMTS - GS | AnG-SmG | -0.05 | 0.02 | 864 | -2.56 | 0.06 |
| CMTS - GT | AnG-SmG | -0.09 | 0.02 | 864 | -4.45 | 0.00 |
| GS - GT | AnG-SmG | -0.04 | 0.02 | 864 | -1.89 | 0.35 |
| CMT - CMTS | FEFs | 0.03 | 0.02 | 864 | 1.53 | 0.75 |
| CMT - GS | FEFs | -0.01 | 0.02 | 864 | -0.29 | 1.00 |
| CMT - GT | FEFs | -0.09 | 0.02 | 864 | -4.51 | 0.00 |
| CMTS - GS | FEFs | -0.04 | 0.02 | 864 | -1.82 | 0.41 |
| CMTS - GT | FEFs | -0.12 | 0.02 | 864 | -6.04 | 0.00 |
| GS - GT | FEFs | -0.08 | 0.02 | 864 | -4.22 | 0.00 |
| CMT - CMTS | IFS-PCSi | 0.02 | 0.02 | 864 | 0.93 | 1.00 |
| CMT - GS | IFS-PCSi | -0.02 | 0.02 | 864 | -0.89 | 1.00 |
| CMT - GT | IFS-PCSi | -0.12 | 0.02 | 864 | -6.12 | 0.00 |
| CMTS - GS | IFS-PCSi | -0.04 | 0.02 | 864 | -1.82 | 0.41 |
| CMTS - GT | IFS-PCSi | -0.14 | 0.02 | 864 | -7.05 | 0.00 |
| GS - GT | IFS-PCSi | -0.11 | 0.02 | 864 | -5.23 | 0.00 |
| CMT - CMTS | LO | -0.42 | 0.02 | 864 | -20.84 | 0.00 |
| CMT - GS | LO | -0.23 | 0.02 | 864 | -11.59 | 0.00 |
| CMT - GT | LO | 0.01 | 0.02 | 864 | 0.46 | 1.00 |
| CMTS - GS | LO | 0.19 | 0.02 | 864 | 9.26 | 0.00 |
| CMTS - GT | LO | 0.43 | 0.02 | 864 | 21.31 | 0.00 |
| GS - GT | LO | 0.24 | 0.02 | 864 | 12.05 | 0.00 |
| CMT - CMTS | OcP | -0.41 | 0.02 | 864 | -20.17 | 0.00 |
| CMT - GS | OcP | -0.22 | 0.02 | 864 | -10.74 | 0.00 |
| CMT - GT | OcP | -0.02 | 0.02 | 864 | -1.05 | 1.00 |
| CMTS - GS | OcP | 0.19 | 0.02 | 864 | 9.43 | 0.00 |
| CMTS - GT | OcP | 0.38 | 0.02 | 864 | 19.12 | 0.00 |
| GS - GT | OcP | 0.19 | 0.02 | 864 | 9.69 | 0.00 |
| CMT - CMTS | SMA | 0.10 | 0.02 | 864 | 5.02 | 0.00 |
| CMT - GS | SMA | 0.07 | 0.02 | 864 | 3.29 | 0.01 |
| CMT - GT | SMA | -0.04 | 0.02 | 864 | -2.07 | 0.23 |
| CMTS - GS | SMA | -0.03 | 0.02 | 864 | -1.73 | 0.51 |
| CMTS - GT | SMA | -0.14 | 0.02 | 864 | -7.09 | 0.00 |
| GS - GT | SMA | -0.11 | 0.02 | 864 | -5.36 | 0.00 |
| CMT - CMTS | IPS-SPL | -0.06 | 0.02 | 864 | -3.09 | 0.01 |
| CMT - GS | IPS-SPL | -0.07 | 0.02 | 864 | -3.48 | 0.00 |
| CMT - GT | IPS-SPL | -0.04 | 0.02 | 864 | -1.83 | 0.41 |
| CMTS - GS | IPS-SPL | -0.01 | 0.02 | 864 | -0.38 | 1.00 |
| CMTS - GT | IPS-SPL | 0.03 | 0.02 | 864 | 1.27 | 1.00 |

| | | | | | | |
|---|---|---|---|---|---|---|
| GS - GT | IPS-SPL | 0.03 | 0.02 | 864 | 1.65 | 0.59 |
| CMT - CMTS | V3AB | -0.17 | 0.02 | 864 | -8.35 | 0.00 |
| CMT - GS | V3AB | -0.12 | 0.02 | 864 | -5.84 | 0.00 |
| CMT - GT | V3AB | -0.03 | 0.02 | 864 | -1.45 | 0.88 |
| CMTS - GS | V3AB | 0.05 | 0.02 | 864 | 2.52 | 0.07 |
| CMTS - GT | V3AB | 0.14 | 0.02 | 864 | 6.90 | 0.00 |
| GS - GT | V3AB | 0.09 | 0.02 | 864 | 4.38 | 0.00 |
| CMT - CMTS | V5/MT | -0.08 | 0.02 | 864 | -3.94 | 0.00 |
| CMT - GS | V5/MT | -0.06 | 0.02 | 864 | -3.01 | 0.02 |
| CMT - GT | V5/MT | -0.02 | 0.02 | 864 | -1.02 | 1.00 |
| CMTS - GS | V5/MT | 0.02 | 0.02 | 864 | 0.93 | 1.00 |
| CMTS - GT | V5/MT | 0.06 | 0.02 | 864 | 2.92 | 0.02 |
| GS - GT | V5/MT | 0.04 | 0.02 | 864 | 1.99 | 0.28 |

Supplementary table 7: CMT, CMTS, GS, and GT models comparison - model types contrasts for each ROI (Bonferroni corrected)

| | Sum.Sq | Mean.Sq | NumDF | DenDF | F.value | Pr(>F) |
|---|---|---|---|---|---|---|
| ROI | 8669.84 | 1238.55 | 7 | 180.00 | 7.41 | 0.00 |
| Hemi | 906.03 | 906.03 | 1 | 180.00 | 5.42 | 0.02 |
| ROI:Hemi | 1356.05 | 193.72 | 7 | 180.00 | 1.16 | 0.33 |

Supplementary table 8: pRF duration preference ($\mu_d$) - type III ANOVA on LME model estimates with Satterthwaite's method for degrees of freedom

| | estimate | SE | df | t.ratio | p.value |
|---|---|---|---|---|---|
| lh - rh | -4.17 | 1.79 | 180 | -2.33 | 0.02 |

Supplementary table 9: pRF duration preference ($\mu_d$) - hemispheres contrast (Bonferroni corrected) with Kenward-Roger's method for degrees of freedom

| | estimate | SE | df | t.ratio | p.value |
|---|---|---|---|---|---|
| FEFs - IPS0 | -10.30 | 3.59 | 180.00 | -2.87 | 0.13 |
| FEFs - IPS1 | -3.02 | 3.59 | 180.00 | -0.84 | 1.00 |
| FEFs - IPS2 | -3.42 | 3.59 | 180.00 | -0.95 | 1.00 |
| FEFs - IPS3 | -6.18 | 3.59 | 180.00 | -1.72 | 1.00 |
| FEFs - MT | -11.76 | 3.59 | 180.00 | -3.28 | 0.03 |
| FEFs - (V1d-V2d) | -21.23 | 3.59 | 180.00 | -5.92 | 0.00 |
| FEFs - V3AB | -13.43 | 3.59 | 180.00 | -3.74 | 0.01 |
| IPS0 - IPS1 | 7.28 | 3.59 | 180.00 | 2.03 | 1.00 |
| IPS0 - IPS2 | 6.88 | 3.59 | 180.00 | 1.92 | 1.00 |
| IPS0 - IPS3 | 4.12 | 3.59 | 180.00 | 1.15 | 1.00 |
| IPS0 - MT | -1.46 | 3.59 | 180.00 | -0.41 | 1.00 |
| IPS0 - (V1d-V2d) | -10.93 | 3.59 | 180.00 | -3.05 | 0.07 |
| IPS0 - V3AB | -3.13 | 3.59 | 180.00 | -0.87 | 1.00 |
| IPS1 - IPS2 | -0.40 | 3.59 | 180.00 | -0.11 | 1.00 |
| IPS1 - IPS3 | -3.16 | 3.59 | 180.00 | -0.88 | 1.00 |
| IPS1 - MT | -8.74 | 3.59 | 180.00 | -2.44 | 0.44 |
| IPS1 - (V1d-V2d) | -18.21 | 3.59 | 180.00 | -5.08 | 0.00 |
| IPS1 - V3AB | -10.41 | 3.59 | 180.00 | -2.90 | 0.12 |
| IPS2 - IPS3 | -2.76 | 3.59 | 180.00 | -0.77 | 1.00 |
| IPS2 - MT | -8.34 | 3.59 | 180.00 | -2.33 | 0.59 |
| IPS2 - (V1d-V2d) | -17.81 | 3.59 | 180.00 | -4.97 | 0.00 |
| IPS2 - V3AB | -10.01 | 3.59 | 180.00 | -2.79 | 0.16 |
| IPS3 - MT | -5.58 | 3.59 | 180.00 | -1.56 | 1.00 |
| IPS3 - (V1d-V2d) | -15.05 | 3.59 | 180.00 | -4.20 | 0.00 |
| IPS3 - V3AB | -7.25 | 3.59 | 180.00 | -2.02 | 1.00 |
| MT - (V1d-V2d) | -9.47 | 3.59 | 180.00 | -2.64 | 0.25 |
| MT - V3AB | -1.67 | 3.59 | 180.00 | -0.47 | 1.00 |
| (V1d-V2d) - V3AB | 7.80 | 3.59 | 180.00 | 2.18 | 0.87 |

Supplementary table 10: pRF duration preference ($\mu_d$) - ROIs contrasts (Bonferroni corrected) with Kenward-Roger's method for degrees of freedom

| | Sum.Sq | Mean.Sq | NumDF | DenDF | F.value | Pr(>F) |
|---|---|---|---|---|---|---|
| ROI | 18.33 | 2.62 | 7 | 180.00 | 29.27 | 0.00 |
| Hemi | 0.08 | 0.08 | 1 | 180.00 | 0.91 | 0.34 |
| ROI:Hemi | 0.96 | 0.14 | 7 | 180.00 | 1.54 | 0.16 |

Supplementary table 11: pRF $\mu_s$-$\mu_r$ correlation - type III ANOVA on LME model estimates with Satterthwaite's method for degrees of freedom

| | estimate | SE | df | t.ratio | p.value |
|---|---|---|---|---|---|
| (V1d-V2d) - FEFs | 1.00 | 0.08 | 180.00 | 12.10 | 0.00 |
| (V1d-V2d) - IPS0 | 0.70 | 0.08 | 180.00 | 8.50 | 0.00 |
| (V1d-V2d) - IPS1 | 0.79 | 0.08 | 180.00 | 9.53 | 0.00 |
| (V1d-V2d) - IPS2 | 0.76 | 0.08 | 180.00 | 9.21 | 0.00 |
| (V1d-V2d) - IPS3 | 0.81 | 0.08 | 180.00 | 9.78 | 0.00 |
| (V1d-V2d) - MT | 0.46 | 0.08 | 180.00 | 5.60 | 0.00 |
| (V1d-V2d) - V3AB | 0.38 | 0.08 | 180.00 | 4.63 | 0.00 |
| FEFs - IPS0 | -0.30 | 0.08 | 180.00 | -3.60 | 0.01 |
| FEFs - IPS1 | -0.21 | 0.08 | 180.00 | -2.57 | 0.31 |
| FEFs - IPS2 | -0.24 | 0.08 | 180.00 | -2.89 | 0.12 |
| FEFs - IPS3 | -0.19 | 0.08 | 180.00 | -2.32 | 0.60 |
| FEFs - MT | -0.54 | 0.08 | 180.00 | -6.50 | 0.00 |
| FEFs - V3AB | -0.62 | 0.08 | 180.00 | -7.47 | 0.00 |
| IPS0 - IPS1 | 0.09 | 0.08 | 180.00 | 1.04 | 1.00 |
| IPS0 - IPS2 | 0.06 | 0.08 | 180.00 | 0.72 | 1.00 |
| IPS0 - IPS3 | 0.11 | 0.08 | 180.00 | 1.28 | 1.00 |
| IPS0 - MT | -0.24 | 0.08 | 180.00 | -2.90 | 0.12 |
| IPS0 - V3AB | -0.32 | 0.08 | 180.00 | -3.86 | 0.00 |
| IPS1 - IPS2 | -0.03 | 0.08 | 180.00 | -0.32 | 1.00 |
| IPS1 - IPS3 | 0.02 | 0.08 | 180.00 | 0.25 | 1.00 |
| IPS1 - MT | -0.33 | 0.08 | 180.00 | -3.93 | 0.00 |
| IPS1 - V3AB | -0.41 | 0.08 | 180.00 | -4.90 | 0.00 |
| IPS2 - IPS3 | 0.05 | 0.08 | 180.00 | 0.56 | 1.00 |
| IPS2 - MT | -0.30 | 0.08 | 180.00 | -3.61 | 0.01 |
| IPS2 - V3AB | -0.38 | 0.08 | 180.00 | -4.58 | 0.00 |
| IPS3 - MT | -0.35 | 0.08 | 180.00 | -4.18 | 0.00 |
| IPS3 - V3AB | -0.43 | 0.08 | 180.00 | -5.15 | 0.00 |
| MT - V3AB | -0.08 | 0.08 | 180.00 | -0.97 | 1.00 |

Supplementary table 12: pRF $\mu_s$-$\mu_r$ correlation - ROIs contrasts (Bonferroni corrected) with Kenward-Roger's method for degrees of freedom

| | Pg($\chi^2$) | df | p($\chi^2$) | p(perm) |
|---|---|---|---|---|
| median $\theta$ | 30.77 | 7 | 0.00 | 0.00 |

Supplementary table 13: pRF orientation ($\theta$) - Fisher's non-parametric test

| | med.diff | Pg($\chi^2$) | p($\chi^2$) | bonf_p($\chi^2$) | p(perm) | bonf_p(perm) |
|---|---|---|---|---|---|---|
| FEFs-IPS0 | 0.01 | 0.15 | 0.69 | 1.00 | 1.00 | 1.00 |
| FEFs-IPS1 | -0.22 | 3.85 | 0.05 | 1.00 | 0.11 | 1.00 |
| FEFs-IPS2 | -0.03 | 0.15 | 0.69 | 1.00 | 1.00 | 1.00 |
| FEFs-IPS3 | -0.12 | 1.38 | 0.24 | 1.00 | 0.44 | 1.00 |
| FEFs-MT | 0.13 | 3.85 | 0.05 | 1.00 | 0.11 | 1.00 |
| FEFs-V1d-V2d | 0.30 | 12.46 | 0.00 | 0.01 | 0.00 | 0.05 |
| FEFs-V3AB | 0.28 | 7.54 | 0.01 | 0.17 | 0.01 | 0.40 |
| IPS0-IPS1 | -0.24 | 3.85 | 0.05 | 1.00 | 0.12 | 1.00 |
| IPS0-IPS2 | -0.05 | 0.15 | 0.69 | 1.00 | 1.00 | 1.00 |
| IPS0-IPS3 | -0.13 | 0.15 | 0.69 | 1.00 | 1.00 | 1.00 |
| IPS0-MT | 0.11 | 1.38 | 0.24 | 1.00 | 0.43 | 1.00 |
| IPS0-V1d-V2d | 0.29 | 18.62 | 0.00 | 0.00 | 0.00 | 0.00 |
| IPS0-V3AB | 0.26 | 3.85 | 0.05 | 1.00 | 0.12 | 1.00 |
| IPS1-IPS2 | 0.19 | 1.38 | 0.24 | 1.00 | 0.43 | 1.00 |
| IPS1-IPS3 | 0.10 | 1.38 | 0.24 | 1.00 | 0.43 | 1.00 |
| IPS1-MT | 0.35 | 3.85 | 0.05 | 1.00 | 0.11 | 1.00 |
| IPS1-V1d-V2d | 0.52 | 18.62 | 0.00 | 0.00 | 0.00 | 0.00 |
| IPS1-V3AB | 0.50 | 18.62 | 0.00 | 0.00 | 0.00 | 0.00 |
| IPS2-IPS3 | -0.09 | 0.15 | 0.69 | 1.00 | 1.00 | 1.00 |
| IPS2-MT | 0.16 | 1.38 | 0.24 | 1.00 | 0.44 | 1.00 |
| IPS2-V1d-V2d | 0.34 | 12.46 | 0.00 | 0.01 | 0.00 | 0.03 |
| IPS2-V3AB | 0.31 | 1.38 | 0.24 | 1.00 | 0.44 | 1.00 |
| IPS3-MT | 0.25 | 3.85 | 0.05 | 1.00 | 0.12 | 1.00 |
| IPS3-V1d-V2d | 0.42 | 18.62 | 0.00 | 0.00 | 0.00 | 0.00 |
| IPS3-V3AB | 0.39 | 7.54 | 0.01 | 0.17 | 0.02 | 0.50 |
| MT-V1d-V2d | 0.17 | 18.62 | 0.00 | 0.00 | 0.00 | 0.00 |
| MT-V3AB | 0.15 | 1.38 | 0.24 | 1.00 | 0.44 | 1.00 |
| V1d-V2d-V3AB | -0.03 | 7.54 | 0.01 | 0.17 | 0.02 | 0.50 |

Supplementary table 14: pRF orientation ($\theta$) - ROIs contrasts (Bonferroni corrected)

| | Sum.Sq | Mean.Sq | NumDF | DenDF | F.value | Pr(>F) |
|---|---|---|---|---|---|---|
| ROI | 0.46 | 0.07 | 7 | 180.00 | 2.80 | 0.01 |
| Hemi | 0.11 | 0.11 | 1 | 180.00 | 4.88 | 0.03 |
| ROI:Hemi | 0.11 | 0.02 | 7 | 180.00 | 0.68 | 0.69 |

Supplementary table 15: pRF aspect ratio - type III ANOVA on LME model estimates with Satterthwaite's method for degrees of freedom

| | estimate | SE | df | t.ratio | p.value |
|---|---|---|---|---|---|
| lh - rh | 0.05 | 0.02 | 180 | 2.21 | 0.03 |

Supplementary table 16: pRF aspect ratio - hemispheres contrast (Bonferroni corrected) with Kenward-Roger's method for degrees of freedom. Contrast estimate is expressed as difference of log scaled values

| | estimate | SE | df | t.ratio | p.value |
|---|---|---|---|---|---|
| FEFs - IPS0 | 0.12 | 0.04 | 180.00 | 2.81 | 0.15 |
| FEFs - IPS1 | 0.09 | 0.04 | 180.00 | 2.20 | 0.81 |
| FEFs - IPS2 | 0.03 | 0.04 | 180.00 | 0.59 | 1.00 |
| FEFs - IPS3 | 0.06 | 0.04 | 180.00 | 1.46 | 1.00 |
| FEFs - MT | 0.09 | 0.04 | 180.00 | 2.03 | 1.00 |
| FEFs - (V1d-V2d) | -0.03 | 0.04 | 180.00 | -0.70 | 1.00 |
| FEFs - V3AB | 0.05 | 0.04 | 180.00 | 1.10 | 1.00 |
| IPS0 - IPS1 | -0.03 | 0.04 | 180.00 | -0.61 | 1.00 |
| IPS0 - IPS2 | -0.09 | 0.04 | 180.00 | -2.22 | 0.77 |
| IPS0 - IPS3 | -0.06 | 0.04 | 180.00 | -1.35 | 1.00 |
| IPS0 - MT | -0.03 | 0.04 | 180.00 | -0.78 | 1.00 |
| IPS0 - (V1d-V2d) | -0.15 | 0.04 | 180.00 | -3.51 | 0.02 |
| IPS0 - V3AB | -0.07 | 0.04 | 180.00 | -1.71 | 1.00 |
| IPS1 - IPS2 | -0.07 | 0.04 | 180.00 | -1.61 | 1.00 |
| IPS1 - IPS3 | -0.03 | 0.04 | 180.00 | -0.74 | 1.00 |
| IPS1 - MT | -0.01 | 0.04 | 180.00 | -0.17 | 1.00 |
| IPS1 - (V1d-V2d) | -0.12 | 0.04 | 180.00 | -2.90 | 0.12 |
| IPS1 - V3AB | -0.05 | 0.04 | 180.00 | -1.10 | 1.00 |
| IPS2 - IPS3 | 0.04 | 0.04 | 180.00 | 0.87 | 1.00 |
| IPS2 - MT | 0.06 | 0.04 | 180.00 | 1.44 | 1.00 |
| IPS2 - (V1d-V2d) | -0.05 | 0.04 | 180.00 | -1.29 | 1.00 |
| IPS2 - V3AB | 0.02 | 0.04 | 180.00 | 0.51 | 1.00 |
| IPS3 - MT | 0.02 | 0.04 | 180.00 | 0.57 | 1.00 |
| IPS3 - (V1d-V2d) | -0.09 | 0.04 | 180.00 | -2.16 | 0.90 |
| IPS3 - V3AB | -0.02 | 0.04 | 180.00 | -0.36 | 1.00 |
| MT - (V1d-V2d) | -0.12 | 0.04 | 180.00 | -2.73 | 0.19 |
| MT - V3AB | -0.04 | 0.04 | 180.00 | -0.93 | 1.00 |
| (V1d-V2d) - V3AB | 0.08 | 0.04 | 180.00 | 1.80 | 1.00 |

Supplementary table 17: pRF aspect ratio - ROIs contrasts (Bonferroni corrected) with Kenward-Roger's method for degrees of freedom. Contrast estimates are expressed as difference of log scaled values

| | DFn | DFd | F | p | p<.05 | ges |
|---|---|---|---|---|---|---|
| ROI | 7 | 84 | 0.00 | 1.00 | | 0.00 |
| Hemi | 1 | 12 | -0.00 | 1.00 | | 0.00 |
| StimDim | 2 | 24 | 72.16 | 0.00 | * | 0.65 |
| ROI:Hemi | 7 | 84 | -0.00 | 1.00 | | 0.00 |
| ROI:StimDim | 14 | 168 | 17.43 | 0.00 | * | 0.34 |
| Hemi:StimDim | 2 | 24 | 0.45 | 0.64 | | 0.00 |
| ROI:Hemi:StimDim | 14 | 168 | 0.85 | 0.61 | | 0.02 |

Supplementary table 18: pRF selectivity - three-way repeated measure ANOVA

| | W | p | p<.05 |
|---|---|---|---|
| StimDim | 0.20 | 0.00 | * |
| Hemi:StimDim | 0.96 | 0.81 | |

Supplementary table 19: pRF selectivity - Mauchly's sphericity test

| | GGe | DF[GG] | p[GG] | p[GG]<.05 | HFe | DF[HF] | p[HF] | p[HF]<.05 |
|---|---|---|---|---|---|---|---|---|
| StimDim | 0.56 | 1.11, 13.34 | 0.00 | * | 0.57 | 1.14, 13.72 | 0.00 | * |
| Hemi:StimDim | 0.96 | 1.93, 23.12 | 0.64 | | 1.14 | 2.29, 27.45 | 0.64 | |

Supplementary table 20: pRF selectivity - deviation from sphericity corrections

| | StimDim | estimate | SE | df | t.ratio | p.value |
|---|---|---|---|---|---|---|
| FEFs - IPS0 | Both | 0.11 | 0.04 | 576 | 2.41 | 0.45 |
| FEFs - IPS1 | Both | -0.02 | 0.04 | 576 | -0.43 | 1.00 |
| FEFs - IPS2 | Both | 0.06 | 0.04 | 576 | 1.34 | 1.00 |
| FEFs - IPS3 | Both | 0.03 | 0.04 | 576 | 0.62 | 1.00 |
| FEFs - MT | Both | 0.15 | 0.04 | 576 | 3.34 | 0.03 |
| FEFs - (V1d-V2d) | Both | 0.32 | 0.04 | 576 | 7.26 | 0.00 |
| FEFs - V3AB | Both | 0.17 | 0.04 | 576 | 3.81 | 0.00 |
| IPS0 - IPS1 | Both | -0.13 | 0.04 | 576 | -2.84 | 0.13 |
| IPS0 - IPS2 | Both | -0.05 | 0.04 | 576 | -1.07 | 1.00 |
| IPS0 - IPS3 | Both | -0.08 | 0.04 | 576 | -1.79 | 1.00 |
| IPS0 - MT | Both | 0.04 | 0.04 | 576 | 0.92 | 1.00 |
| IPS0 - (V1d-V2d) | Both | 0.21 | 0.04 | 576 | 4.85 | 0.00 |
| IPS0 - V3AB | Both | 0.06 | 0.04 | 576 | 1.40 | 1.00 |
| IPS1 - IPS2 | Both | 0.08 | 0.04 | 576 | 1.77 | 1.00 |
| IPS1 - IPS3 | Both | 0.05 | 0.04 | 576 | 1.05 | 1.00 |
| IPS1 - MT | Both | 0.17 | 0.04 | 576 | 3.76 | 0.01 |
| IPS1 - (V1d-V2d) | Both | 0.34 | 0.04 | 576 | 7.69 | 0.00 |
| IPS1 - V3AB | Both | 0.19 | 0.04 | 576 | 4.24 | 0.00 |
| IPS2 - IPS3 | Both | -0.03 | 0.04 | 576 | -0.73 | 1.00 |
| IPS2 - MT | Both | 0.09 | 0.04 | 576 | 1.99 | 1.00 |
| IPS2 - (V1d-V2d) | Both | 0.26 | 0.04 | 576 | 5.92 | 0.00 |
| IPS2 - V3AB | Both | 0.11 | 0.04 | 576 | 2.47 | 0.39 |
| IPS3 - MT | Both | 0.12 | 0.04 | 576 | 2.72 | 0.19 |
| IPS3 - (V1d-V2d) | Both | 0.29 | 0.04 | 576 | 6.64 | 0.00 |
| IPS3 - V3AB | Both | 0.14 | 0.04 | 576 | 3.19 | 0.04 |
| MT - (V1d-V2d) | Both | 0.17 | 0.04 | 576 | 3.93 | 0.00 |
| MT - V3AB | Both | 0.02 | 0.04 | 576 | 0.48 | 1.00 |
| (V1d-V2d) - V3AB | Both | -0.15 | 0.04 | 576 | -3.45 | 0.02 |
| FEFs - IPS0 | Space | -0.19 | 0.04 | 576 | -4.39 | 0.00 |
| FEFs - IPS1 | Space | -0.04 | 0.04 | 576 | -0.91 | 1.00 |
| FEFs - IPS2 | Space | -0.09 | 0.04 | 576 | -2.14 | 0.91 |
| FEFs - IPS3 | Space | -0.06 | 0.04 | 576 | -1.41 | 1.00 |
| FEFs - MT | Space | -0.26 | 0.04 | 576 | -5.87 | 0.00 |
| FEFs - (V1d-V2d) | Space | -0.49 | 0.04 | 576 | -11.04 | 0.00 |
| FEFs - V3AB | Space | -0.29 | 0.04 | 576 | -6.62 | 0.00 |
| IPS0 - IPS1 | Space | 0.15 | 0.04 | 576 | 3.47 | 0.02 |
| IPS0 - IPS2 | Space | 0.10 | 0.04 | 576 | 2.24 | 0.71 |
| IPS0 - IPS3 | Space | 0.13 | 0.04 | 576 | 2.97 | 0.09 |
| IPS0 - MT | Space | -0.07 | 0.04 | 576 | -1.49 | 1.00 |
| IPS0 - (V1d-V2d) | Space | -0.29 | 0.04 | 576 | -6.65 | 0.00 |
| IPS0 - V3AB | Space | -0.10 | 0.04 | 576 | -2.23 | 0.73 |

| | | | | | | |
|---|---|---|---|---|---|---|
| IPS1 - IPS2 | Space | -0.05 | 0.04 | 576 | -1.23 | 1.00 |
| IPS1 - IPS3 | Space | -0.02 | 0.04 | 576 | -0.50 | 1.00 |
| IPS1 - MT | Space | -0.22 | 0.04 | 576 | -4.96 | 0.00 |
| IPS1 - (V1d-V2d) | Space | -0.45 | 0.04 | 576 | -10.13 | 0.00 |
| IPS1 - V3AB | Space | -0.25 | 0.04 | 576 | -5.70 | 0.00 |
| IPS2 - IPS3 | Space | 0.03 | 0.04 | 576 | 0.73 | 1.00 |
| IPS2 - MT | Space | -0.16 | 0.04 | 576 | -3.73 | 0.01 |
| IPS2 - (V1d-V2d) | Space | -0.39 | 0.04 | 576 | -8.90 | 0.00 |
| IPS2 - V3AB | Space | -0.20 | 0.04 | 576 | -4.47 | 0.00 |
| IPS3 - MT | Space | -0.20 | 0.04 | 576 | -4.46 | 0.00 |
| IPS3 - (V1d-V2d) | Space | -0.42 | 0.04 | 576 | -9.63 | 0.00 |
| IPS3 - V3AB | Space | -0.23 | 0.04 | 576 | -5.20 | 0.00 |
| MT - (V1d-V2d) | Space | -0.23 | 0.04 | 576 | -5.17 | 0.00 |
| MT - V3AB | Space | -0.03 | 0.04 | 576 | -0.74 | 1.00 |
| (V1d-V2d) - V3AB | Space | 0.20 | 0.04 | 576 | 4.43 | 0.00 |
| FEFs - IPS0 | Time | 0.09 | 0.04 | 576 | 1.97 | 1.00 |
| FEFs - IPS1 | Time | 0.06 | 0.04 | 576 | 1.34 | 1.00 |
| FEFs - IPS2 | Time | 0.04 | 0.04 | 576 | 0.80 | 1.00 |
| FEFs - IPS3 | Time | 0.04 | 0.04 | 576 | 0.79 | 1.00 |
| FEFs - MT | Time | 0.11 | 0.04 | 576 | 2.54 | 0.32 |
| FEFs - (V1d-V2d) | Time | 0.17 | 0.04 | 576 | 3.78 | 0.00 |
| FEFs - V3AB | Time | 0.12 | 0.04 | 576 | 2.80 | 0.15 |
| IPS0 - IPS1 | Time | -0.03 | 0.04 | 576 | -0.63 | 1.00 |
| IPS0 - IPS2 | Time | -0.05 | 0.04 | 576 | -1.17 | 1.00 |
| IPS0 - IPS3 | Time | -0.05 | 0.04 | 576 | -1.18 | 1.00 |
| IPS0 - MT | Time | 0.02 | 0.04 | 576 | 0.56 | 1.00 |
| IPS0 - (V1d-V2d) | Time | 0.08 | 0.04 | 576 | 1.80 | 1.00 |
| IPS0 - V3AB | Time | 0.04 | 0.04 | 576 | 0.83 | 1.00 |
| IPS1 - IPS2 | Time | -0.02 | 0.04 | 576 | -0.54 | 1.00 |
| IPS1 - IPS3 | Time | -0.02 | 0.04 | 576 | -0.55 | 1.00 |
| IPS1 - MT | Time | 0.05 | 0.04 | 576 | 1.19 | 1.00 |
| IPS1 - (V1d-V2d) | Time | 0.11 | 0.04 | 576 | 2.44 | 0.42 |
| IPS1 - V3AB | Time | 0.06 | 0.04 | 576 | 1.46 | 1.00 |
| IPS2 - IPS3 | Time | -0.00 | 0.04 | 576 | -0.00 | 1.00 |
| IPS2 - MT | Time | 0.08 | 0.04 | 576 | 1.74 | 1.00 |
| IPS2 - (V1d-V2d) | Time | 0.13 | 0.04 | 576 | 2.98 | 0.08 |
| IPS2 - V3AB | Time | 0.09 | 0.04 | 576 | 2.00 | 1.00 |
| IPS3 - MT | Time | 0.08 | 0.04 | 576 | 1.74 | 1.00 |
| IPS3 - (V1d-V2d) | Time | 0.13 | 0.04 | 576 | 2.98 | 0.08 |
| IPS3 - V3AB | Time | 0.09 | 0.04 | 576 | 2.01 | 1.00 |
| MT - (V1d-V2d) | Time | 0.05 | 0.04 | 576 | 1.24 | 1.00 |
| MT - V3AB | Time | 0.01 | 0.04 | 576 | 0.27 | 1.00 |

| | | | | | | |
|---|---|---|---|---|---|---|
| (V1d-V2d) - V3AB | Time | -0.04 | 0.04 | 576 | -0.98 | 1.00 |

Supplementary table 21: pRF selectivity - ROIs contrasts for each stimulus dimension (Bonferroni corrected)

| | ROI | estimate | SE | df | t.ratio | p.value |
|---|---|---|---|---|---|---|
| Both - Space | FEFs | -0.04 | 0.04 | 576 | -0.97 | 1.00 |
| Both - Time | FEFs | 0.21 | 0.04 | 576 | 4.67 | 0.00 |
| Space - Time | FEFs | 0.25 | 0.04 | 576 | 5.64 | 0.00 |
| Both - Space | IPS0 | -0.34 | 0.04 | 576 | -7.77 | 0.00 |
| Both - Time | IPS0 | 0.19 | 0.04 | 576 | 4.23 | 0.00 |
| Space - Time | IPS0 | 0.53 | 0.04 | 576 | 12.00 | 0.00 |
| Both - Space | IPS1 | -0.06 | 0.04 | 576 | -1.46 | 0.44 |
| Both - Time | IPS1 | 0.28 | 0.04 | 576 | 6.44 | 0.00 |
| Space - Time | IPS1 | 0.35 | 0.04 | 576 | 7.90 | 0.00 |
| Both - Space | IPS2 | -0.20 | 0.04 | 576 | -4.46 | 0.00 |
| Both - Time | IPS2 | 0.18 | 0.04 | 576 | 4.13 | 0.00 |
| Space - Time | IPS2 | 0.38 | 0.04 | 576 | 8.59 | 0.00 |
| Both - Space | IPS3 | -0.13 | 0.04 | 576 | -3.00 | 0.01 |
| Both - Time | IPS3 | 0.21 | 0.04 | 576 | 4.85 | 0.00 |
| Space - Time | IPS3 | 0.35 | 0.04 | 576 | 7.85 | 0.00 |
| Both - Space | MT | -0.45 | 0.04 | 576 | -10.18 | 0.00 |
| Both - Time | MT | 0.17 | 0.04 | 576 | 3.87 | 0.00 |
| Space - Time | MT | 0.62 | 0.04 | 576 | 14.05 | 0.00 |
| Both - Space | V1d-V2d | -0.85 | 0.04 | 576 | -19.27 | 0.00 |
| Both - Time | V1d-V2d | 0.05 | 0.04 | 576 | 1.19 | 0.70 |
| Space - Time | V1d-V2d | 0.90 | 0.04 | 576 | 20.46 | 0.00 |
| Both - Space | V3AB | -0.50 | 0.04 | 576 | -11.40 | 0.00 |
| Both - Time | V3AB | 0.16 | 0.04 | 576 | 3.66 | 0.00 |
| Space - Time | V3AB | 0.66 | 0.04 | 576 | 15.06 | 0.00 |

Supplementary table 22: pRF selectivity - stimulus dimensions contrasts for each ROI (Bonferroni corrected)

| | Pg($\chi^2$) | df | p($\chi^2$) | p(perm) |
|---|---|---|---|---|
| median $\alpha_g$ | 27.08 | 7 | 0.00 | 0.00 |

Supplementary table 23: $\alpha_g$ - Fisher's non-parametric test

| | med.diff | Pg($\chi^2$) | p($\chi^2$) | bonf_p($\chi^2$) | p(perm) | bonf_p(perm) |
|---|---|---|---|---|---|---|
| FEFs-IPS0 | -0.22 | 1.38 | 0.24 | 1.00 | 0.45 | 1.00 |
| FEFs-IPS1 | -0.12 | 0.15 | 0.69 | 1.00 | 1.00 | 1.00 |
| FEFs-IPS2 | -0.82 | 7.54 | 0.01 | 0.17 | 0.02 | 0.43 |
| FEFs-IPS3 | -0.11 | 1.38 | 0.24 | 1.00 | 0.43 | 1.00 |
| FEFs-MT | -0.31 | 3.85 | 0.05 | 1.00 | 0.12 | 1.00 |
| FEFs-V1d-V2d | -1.17 | 18.62 | 0.00 | 0.00 | 0.00 | 0.00 |
| FEFs-V3AB | -0.53 | 3.85 | 0.05 | 1.00 | 0.12 | 1.00 |
| IPS0-IPS1 | 0.10 | 0.15 | 0.69 | 1.00 | 1.00 | 1.00 |
| IPS0-IPS2 | -0.61 | 7.54 | 0.01 | 0.17 | 0.02 | 0.48 |
| IPS0-IPS3 | 0.11 | 0.15 | 0.69 | 1.00 | 1.00 | 1.00 |
| IPS0-MT | -0.09 | 1.38 | 0.24 | 1.00 | 0.44 | 1.00 |
| IPS0-V1d-V2d | -0.95 | 12.46 | 0.00 | 0.01 | 0.00 | 0.05 |
| IPS0-V3AB | -0.32 | 3.85 | 0.05 | 1.00 | 0.11 | 1.00 |
| IPS1-IPS2 | -0.71 | 3.85 | 0.05 | 1.00 | 0.11 | 1.00 |
| IPS1-IPS3 | 0.01 | 0.15 | 0.69 | 1.00 | 1.00 | 1.00 |
| IPS1-MT | -0.19 | 0.15 | 0.69 | 1.00 | 1.00 | 1.00 |
| IPS1-V1d-V2d | -1.05 | 7.54 | 0.01 | 0.17 | 0.02 | 0.49 |
| IPS1-V3AB | -0.42 | 1.38 | 0.24 | 1.00 | 0.43 | 1.00 |
| IPS2-IPS3 | 0.71 | 7.54 | 0.01 | 0.17 | 0.02 | 0.50 |
| IPS2-MT | 0.52 | 3.85 | 0.05 | 1.00 | 0.11 | 1.00 |
| IPS2-V1d-V2d | -0.35 | 3.85 | 0.05 | 1.00 | 0.11 | 1.00 |
| IPS2-V3AB | 0.29 | 1.38 | 0.24 | 1.00 | 0.43 | 1.00 |
| IPS3-MT | -0.20 | 1.38 | 0.24 | 1.00 | 0.44 | 1.00 |
| IPS3-V1d-V2d | -1.06 | 18.62 | 0.00 | 0.00 | 0.00 | 0.00 |
| IPS3-V3AB | -0.42 | 3.85 | 0.05 | 1.00 | 0.11 | 1.00 |
| MT-V1d-V2d | -0.86 | 12.46 | 0.00 | 0.01 | 0.00 | 0.05 |
| MT-V3AB | -0.23 | 1.38 | 0.24 | 1.00 | 0.45 | 1.00 |
| V1d-V2d-V3AB | 0.64 | 3.85 | 0.05 | 1.00 | 0.12 | 1.00 |

Supplementary table 24: $\alpha_g$ - ROIs contrasts (Bonferroni corrected)

| | Sum.Sq | Mean.Sq | NumDF | DenDF | F.value | Pr(>F) |
|---|---|---|---|---|---|---|
| ROI | 0.30 | 0.04 | 7 | 372.00 | 6.00 | 0.00 |
| MapType | 2.84 | 2.84 | 1 | 372.00 | 396.34 | 0.00 |
| Hemi | 0.01 | 0.01 | 1 | 372.00 | 0.79 | 0.37 |
| ROI:MapType | 0.42 | 0.06 | 7 | 372.00 | 8.32 | 0.00 |
| ROI:Hemi | 0.04 | 0.01 | 7 | 372.00 | 0.79 | 0.60 |
| MapType:Hemi | 0.00 | 0.00 | 1 | 372.00 | 0.34 | 0.56 |
| ROI:MapType:Hemi | 0.03 | 0.00 | 7 | 372.00 | 0.58 | 0.77 |

Supplementary table 25: Nuggets - type III ANOVA on LME model estimates with Satterthwaite's method for degrees of freedom

| | ROI | estimate | SE | df | t.ratio | p.value |
|---|---|---|---|---|---|---|
| Space - Time | FEFs | -0.13 | 0.02 | 372.00 | -5.71 | 0.00 |
| Space - Time | IPS0 | -0.14 | 0.02 | 372.00 | -6.06 | 0.00 |
| Space - Time | IPS1 | -0.09 | 0.02 | 372.00 | -3.89 | 0.00 |
| Space - Time | IPS2 | -0.10 | 0.02 | 372.00 | -4.22 | 0.00 |
| Space - Time | IPS3 | -0.14 | 0.02 | 372.00 | -6.10 | 0.00 |
| Space - Time | MT | -0.18 | 0.02 | 372.00 | -7.74 | 0.00 |
| Space - Time | V1d-V2d | -0.27 | 0.02 | 372.00 | -11.47 | 0.00 |
| Space - Time | V3AB | -0.26 | 0.02 | 372.00 | -11.12 | 0.00 |

Supplementary table 26: Nuggets - map types contrasts for each ROI (Bonferroni corrected) with Kenward-Roger's method for degrees of freedom

| | MapType | estimate | SE | df | t.ratio | p.value |
|---|---|---|---|---|---|---|
| FEFs - IPS0 | Space | 0.03 | 0.02 | 372.00 | 1.34 | 1.00 |
| FEFs - IPS1 | Space | -0.03 | 0.02 | 372.00 | -1.20 | 1.00 |
| FEFs - IPS2 | Space | -0.05 | 0.02 | 372.00 | -1.98 | 1.00 |
| FEFs - IPS3 | Space | -0.01 | 0.02 | 372.00 | -0.56 | 1.00 |
| FEFs - MT | Space | 0.06 | 0.02 | 372.00 | 2.36 | 0.52 |
| FEFs - (V1d-V2d) | Space | 0.12 | 0.02 | 372.00 | 5.16 | 0.00 |
| FEFs - V3AB | Space | 0.10 | 0.02 | 372.00 | 4.08 | 0.00 |
| IPS0 - IPS1 | Space | -0.06 | 0.02 | 372.00 | -2.53 | 0.33 |
| IPS0 - IPS2 | Space | -0.08 | 0.02 | 372.00 | -3.31 | 0.03 |
| IPS0 - IPS3 | Space | -0.04 | 0.02 | 372.00 | -1.89 | 1.00 |
| IPS0 - MT | Space | 0.02 | 0.02 | 372.00 | 1.03 | 1.00 |
| IPS0 - (V1d-V2d) | Space | 0.09 | 0.02 | 372.00 | 3.82 | 0.00 |
| IPS0 - V3AB | Space | 0.06 | 0.02 | 372.00 | 2.74 | 0.18 |
| IPS1 - IPS2 | Space | -0.02 | 0.02 | 372.00 | -0.78 | 1.00 |
| IPS1 - IPS3 | Space | 0.02 | 0.02 | 372.00 | 0.64 | 1.00 |
| IPS1 - MT | Space | 0.08 | 0.02 | 372.00 | 3.56 | 0.01 |
| IPS1 - (V1d-V2d) | Space | 0.15 | 0.02 | 372.00 | 6.35 | 0.00 |
| IPS1 - V3AB | Space | 0.12 | 0.02 | 372.00 | 5.27 | 0.00 |
| IPS2 - IPS3 | Space | 0.03 | 0.02 | 372.00 | 1.42 | 1.00 |
| IPS2 - MT | Space | 0.10 | 0.02 | 372.00 | 4.34 | 0.00 |
| IPS2 - (V1d-V2d) | Space | 0.17 | 0.02 | 372.00 | 7.13 | 0.00 |
| IPS2 - V3AB | Space | 0.14 | 0.02 | 372.00 | 6.05 | 0.00 |
| IPS3 - MT | Space | 0.07 | 0.02 | 372.00 | 2.92 | 0.10 |
| IPS3 - (V1d-V2d) | Space | 0.13 | 0.02 | 372.00 | 5.71 | 0.00 |
| IPS3 - V3AB | Space | 0.11 | 0.02 | 372.00 | 4.63 | 0.00 |
| MT - (V1d-V2d) | Space | 0.07 | 0.02 | 372.00 | 2.79 | 0.15 |
| MT - V3AB | Space | 0.04 | 0.02 | 372.00 | 1.71 | 1.00 |
| (V1d-V2d) - V3AB | Space | -0.03 | 0.02 | 372.00 | -1.08 | 1.00 |
| FEFs - IPS0 | Time | 0.02 | 0.02 | 372.00 | 0.98 | 1.00 |
| FEFs - IPS1 | Time | 0.01 | 0.02 | 372.00 | 0.62 | 1.00 |
| FEFs - IPS2 | Time | -0.01 | 0.02 | 372.00 | -0.49 | 1.00 |
| FEFs - IPS3 | Time | -0.02 | 0.02 | 372.00 | -0.95 | 1.00 |
| FEFs - MT | Time | 0.01 | 0.02 | 372.00 | 0.33 | 1.00 |
| FEFs - (V1d-V2d) | Time | -0.01 | 0.02 | 372.00 | -0.60 | 1.00 |
| FEFs - V3AB | Time | -0.03 | 0.02 | 372.00 | -1.33 | 1.00 |
| IPS0 - IPS1 | Time | -0.01 | 0.02 | 372.00 | -0.36 | 1.00 |
| IPS0 - IPS2 | Time | -0.03 | 0.02 | 372.00 | -1.47 | 1.00 |
| IPS0 - IPS3 | Time | -0.05 | 0.02 | 372.00 | -1.93 | 1.00 |
| IPS0 - MT | Time | -0.02 | 0.02 | 372.00 | -0.65 | 1.00 |
| IPS0 - (V1d-V2d) | Time | -0.04 | 0.02 | 372.00 | -1.58 | 1.00 |
| IPS0 - V3AB | Time | -0.05 | 0.02 | 372.00 | -2.31 | 0.60 |

| | | | | | | |
|---|---|---|---|---|---|---|
| IPS1 - IPS2 | Time | -0.03 | 0.02 | 372.00 | -1.11 | 1.00 |
| IPS1 - IPS3 | Time | -0.04 | 0.02 | 372.00 | -1.58 | 1.00 |
| IPS1 - MT | Time | -0.01 | 0.02 | 372.00 | -0.29 | 1.00 |
| IPS1 - (V1d-V2d) | Time | -0.03 | 0.02 | 372.00 | -1.23 | 1.00 |
| IPS1 - V3AB | Time | -0.05 | 0.02 | 372.00 | -1.96 | 1.00 |
| IPS2 - IPS3 | Time | -0.01 | 0.02 | 372.00 | -0.47 | 1.00 |
| IPS2 - MT | Time | 0.02 | 0.02 | 372.00 | 0.82 | 1.00 |
| IPS2 - (V1d-V2d) | Time | -0.00 | 0.02 | 372.00 | -0.12 | 1.00 |
| IPS2 - V3AB | Time | -0.02 | 0.02 | 372.00 | -0.85 | 1.00 |
| IPS3 - MT | Time | 0.03 | 0.02 | 372.00 | 1.28 | 1.00 |
| IPS3 - (V1d-V2d) | Time | 0.01 | 0.02 | 372.00 | 0.35 | 1.00 |
| IPS3 - V3AB | Time | -0.01 | 0.02 | 372.00 | -0.38 | 1.00 |
| MT - (V1d-V2d) | Time | -0.02 | 0.02 | 372.00 | -0.93 | 1.00 |
| MT - V3AB | Time | -0.04 | 0.02 | 372.00 | -1.66 | 1.00 |
| (V1d-V2d) - V3AB | Time | -0.02 | 0.02 | 372.00 | -0.73 | 1.00 |

Supplementary table 27: Nuggets - ROIs contrasts for each map type (Bonferroni corrected) with Kenward-Roger's method for degrees of freedom

| | Sum.Sq | Mean.Sq | NumDF | DenDF | F.value | Pr(>F) |
|---|---|---|---|---|---|---|
| ROI | 0.31 | 0.04 | 7 | 372.00 | 5.09 | 0.00 |
| MapType | 0.61 | 0.61 | 1 | 372.00 | 69.64 | 0.00 |
| Hemi | 0.00 | 0.00 | 1 | 372.00 | 0.00 | 0.99 |
| ROI:MapType | 0.08 | 0.01 | 7 | 372.00 | 1.36 | 0.22 |
| ROI:Hemi | 0.10 | 0.01 | 7 | 372.00 | 1.58 | 0.14 |
| MapType:Hemi | 0.00 | 0.00 | 1 | 372.00 | 0.35 | 0.56 |
| ROI:MapType:Hemi | 0.08 | 0.01 | 7 | 372.00 | 1.33 | 0.23 |

Supplementary table 28: Ranges - type III ANOVA on LME model estimates with Satterthwaite's method for degrees of freedom

| | estimate | SE | df | t.ratio | p.value |
|---|---|---|---|---|---|
| FEFs - IPS0 | -0.02 | 0.02 | 372.00 | -0.87 | 1.00 |
| FEFs - IPS1 | 0.01 | 0.02 | 372.00 | 0.65 | 1.00 |
| FEFs - IPS2 | 0.01 | 0.02 | 372.00 | 0.56 | 1.00 |
| FEFs - IPS3 | -0.01 | 0.02 | 372.00 | -0.29 | 1.00 |
| FEFs - MT | -0.01 | 0.02 | 372.00 | -0.58 | 1.00 |
| FEFs - (V1d-V2d) | -0.08 | 0.02 | 372.00 | -4.35 | 0.00 |
| FEFs - V3AB | -0.00 | 0.02 | 372.00 | -0.06 | 1.00 |
| IPS0 - IPS1 | 0.03 | 0.02 | 372.00 | 1.53 | 1.00 |
| IPS0 - IPS2 | 0.03 | 0.02 | 372.00 | 1.43 | 1.00 |
| IPS0 - IPS3 | 0.01 | 0.02 | 372.00 | 0.58 | 1.00 |
| IPS0 - MT | 0.01 | 0.02 | 372.00 | 0.29 | 1.00 |
| IPS0 - (V1d-V2d) | -0.06 | 0.02 | 372.00 | -3.48 | 0.02 |
| IPS0 - V3AB | 0.01 | 0.02 | 372.00 | 0.82 | 1.00 |
| IPS1 - IPS2 | -0.00 | 0.02 | 372.00 | -0.10 | 1.00 |
| IPS1 - IPS3 | -0.02 | 0.02 | 372.00 | -0.94 | 1.00 |
| IPS1 - MT | -0.02 | 0.02 | 372.00 | -1.24 | 1.00 |
| IPS1 - (V1d-V2d) | -0.09 | 0.02 | 372.00 | -5.01 | 0.00 |
| IPS1 - V3AB | -0.01 | 0.02 | 372.00 | -0.71 | 1.00 |
| IPS2 - IPS3 | -0.02 | 0.02 | 372.00 | -0.84 | 1.00 |
| IPS2 - MT | -0.02 | 0.02 | 372.00 | -1.14 | 1.00 |
| IPS2 - (V1d-V2d) | -0.09 | 0.02 | 372.00 | -4.91 | 0.00 |
| IPS2 - V3AB | -0.01 | 0.02 | 372.00 | -0.61 | 1.00 |
| IPS3 - MT | -0.01 | 0.02 | 372.00 | -0.30 | 1.00 |
| IPS3 - (V1d-V2d) | -0.07 | 0.02 | 372.00 | -4.06 | 0.00 |
| IPS3 - V3AB | 0.00 | 0.02 | 372.00 | 0.23 | 1.00 |
| MT - (V1d-V2d) | -0.07 | 0.02 | 372.00 | -3.77 | 0.01 |
| MT - V3AB | 0.01 | 0.02 | 372.00 | 0.53 | 1.00 |
| (V1d-V2d) - V3AB | 0.08 | 0.02 | 372.00 | 4.30 | 0.00 |

Supplementary table 29: Ranges - ROIs contrasts (Bonferroni corrected) with Kenward-Roger's method for degrees of freedom

| | estimate | SE | df | t.ratio | p.value |
|---|---|---|---|---|---|
| Space - Time | 0.08 | 0.01 | 372.00 | 8.35 | 0.00 |

Supplementary table 30: Ranges - map types contrast (Bonferroni corrected) with Kenward-Roger's method for degrees of freedom

### Supplementary Methods - MRI data processing

Results included in this manuscript come from preprocessing performed using *fMRIPrep* 21.0.2 ([1]; [2]; RRID:SCR_016216), which is based on *Nipype* 1.6.1 ([3]; [4]; RRID:SCR_002502).

#### Preprocessing of B0 inhomogeneity mappings

A total of 12 fieldmaps were found available within the input BIDS structure for this particular subject. A *B0*-nonuniformity map (or *fieldmap*) was estimated based on two (or more) echo-planar imaging (EPI) references with `topup` ([5]; FSL 6.0.5.1:57b01774).

#### Anatomical data preprocessing

A total of 1 T1-weighted (T1w) images were found within the input BIDS dataset. The T1-weighted (T1w) image was corrected for intensity non-uniformity (INU) with `N4BiasFieldCorrection` [6], distributed with ANTs 2.3.3 [7, RRID:SCR_004757], and used as T1w-reference throughout the workflow. The T1w-reference was then skull-stripped with a *Nipype* implementation of the `antsBrainExtraction.sh` workflow (from ANTs), using OASIS30ANTs as target template. Brain tissue segmentation of cerebrospinal fluid (CSF), white-matter (WM) and gray-matter (GM) was performed on the brain-extracted T1w using `fast` [FSL 6.0.5.1:57b01774, RRID:SCR_002823, 8]. Brain surfaces were reconstructed using `recon-all` [FreeSurfer 6.0.1, RRID:SCR_001847, 9], and the brain mask estimated previously was refined with a custom variation of the method to reconcile ANTs-derived and FreeSurfer-derived segmentations of the cortical gray-matter of Mindboggle [RRID:SCR_002438, 10]. Volume-based spatial normalization to two standard spaces (MNI152NLin6Asym, MNI152NLin2009cAsym) was performed through nonlinear registration with `antsRegistration` (ANTs 2.3.3), using brain-extracted versions of both T1w reference and the T1w template. The following templates were selected for spatial normalization: *FSL's MNI ICBM 152 non-linear 6th Generation Asymmetric Average Brain Stereotaxic Registration Model*[[11], RRID:SCR_002823; TemplateFlow ID: MNI152NLin6Asym], *ICBM 152 Nonlinear Asymmetrical template version 2009c*[[12], RRID:SCR_008796; TemplateFlow ID: MNI152NLin2009cAsym].

#### Functional data preprocessing

For each of the 12 BOLD runs found per subject (across all tasks and sessions), the following preprocessing was performed. First, a reference volume and its skull-stripped version were generated using a custom methodology of *fMRIPrep*. Head-motion parameters with respect to the BOLD reference (transformation matrices, and six corresponding rotation and translation parameters) are estimated before any spatiotemporal filtering using `mcflirt` [FSL 6.0.5.1:57b01774, 13]. The estimated *fieldmap* was then aligned with rigid-registration to the target EPI (echo-planar imaging) reference run. The field coefficients were mapped on to the reference EPI using the transform. The BOLD reference was then co-registered to the T1w reference using `bbregister` (FreeSurfer) which implements boundary-based registration [14]. Co-registration was configured with six degrees of freedom. Several confounding time-series were calculated based on the *preprocessed BOLD*: framewise displacement (FD), DVARS and three region-wise global signals. FD was computed using two formulations following Power (absolute sum of relative motions, [15]) and Jenkinson (relative root mean square displacement between affines, [13]). FD and DVARS are calculated for each functional run, both using their implementations in *Nipype* [following the definitions by 15]. The three global signals are extracted within the CSF, the WM, and the whole-brain masks. Additionally, a set of physiological regressors were extracted to allow for component-based noise correction [*CompCor*, 16]. Principal components are estimated after high-pass filtering the *preprocessed BOLD* time-series (using a discrete cosine filter with 128s cut-off) for the two *CompCor* variants: temporal (tCompCor) and anatomical (aCompCor). tCompCor components are then calculated from the top 2% variable voxels within the brain mask. For aCompCor, three probabilistic masks (CSF, WM and combined CSF+WM) are generated in anatomical space. The implementation differs from that of Behzadi et al. in that instead of eroding the masks by 2 pixels on BOLD space, the aCompCor masks are subtracted a mask of pixels that likely contain a volume fraction of GM. This mask is obtained by dilating a GM mask extracted from the FreeSurfer's *aseg* segmentation, and it ensures components are not extracted from voxels containing a minimal fraction of GM. Finally, these masks are resampled into BOLD space and binarized by thresholding at 0.99 (as in the original implementation). Components are also calculated separately within the WM and CSF masks. For each CompCor decomposition, the $k$ components with the largest singular values are retained, such that the retained components' time series are sufficient to explain 50 percent of variance across the nuisance mask (CSF, WM, combined, or temporal). The remaining components are dropped from consideration. The head-motion estimates calculated in the correction step were also placed within the corresponding confounds file. The confound time series derived from head motion estimates and global signals were expanded with the inclusion of temporal derivatives and quadratic terms for each [17]. Frames that exceeded a threshold of 0.5 mm FD or 1.5 standardised DVARS were annotated as motion outliers. The BOLD time-series were resampled into standard space, generating a *preprocessed BOLD run in MNI152NLin6Asym space*. First, a reference volume and its skull-stripped version were generated using a custom methodology of *fMRIPrep*. The BOLD time-series were resampled onto the following surfaces (FreeSurfer reconstruction nomenclature): *fsnative*, *fsaverage*. Automatic removal of motion artifacts using independent component analysis [ICA-AROMA, 18] was performed on the *preprocessed BOLD on MNI space* time-series after removal of non-steady state volumes and spatial smoothing with an isotropic, Gaussian kernel of 6mm FWHM (full-width half-maximum). Corresponding "non-aggresively" denoised runs were produced after such smoothing. Additionally, the "aggressive" noise-regressors were collected and placed in the corresponding confounds file. All resamplings can be performed with *a single interpolation step* by composing all the pertinent transformations (i.e. head-motion transform matrices, susceptibility distortion correction when available, and co-registrations to anatomical and output spaces). Gridded (volumetric) resamplings were performed using `antsApplyTransforms` (ANTs), configured with Lanczos interpolation to minimize the smoothing effects of other kernels [19]. Non-gridded (surface) resamplings were performed using `mri_vol2surf` (FreeSurfer).

Many internal operations of *fMRIPrep* use *Nilearn* 0.8.1 [20, RRID:SCR_001362], mostly within the functional processing workflow. For more details of the pipeline, see the section corresponding to workflows in *fMRIPrep*'s documentation.

**Copyright Waiver**

The above boilerplate text was automatically generated by fMRIPrep with the express intention that users should copy and paste this text into their manuscripts *unchanged*. It is released under the CC0 license.

[13] M. Jenkinson, P. Bannister, M. Brady, and S. Smith, "Improved optimization for the robust and accurate linear registration and motion correction of brain images," *NeuroImage*, vol. 17, no. 2, pp. 825–

841, 2002, ISSN: 1053-8119. DOI: 10.1006/nimg.2002.1132. [Online]. Available: http://www.sciencedirect.com/science/article/pii/S1053811902911328.
